# Clair3-RNA: A deep learning-based small variant caller for long-read RNA sequencing data

**DOI:** 10.1101/2024.11.17.624050

**Authors:** Zhenxian Zheng, Xian Yu, Lei Chen, Yan-Lam Lee, Cheng Xin, Angel On Ki Wong, Miten Jain, Rupesh K Kesharwani, Fritz J Sedlazeck, Ruibang Luo

## Abstract

Variant calling using long-read RNA sequencing (lrRNA-seq) can be applied to diverse tasks, such as capturing full-length isoforms and gene expression profiling. It poses challenges, however, due to higher error rates than DNA data, the complexities of transcript diversity, RNA editing events, etc. In this paper, we propose Clair3-RNA, the first deep learning-based variant caller tailored for lrRNA-seq data. Clair3-RNA leverages the strengths of the Clair series’ pipelines and incorporates several techniques optimized for lrRNA-seq data, such as uneven coverage normalization, refinement of training materials, editing site discovery, and the incorporation of phasing haplotype to enhance variant-calling performance. Clair3-RNA is available for various platforms, including PacBio and ONT complementary DNA sequencing (cDNA), and ONT direct RNA sequencing (dRNA). Our results demonstrated that Clair3-RNA achieved a ∼91% SNP F1-score on the ONT platform using the latest ONT SQK-RNA004 kit (dRNA004) and a ∼92% SNP F1-score in PacBio Iso-Seq and MAS-Seq for variants supported by at least four reads. The performance reached a ∼95% and ∼96% F1-score for ONT and PacBio, respectively, with at least ten supporting reads and disregarding the zygosity. With read phased, the performance reached ∼97% for ONT and ∼98% for PacBio. Extensive evaluation of various GIAB samples demonstrated that Clair3-RNA consistently outperformed existing callers and is capable of distinguishing RNA high-quality editing sites from variants accurately. Clair3-RNA is open-source and available at (https://github.com/HKU-BAL/Clair3-RNA).

## Introduction

RNA sequencing (RNA-seq) is extensively utilized for gene expression quantification, isoform identification and quantification, and gene fusion analysis [1, 2]. Recent advancements highlight the potential of variant calling [3–6]in RNA-seq, showcasing its numerous advantages for this application. First, it is more cost-effective than DNA sequencing (DNA-seq). Second, the presence of multiple isoforms of some genes increases sequencing depth, enhancing the efficacy of variant detection. Third, RNA-seq allows the identification of post-transcriptional modifications [7], which are not detectable using DNA-seq.

Long-read RNA sequencing (lrRNA-seq) has become popular for its capability to capture full-length isoforms [8] and novel transcript discovery [9]. Pacific Biosciences (PacBio) and Oxford Nanopore Technology (ONT) are two leading platforms in this field. PacBio introduced Iso-Seq for full-length cDNA sequencing using PacBio SMRT sequencing technology [10]. PacBio also provides the advanced Multiplexed Arrays Sequencing (MAS-Seq) method, which increases throughput by combining cDNA molecules into longer concatenated fragments. ONT also introduced direct RNA (dRNA) sequencing, which is a nanopore-based technique that directly sequences RNA without enzymatic conversions [11]. ONT dRNA sequencing enables the observation of full-length transcripts, RNA modifications, and RNA epigenetics analysis [12], which are challenging to detect using cDNA sequencing. Recently, deep-learning methods have improved the capabilities of germline and somatic DNA variant calling [13–15] and short-read RNA variant calling [3]. However, there exists no deep learning-based caller designed specifically for variant calling in lrRNA-seq yet.

RNA-seq offers advantages for variant calling and interpretation, such as that the identified allele is guaranteed to be expressed in contrast to DNA variant calling. Nevertheless, several disadvantages need to be considered such as a higher error rate than DNA-seq, which averages a 1-5% error rate. This necessitates robust variant-calling systems for distinguishing true variants from sequencing artifacts. Second, unlike the uniform read coverage typically observed in DNA-seq data [4], the coverage is uneven across genomic regions in RNA-seq, and the variable coverage poses challenges for accurate variant calling, particularly in regions characterized by inadequate read support or excessive read coverage that is not accounted for in neural network training. Additionally, RNA-seq faces RNA editing events, especially A-to-I editing catalyzed by adenosine deaminases acting on RNA (ADAR) [16], leading to A-to-G or T-to-C substitutions that can resemble genuine variants, thus introducing false positives in RNA-seq. Moreover, while phasing has been demonstrated to be advantageous for lrDNA-seq analyses [13, 14], no studies have yet investigated the impact of phasing on variant calling performance in lrRNA-seq, particularly when reads are shorter and zygosity switching can happen.

In this study, we introduce Clair3-RNA, the first deep learning-based small-variant caller designed for long-read RNA sequencing data. Clair3-RNA utilizes a Bi-LSTM pileup model with multi-task output for variant prediction. The model was trained on lrRNA-seq datasets using various techniques, such as coverage normalization, variant zygosity switching, training sample augmentation, and RNA editing-site tagging to optimize variant-calling accuracy. We further developed a workflow that incorporates additional phasing information into the neural network, thereby enhancing variant calling performance and demonstrating the efficacy of phasing for lrRNA-seq. We also designed a benchmarking pipeline that defines callable regions and benchmarking metrics, enabling a fair evaluation of variant-calling performance. Our benchmarking results demonstrate that Clair3-RNA outperformed other variant callers and can achieve ∼93%, ∼93%, and ∼91% F1-score in PacBio Iso-Seq, PacBio MAS-Seq, and the ONT dRNA004 platform, respectively, for variants with at least four supporting reads. With the incorporation of additional pileup phasing features, Clair3-RNA reached 98.59% SNP F1-score for PacBio MAS-Seq, 98.24% for Iso-Seq, and 97.16% for ONT dRNA004 when excluding zygosity and requiring a minimum of 10 supporting reads. Furthermore, in our analysis of GENCODE-annotated genes, we identified 1,085 and 507 genes that could be completely phased, as well as 1,165 and 655 genes without SNP, Indel or switch errors in the PacBio MAS-Seq HG004 and ONT dRNA004 HG004 datasets, respectively. Furthermore, we established a high-quality benchmark set of RNA editing sites by cross-evaluating RNA samples with the corresponding DNA samples. Clair3-RNA also demonstrated outperforming runtime and memory usage compared to other variant callers, requiring ∼20 minutes for PacBio Iso-Seq and ∼30 minutes for ONT dRNA004 variant calling using 16 CPU cores, with a memory footprint of about 7 GB.

## Results

### Overview of Clair3-RNA

Clair3-RNA is a multi-task bidirectional long short-term memory (Bi-LSTM) neural network designed for lrRNA-seq variant calling. Clair3-RNA takes a BAM/CRAM file as input to identify candidate variants with any minor allelic fraction exceeding a specific threshold (typically 0.08, configurable by option) and then computes a summarized pileup tensor of the candidate and employs a neural network for prediction. Clair3-RNA includes two probabilistic output tasks - 21-genotype and zygosity - facilitating the prediction of variants and artifacts. The **Methods** section details techniques for training data augmentation and optimizations specific to lrRNA-seq, addressing challenges like uneven coverage, variant zygosity inconsistency, and RNA editing events. Clair3-RNA was trained with the Genome in a Bottle (GIAB) HG002 sample and supports variant calling on multiple PacBio and ONT instruments and chemistry configurations.

In this section, we assess the performance of Clair3-RNA on PacBio cDNA, ONT cDNA, and ONT dRNA datasets. Our evaluation involved comparing results with four other variant callers, including LongcallR [17] (a long-read SNP caller for RNA data), Clair3 [13] (our deep learning-based germline variant caller for DNA data), and DeepVariant [14] (the first deep learning-based variant caller, developed by Google). For Clair3-RNA and LongcallR, we used the corresponding specific RNA model and parameter settings for RNA variant calling. Although DeepVariant supports RNA variant calling [3], it is optimized for the Illumina platform, not for lrRNA-seq. Clair3 and the other tools in the series [15, 18–21] before Clair3-RNA were used for germline and somatic small variant calling, but none of them were designed for lrRNA-seq. The performance of Clair3 and DeepVariant on lrRNA-seq is not optimized and is provided solely for reference purposes. Therefore, for performance comparison on long-read data, the best available DNA models of Clair3 and DeepVariant were employed for benchmarking. We used two GIAB samples, HG004 and HG005, for benchmarking. Since LongcallR is limited to SNP variant detection, the comparisons with Clair3-RNA were conducted on SNP performance only. For other methods, we benchmarked both SNP and Indel performance.

### Generating callable high-confidence benchmarking regions for evaluation

RNA transcripts represent only subsets of the corresponding DNA sequence. Certain regions may have insufficient coverage, resulting in undetectable variants in those regions. To address this, we identified callable high-confidence benchmarking regions by intersecting callable regions with adequate read coverage, with those high-confidence regions defined by GIAB. We also considered the zygosity imbalance in the RNA data. In certain cases, a heterozygous variant in DNA could be either a homozygous variant or a homozygous reference in RNA. This discrepancy arises from the dominant expression of one of the haplotypes. By disregarding zygosity, we aim to understand the impact of zygosity during evaluation better. More details are listed in the **Methods** - **Benchmarking metrics** section.

We further introduced two key benchmarking factors for performance assessment: read coverage (DP) and allele depth (AD). DP signifies the read coverage of the candidate site meeting minimum mapping quality support and excluding all supplementary and secondary alignments. AD represents the read coverage that supports the true alternate allele at the site. We also considered RNA editing events and created a reliable benchmark set of editing sites for comprehensive evaluation. More details are provided in the **Methods** - **Orthogonal high-quality RNA editing benchmark set generation** section.

### Performance on ONT

The performance on the ONT platform was carried out in six datasets. Details of the benchmarking datasets are provided in **Supplementary Table 1**. These ONT datasets include ONT cDNA sequencing, dRNA sequencing using the SQK-RNA002 kit (hereafter referred to as dRNA002), and dRNA sequencing employing the ONT latest SQK-RNA004 kit (hereafter dRNA004). The ONT cDNA and dRNA002 data were made available by GIAB. The ONT cDNA data was sequenced using R9.4.1 chemistry and basecalled using Guppy [22] version 6.4.6. The ONT dRNA002 data was sequenced using the dRNA SQK-RNA002 sequencing kit and basecalled with Guppy 6.4.6. In 2023, ONT released the dRNA SQK-RNA004 kit [23] with an enhanced throughput and a reduced error rates compared to the SQK-RNA002 kit. We sequenced GIAB HG002, HG004, and HG005 using the dRNA SQK-RNA004 kit at the University of Hong Kong. The total RNA of three samples was acquired from the Coriell Institute, sequenced using the SQK-RNA004 kit, and basecalled using the Dorado basecaller [24] version 0.7.2. All basecalled alignments were mapped to the GRCh38 reference genome using the minimap2 aligner in splicing alignment mode. More details can be found in the **Methods - ONT SQK-RNA004 kit library preparation and sequencing** section. As shown in **Figure 2a**, the overall throughput increased from an average of 3 GB to 18 GB per flowcell using dRNA004 compared with dRNA002 and the average error rate decreased from 9.7% to 1.8%.

**Figure 1.**
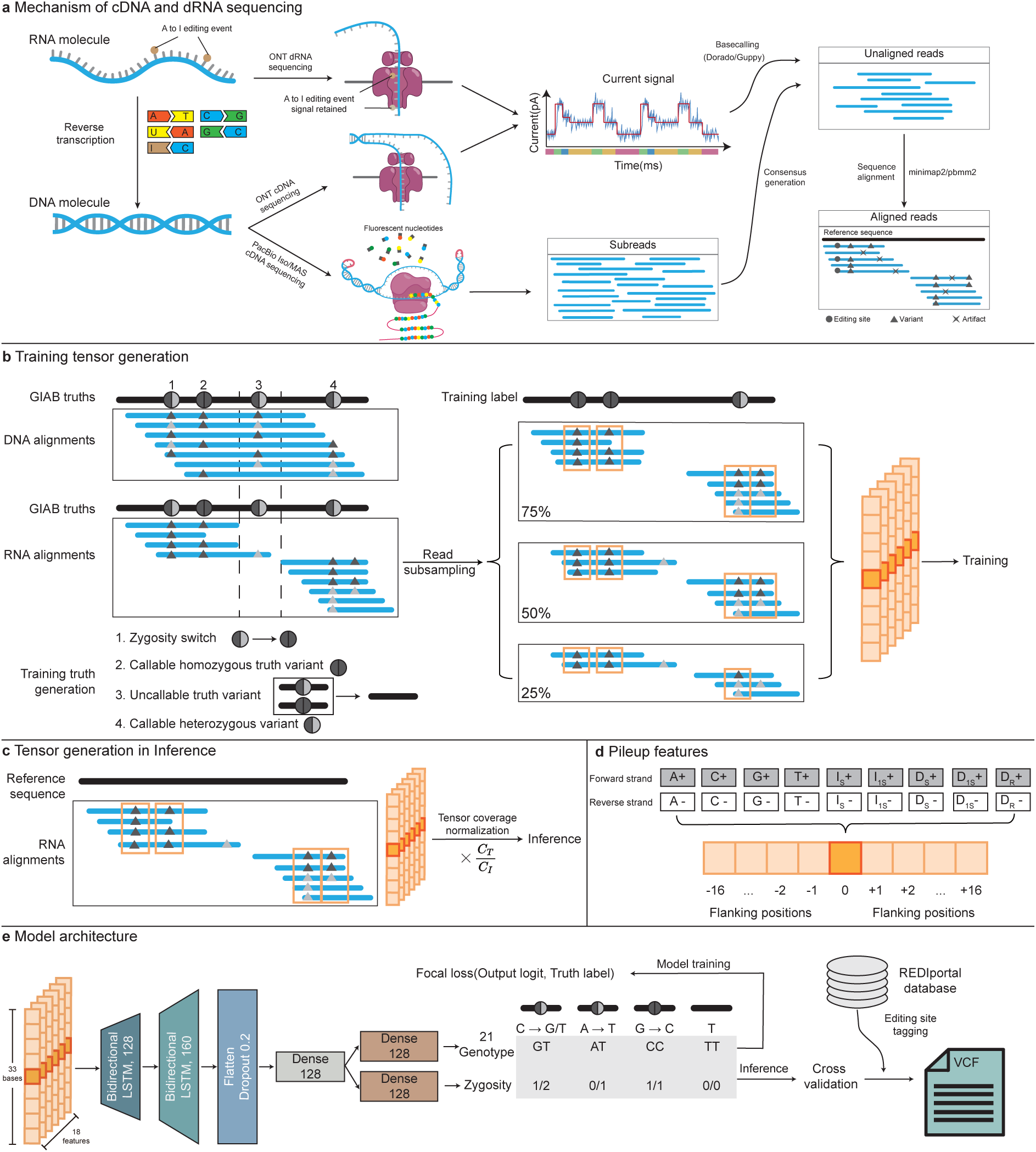
Overview of Clair3-RNA variant-calling workflow. (a) The figure contrasts RNA sequencing techniques, specifically highlighting the mechanisms of direct RNA sequencing (dRNA) and complementary DNA (cDNA) sequencing with reverse transcription. In dRNA sequencing, inosine editing is sequenced as is, while in cDNA sequencing, it is converted to Guanine nucleotide. (b) The illustration of training tensor generation outlines the process of deriving training tensors from DNA and RNA alignments, incorporating labels obtained from GIAB truths. Variants located outside the callable region are omitted, and the zygosity switch is applied, whereby heterozygous variants displaying allelic fractions indicative of homozygous variant or homozygous reference in RNA are reclassified. Read subsampling is employed for data augmentation, and pileup tensors are produced for every candidate site, along with flanking positions. (c) The illustration of tensor generation in inference shows the process of creating tensors in inference. Coverage normalization is adopted for exceeding coverage in inference. (d) The figure illustrates 18 pileup features for each position in the forward and reverse strands. More details are provided in the **Supplementary methods − Description of RNA pileup input features** section. (e) In this illustration of the Clair3-RNA model architecture, the pileup network includes a two-layer LSTMs with two dense layers. The output supports two probabilistic tasks for classification: 21-genotype and zygosity. Variants identified by the model are tagged by the REDIportal database and recorded in VCF files.

**Figure 2.**
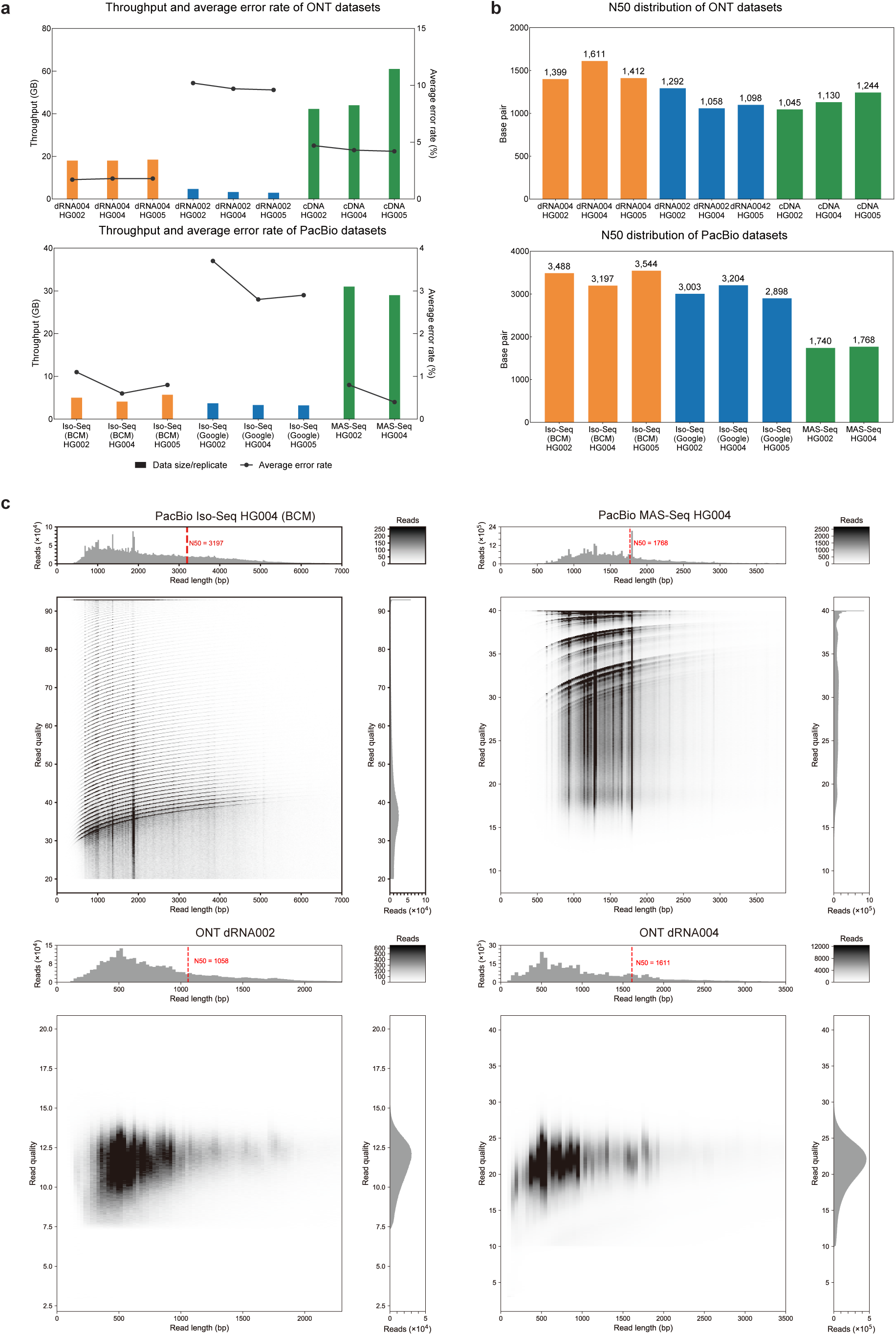
Dataset distribution of different platforms. (a) The average sequencing throughput and sequencing error rate for all sequencing replicated across different datasets are illustrated. (b) N50distributions are compared among different datasets. (c) The read length and read quality distributions of the HG004 sample are depicted across four platforms: PacBio Iso-Seq, PacBio MAS-Seq, ONT dRNA002, and ONT dRNA004.

As shown in **Figure 3b** and **Supplementary Table 2**, Clair3-RNA consistently outperformed other variant callers in the ONT dRNA004 and cDNA datasets for both the HG004 and HG005 samples. Specifically, Clair3-RNA achieved 91.00% and 91.73% F1-score in HG004 and HG005, respectively, using dRNA004 sequencing when DP≥4 and AD≥2. In comparison, LongcallR had HG004/61.07% and HG005/65.20% F1-score, respectively; Clair3 had HG004/74.85% and HG005/77.21%, and DeepVariant had HG004/57.94% and HG005/56.96% F1-score. LongcallR employs a Bayesian statistical model for variant calling, demanding higher sequencing coverage for accuracy, as suggested by the authors [17]. When we increased the minimal requirement of DP to 10 and AD to 4, we observed that LongcallR performance improved; the F1-score reached HG004/72.42% and HG005/74.42%, resulting in an average ∼10% improvement against the default setting. Using the same setting, while the performance gap narrowed, Clair3-RNA still performed better than longcallR, reaching HG004/94.75% and HG005/95.17%, respectively.

**Figure 3.**
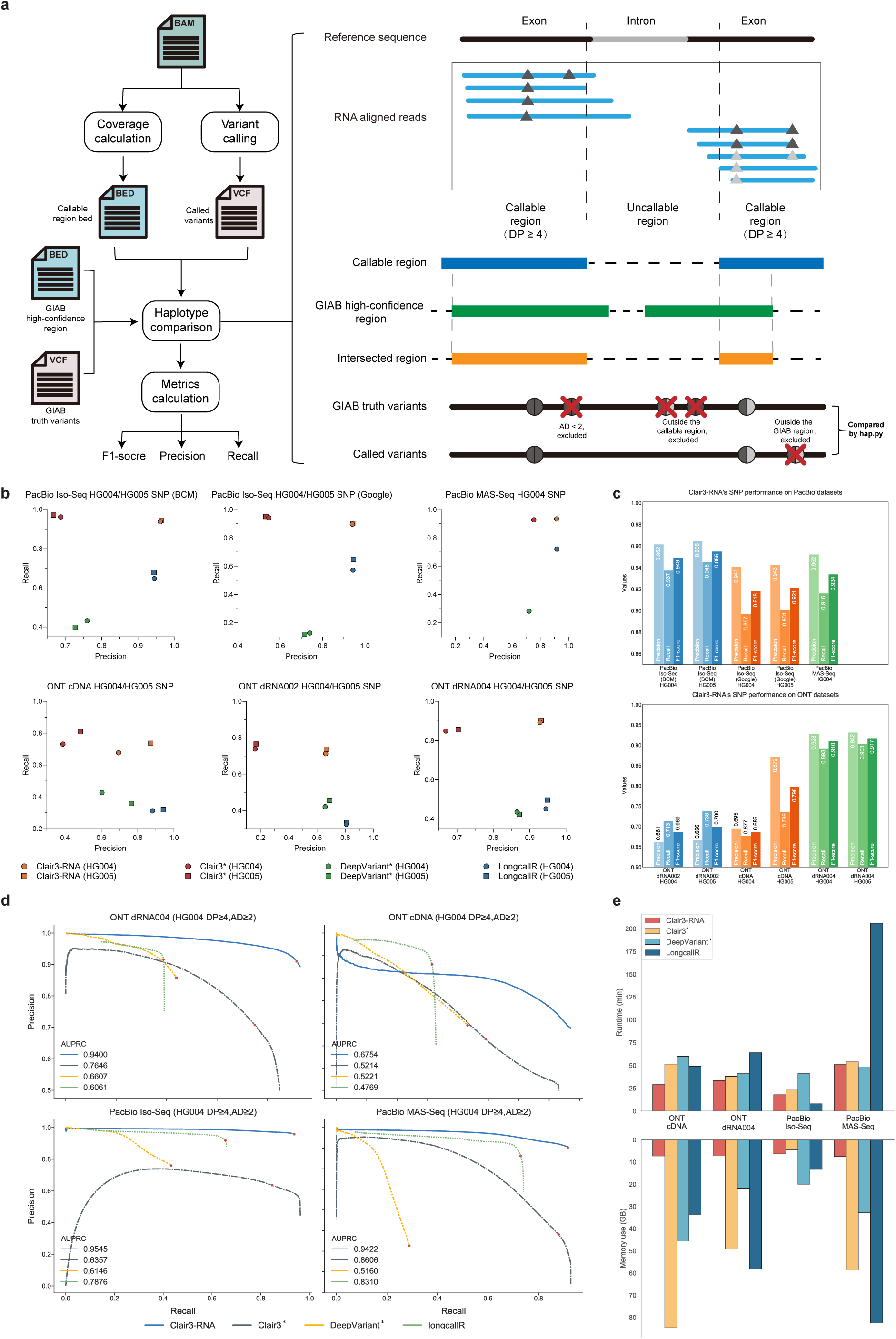
PacBio and ONT benchmarking results. (a) The RNA variant benchmarking workflow is depicted, starting with alignments utilized for coverage calculation to determine the callable region. The callable region was then refined by intersecting it with GIAB high-confidence regions. Variants with coverage depth (DP) of less than 4 or allele depth (AD) of less than 2 were excluded from benchmarking. Evaluation metrics such as precision, recall, and F1-score for SNP and Indel were obtained by comparing called variants with truth variants using hap.py. (b) This shows the precision and recall performance on different sequencing datasets. The star superscript indicates that Clair3 and DeepVariant are two DNA variant callers. (c) This shows the SNP performance of Clair3-RNA on PacBio and ONT datasets. (d) These are the precision-recall curves of four variant callers on different datasets; the dot indicates the best F1-score. AUPRC is the area under the precision-recall curve. (e) This shows the runtime and memory usage of four variant callers on different datasets.

In the cDNA dataset, Clair3-RNA yielded HG004/68.60% and HG005/79.83% F1-score and achieved better performance than longcallR (HG004/+23.72%, HG005/+23.72%), Clair3 (HG004/+17.85%, HG005/+19.19%) and DeepVariant (HG004/+18.60%, HG005/+30.99%). The lower performance across all callers might be because of a higher base error rate of the cDNA dataset generated with R9.4.1 chemistry. We expect the performance to improve notably with cDNA sequencing using R10 chemistry.

### Boosting the performance across ONT RNA-specific flow cells

In our evaluation, we compared the performance of different callers in the GIAB HG004 and HG005 dRNA002 and dRNA004 datasets. As illustrated in **Figure 2b and Figure 2c**, the data throughput increased ∼six times per flowcell, from ∼3GB in dRNA002 to ∼18GB in dRNA004. The read N50 also increased from ∼1100bp in dRNA002 to ∼1500bp in dRNA004. Across all callers, we observed improvements in precision, recall and F1-score, signifying improved read accuracy in the dRNA004 data. Notably, Clair3-RNA exhibited an average F1-score improvement of 22.07% (rising from 68.61% to 91.00% in HG004 and from 69.98% to 91.73% in HG005), and LongcallR improvement by 16.35%, Clair3 by 48.69%, and DeepVariant by 4.36%.

Among the three ONT benchmarking datasets, we noted that Clair3 had lower precision and DeepVariant had lower recall. In the dRNA004 dataset, Clair3 had only HG004/66.93% and HG005/70.33% for precision, but HG004/84.89% and HG005/85.58% for recall. DeepVariant had HG004 /86.53% and HG005/87.14% for precision, but almost half the recall with HG004/43.55% and HG005/42.31%. LongcallR also had better precision than its recall, with HG004/94.49% and HG005/94.89% precision, but HG004/45.11% and HG005/49.67% recall. Compared with other callers, Clair3-RNA demonstrated balanced precision and recall, underscoring the necessity of a variant caller specific for lrRNA-seq.

### Performance on PacBio

We benchmarked the performance on the PacBio platform using two sequencing technologies, Iso-Seq and MAS-Seq, both developed for PacBio cDNA sequencing. We collected the HG004 and HG005 Iso-Seq RNA sequencing datasets from two sources, Baylor College of Medicine (BCM) and Google, and the HG004 MAS-Seq RNA sequencing dataset from BCM. More details of sequencing data are provided in **Supplementary Table 1**. Compared with ONT sequencing, PacBio MAS-Seq and Iso-Seq have a higher N50 read length, with ∼3000bp for PacBio Iso-Seq and ∼1700bp for MAS-Seq, versus ∼1200bp for ONT sequencing.

As indicated in **Figure 3** and **Supplementary Table 3**, Clair3-RNA consistently achieved a SNP F1-score exceeding 91% across all Iso-Seq and MAS-Seq datasets for HG004 and HG005, showcasing better performance than the second-best LongcallR, with an F1-score ranging from 71% to 80%. Clair3-RNA also demonstrated consistent performance across all PacBio sequencing sources, achieving HG004/94.93% and HG005/95.50% F1-score for BCM Iso-Seq, HG004/91.84%, an HG005/92.13% F1-score for Google Iso-Seq, and an HG004/93.38% F1-score for BCM MAS-Seq. In terms of precision and recall balance, Clair3-RNA exhibited a balanced performance with 95.25% precision and 91.93% recall. Notably, Clair3 maintained an average recall rate of 95.08%, but had a low precision (63.76%). Conversely, LongcallR had 93.94% precision but 65.31% recall and DeepVariant showed lower precision (73.25%) and recall (27.15%).

The PacBio datasets were originally aligned with the GRCh38 reference genome using pbmm2, a minimap2 aligner wrapper specifically designed for the PacBio platform. During our experimental analysis focusing on alignment methods, we observed that the pbmm2 aligner tended to introduce more false positives and false negatives compared to the minimap2 aligner when using Clair3-RNA for small variant calling on PacBio. We aligned five datasets using pbmm2 and minimap2 splice mode to assess the impact of different alignment strategies. Further details regarding the alignment procedure are included in the **Supplementary Notes** - **Command line used** section.

As shown in **Figure 8** and **Supplementary Table 5**, when utilizing minimap2 in place of pbmm2, Clair3-RNA’s performance improved across all datasets. The average number of false positives and false negatives decreased by 25.42% and 0.79%, respectively. The averaged F1-score of Clair3-RNA improved from 94.57% to 95.10% (+0.53%) for BCM Iso-Seq, from 91.37% to 91.98% (+0.61%) for Google Iso-Seq, and from 92.76% to 93.38% (+0.62%) for BCM MAS-Seq. Similar trends were observed in other callers; longcallR, Clair3, and DeepVariant demonstrated average F1-score improvements of 0.46%, 14.40%, and 2.17%, respectively. These improvements suggest that using minimap2 had better performance compared to using pbmm2 when using the default splice mode setting in PacBio.

### Sequencing coverage and allele depth analysis

Insufficient coverage and inadequate read support for alternate bases may lead to false negatives in evaluations. **Figure 4a** shows the benchmarking performance across different DP and AD thresholds.

**Figure 4.**
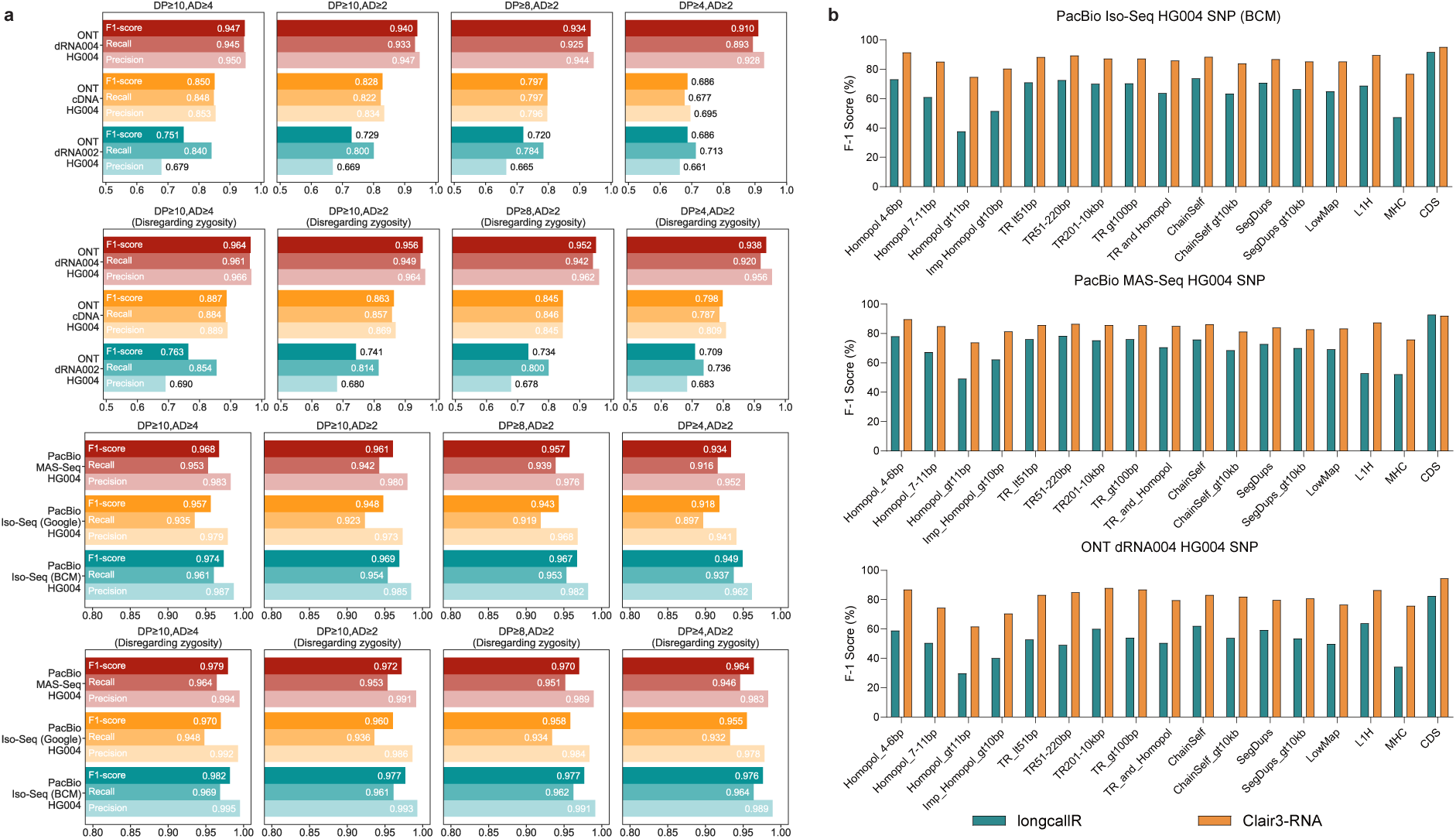
Performance evaluation under different settings. (a) This shows Clair3-RNA’s SNP performance on different datasets for different coverage depth (DP) and allele depth (AD) settings. The performance of “Disregarding zygosity” was determined by enabling the “--skip_genotyping” option when using the “get_overall_metrics” for metrics calculation. (b) This shows the SNP performance comparison between Clair3-RNA and LongcallR in different genomic contexts. The figure was created using Genome in a Bottle stratification v3 and hap.py. Detailed descriptions of each sub-category are provided in the **Supplementary Table 6**.

In the ONT dRNA004 HG004 dataset, increasing the depth from 4 to 8 resulted in a 1.54% increase in precision and a 2.67% improvement in recall. Raising the depth to 10 showed marginal gains of +0.33% in precision and +0.58% in recall, highlighting the difficulty in capturing low-coverage variants. Similar trends were observed in the PacBio datasets. In the BCM Iso-Seq HG004 dataset, raising the depth benchmark from 4 to 8 led to precision and recall improvements of 2.15% and 1.90%, respectively. Increasing to a depth of 10 showed modest gains of 0.38% in precision and 0.18% in recall. AD is another crucial factor for variant evaluation. When the AD was increased from 2 to 4, the ONT dRNA004 HG004 dataset showed a 0.28% improvement in precision and a 1.91% increase in recall. Similarly, in the BCM Iso-Seq HG004 dataset, there was an improvement of 0.51% in precision and 1.51% in recall. This highlights the importance of allele depth in accurately calling missing variants.

### Zygosity switch analysis in RNA dataset

Zygosity switch is also common in RNA data due to factors like differential gene expression and RNA splicing [3]. To evaluate this, we used the Clair3-RNA’s "calculate_overall_metrics" function to consider the zygosity switch by disregarding zygosity in benchmarking. As shown in **Figure 4a**, **Supplementary Table 2, and Supplementary Table 3**, disregarding zygosity led to an average 2.43% and 2.88% improvement in the F1-score on the ONT and PacBio, respectively. Benchmarking on the ONT dRNA004 HG004 dataset, disregarding zygosity resulted in a reduction of 2,699 FPs and 2,698 FNs, enhancing the F1-score by 2.76%. Similarly, in the BCM Iso-Seq HG004 dataset, the number of FPs decreased by 4334 and FNs by 4331, leading to an improvement of 2.69% in the F1-score. Moreover, the impact of disregarding zygosity lessened as DP increased. For instance, in the ONT dRNA HG004 dataset, setting DP≥8 while considering zygosity boosted the F1-score by 4.19% over DP≥4, whereas disregarding genotyping resulted in a 2.45% F1-score improvement. These results emphasize the importance of read coverage in variant detection, especially when incorrect genotypes are introduced without adequate reads to determine variant zygosities.

### Performance analysis by genomic context

We also carried out benchmarking in challenging genomic regions. Specifically, we compared the SNP performance of Clair3-RNA and LongcallR in areas defined as difficult by GIAB Stratification V3 [25]. We categorized the analysis into five groups: “Low complexity”, “Segmental duplications”, “Low mappability”, “Functional regions”, and “Other difficult regions” − and conducted it on ONT dRNA004 HG004, PacBio BCM Iso-Seq HG004, and PacBio MAS-Seq HG004 datasets. As shown in **Figure 4b** and **Supplementary Table 6**, Clair3-RNA consistently outperformed LongcallR in SNP F1-score.

Benchmarking on the ONT dRNA004 dataset, Clair3-RNA demonstrated varying performance, achieving an average 80% F1-score. Notably, Clair3-RNA excelled in coding regions, attaining a 94.58% F1-score in the Coding Sequence (CDS) regions. In contrast, its performance dropped to 61.80% in low complexity regions characterized by homopolymer regions greater than 11 bp. In comparison, LongcallR exhibited more significant variability on the same dataset, peaking at 82.51% in the CDS regions, while dropping to 29.85% in homopolymer regions greater than 11 bp. Similar trends were observed in the PacBio Iso-Seq and MAS-Seq datasets, Clair3-RNA’s F1-score averaged ∼86%, with the highest at 95.22% in the CDS regions and the lowest at 74.76% in the homopolymer regions. In this dataset, LongcallR averaged about 66%, with a peak of 91.74% in the CDS regions and a low of 37.07% in the homopolymer regions. Among all categories, “Low complexity” and “Segmental duplications” were the most challenging for variant detection, with the lowest average F1-score of all categories.

### Indel performance

In our evaluation, we compared the performance of various callers, excluding LongcallR, which identifies only SNP variants. For the ONT platform, as shown in **Supplementary Table 2**, using DP≥10 and AD≥4 coverage settings, Clair3-RNA achieved HG004/66.50% and HG005/72.37% F1-score for the dRNA004 dataset, and HG004/50.63% and HG005/60.88% for the cDNA dataset. It consistently outperformed both Clair3 (dRNA004 HG004/52.65% and HG005/60.64%, cDNA HG004/45.63% and HG005/55.04%) and DeepVariant (dRNA004 HG004/40.03% and HG005 44.94%, cDNA HG004/36.82% and HG005 39.45%) in both the cDNA and dRNA004 datasets. In a comparison between the dRNA004 and dRNA002 datasets, Clair3-RNA demonstrated a significant performance improvement of 43.11%, with the average F1-score increasing from 26.33% to 69.44%. Even in the most precise dRNA004 dataset, the averaged indel recall was lower than 60%, attributed primarily to the presence of indel signals with varying lengths clustered at the same site, confusing the model from calling Indel precisely.

On the PacBio platform, as shown in **Supplementary Table 3,** using the DP≥10 and AD≥4 coverage setting, Clair3-RNA achieved an Indel F1-score of HG004/89.42% and HG005/92.90% for BCM Iso-Seq, HG004/82.33% and HG005/82.82% for Google Iso-Seq, and HG004/80.05% for BCM MAS-Seq. With an average indel performance of 85.50%, Clair3-RNA significantly outperformed Clair3 and DeepVariant, where the average F1-score was 63.25% (+22.25%) and 40.31% (+45.19%), respectively.

### RNA editing tagging using the REDIportal database

The REDIportal has compiled a total of 36,234,771 RNA editing sites, categorized as follows: 1) 13,101,296 editing sites collected only in ATLAS (A); 2) 22,836,582 editing sites collected by ATLAS and RADAR (A&R); 3) 4,708 editing sites collected by ATLAS and DARNED (A&D), and 4) 292,185 editing sites reported by all three sources (A&R&D). We tagged the variant as an RNA editing site that was validated by at least two independent sources within the REDIportal, specifically in ATLAS, DARNED, and RADAR. As shown in **Supplementary Table 4**, excluding the variants tagged by the REDIportal database, there was enhanced SNP performance across all platforms. Specifically, Clair3-RNA achieved a 0.82% improvement in the ONT dRNA004 HG004 dataset, with a total of 1,898 editing sites tagged. For the PacBio datasets, the improvement was less significant, with 1,033 and 912 editing sites tagged in the BCM Iso-Seq HG004 and MAS-Seq HG004 datasets, respectively, resulting in a 0.28% and 0.13% F1-score improvement.

### A-to-G and T-to-C false positives analysis

A-to-I RNA editing poses a significant challenge for RNA variant calling algorithms because they must distinguish between A-to-T and T-to-C SNPs induced by editing events and genuine A-T and T-to-C SNPs. To address this issue, we analyzed A-to-G and T-C SNPs identified by Clair3-RNA and Clair3 that were not present in the GIAB ground truth.

The high-quality benchmark set of the editing sites were established and analyzed using the PacBio HG004 BCM Iso-Seq dataset and the ONT HG004 dRNA004 dataset. The dRNA004 dataset was aligned with the HG004 DNA dataset, sequenced by ONT R10.4.1 Q20+ chemistries, and basecalled using Dorado, provided by Nanopore EPI2ME Labs [26]. For the BCM HG004 dataset, we compared the RNA dataset with the PacBio HiFi HG004 DNA dataset generated by Google [27]. We then identified potential A-to-I editing sites by looking for evident A-to-T or T-to-C substitutions in the RNA data that were absent in the DNA data. More details of the benchmarking editing site generation are described in the **Methods** - **Orthogonal high-quality RNA editing benchmark set generation** section. Finally, we identified 3,800 and 6,086 editing sites on the ONT dRNA004 and PacBio Iso-Seq datasets, respectively. 46.21% (1,756 over 3,800) and 55.71% (3,391 over 6,086) of the benchmark set of editing sites were found in the REDIportal database.

In the Venn diagrams presented in **Figure 5b**, the A-to-G and T-to-C false positives identified by Clair3-RNA and Clair3 were extracted for comparison. Clair3-RNA reported 3,940 false positives in ONT and 3,979 in PacBio, significantly fewer than Clair3, which identified 30,411 false positives in ONT and 94,476 in PacBio. Among these false positives, 1,251 (539+712) and 409 (81+328) variants intersected with the benchmark set of editing sites for ONT and PacBio, respectively, indicating that Clair3-RNA can classify 2,549 (1,505+1,044) and 5,677 (2,614+3,063) high-quality RNA editing sites as non-variants through network prediction. In comparison, Clair3 detected only 1,592 and 1,011 RNA editing sites for ONT and PacBio, respectively. The results indicate that Clair3-RNA demonstrates a capability in rejecting RNA editing sites during variant detection. Notably, the latest ONT basecaller, Dorado v0.9.0, released in Dec 2024, has significantly enhanced its ability to identify modified bases. Additionally, it can now directly call A-to-I editing sites as an ‘A’. Although the new basecaller was release after we had completed drafting the manuscript, so it wasn’t included in our benchmark, we believe that the number of high-quality RNA editing sites identifiable in the ONT dataset can be substantially increased, particularly when using dRNA data only. As shown in **Figure 5c**, for the remaining false positives in Clair3-RNA, 29.87% (1,177 over 3,940) in ONT and 26.94% (1,072 over 3,979) in PacBio were annotated by the REDIportal database. After network prediction and REDIportal database tagging, 62.77% of the false positives in the PacBio dataset and 38.37% of the false positives in ONT were identified. After network prediction and editing site tagging by the REDIportal database, the remaining A-to-G and T-to-C false positives can be reduced from 27,175 to 2,051 (1,512+539) on ONT and from 54,942 to 2,579 (2,498+81) on the PacBio platform. The results suggest the feasibility of utilizing the REDIportal database to annotate challenging editing sites that are missed by network prediction.

**Figure 5.**
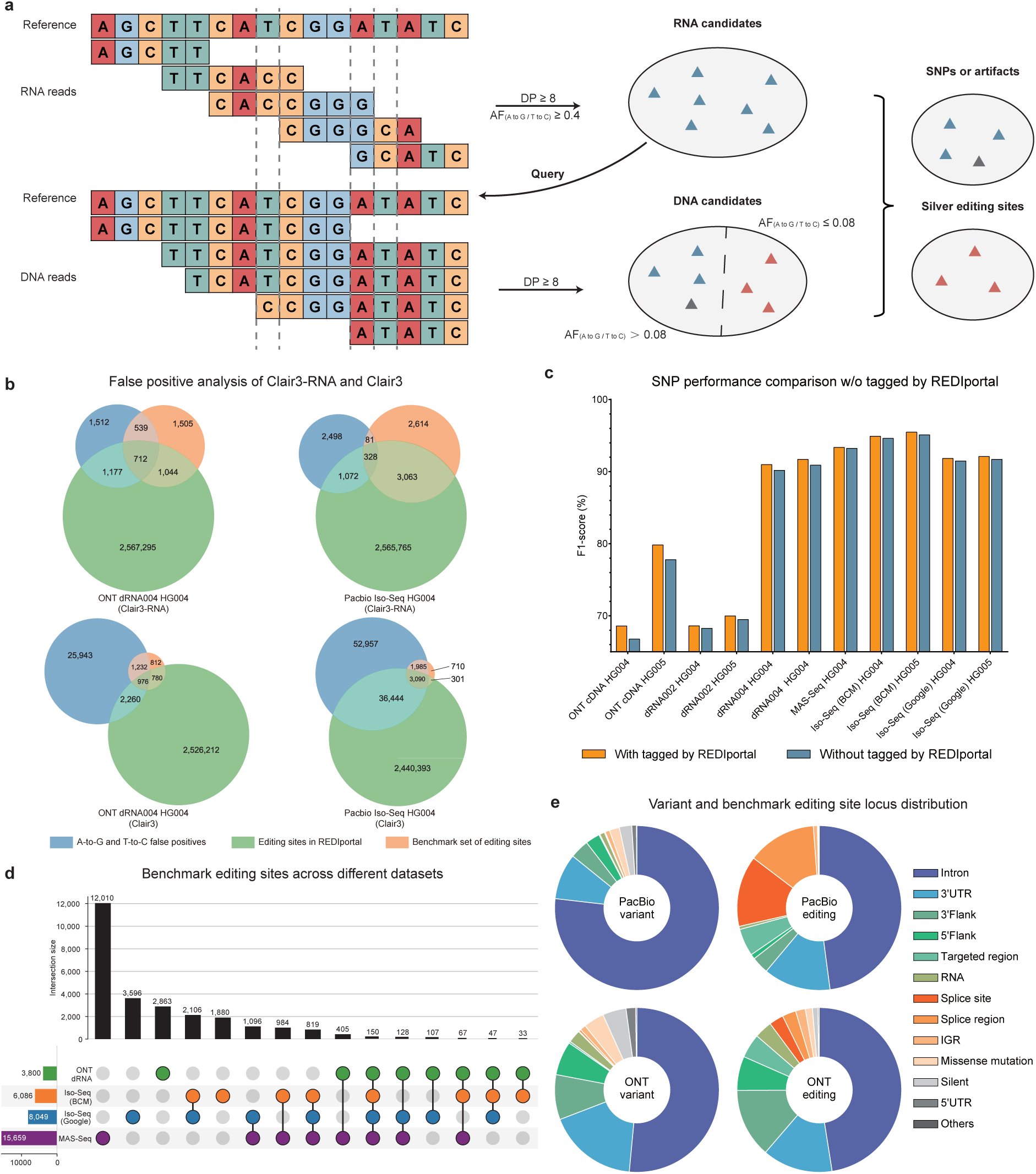
RNA editing event analysis. (a) This illustrates the workflow of acquiring high-quality benchmark set of editing sites. First, candidate sites are assessed based on coverage depth and allelic fraction at various positions in RNA alignments. A site qualifies as a high-quality editing site candidate if the coverage depth exceeds 8and the allelic fraction from A to G or T to C is above 0.75. These candidates are then cross-referenced with DNA alignments. If a position in DNA has a depth exceeding 8 and the allelic fraction from A to G or T to C is below 0.08, it is deemed a high-quality editing site; otherwise, it is classified as an SNP or an artifact. (b) Venn diagrams are presented for three sets: editing sites in REDIportal, A-to-G and T-to-C false positives, and the benchmark set of editing sites. A smaller overlap between A-to-G and T-to-C false positives and detected editing sites indicates Clair3-RNA’s capability of distinguishing genuine SNPs and A-to-I editing sites. (c) This provides an SNP performance comparison across different datasets when with (w) and without (o) variants tagged by the REDIportal database. (d) The upset plot illustrates the distribution of benchmark set of editing sites across different platforms. (e) This presents the distribution of benchmark editing sites and variants within the PacBio Iso-Seq (BCM) HG004 dataset and the ONT dRNA004 HG004 dataset.

### Editing sites comparison and analysis

We identified the high-quality editing sites across additional datasets and analyzed their intersections, as depicted in **Figure 5d**. In the Iso-Seq (Google) dataset, we identified 8,049 sites, while the MAS-Seq dataset yielded 15,659 sites. It is noteworthy that the ONT dRNA004 dataset contained fewer sites compared to the PacBio datasets, primarily due to the absence of a reverse transcription step in the ONT protocol. This is in contrast to PacBio’s cDNA preparation, which includes the reverse transcription step as shown in **Supplementary Figure 2**. Additionally, the MAS-Seq dataset exhibited a significantly higher number of sites than the other datasets, which can be attributed to its broader genomic region coverage. This resulted in a greater number of detected variants on the MAS-Seq dataset. Among the four datasets, the PacBio datasets demonstrated a higher degree of overlap with each other than with the ONT dataset, highlighting the distinct methodological differences between ONT dRNA and PacBio cDNA sequencing approaches.

We used the vcf2maf tool [28] to annotate the variants identified by Clair3-RNA and editing sites in the PacBio Iso-Seq HG004 and ONT dRNA004 HG004 datasets. As shown in **Figure 5e and Supplementary Table 7**, both the variants and editing sites were found to be predominantly located in intronic regions. About half of the PacBio editing sites, ONT variants, and ONT editing sites were in introns, while over three-quarters of PacBio variants were in intron. Editing sites, however, occur more often in splice sites and splice regions. Specifically, 27.43% of the editing sites were in these regions, compared to just 0.19% of the variants. In the ONT dRNA004 dataset, 5.50% of the editing sites but 0.53% of the variants, were found in splice sites and splice regions. These findings align with research on the A-to-I editing survey [16, 29].

### Enhancing variant calling performance with phasing information

A heterozygous variant identified in a single haplotype is considered more reliable than sequencing errors that affect both haplotypes [13]. Based on this, we attempted to incorporate phasing haplotype information into the neural network, with more details provided in the **Methods** - **Integrating phasing information into neural network** section. In the variant phasing phase, as illustrated in **Figure 6d**, we analyzed phased heterozygous variants and found that at least 63.34% of heterozygous variants can be phased. The percentage rises to 78.05% and 79.07% when using the latest PacBio MAS-Seq HG004 and ONT dRNA004 HG004 datasets, respectively. These datasets have average read lengths of 1,731 and 1,404 for variant phasing.

**Figure 6.**
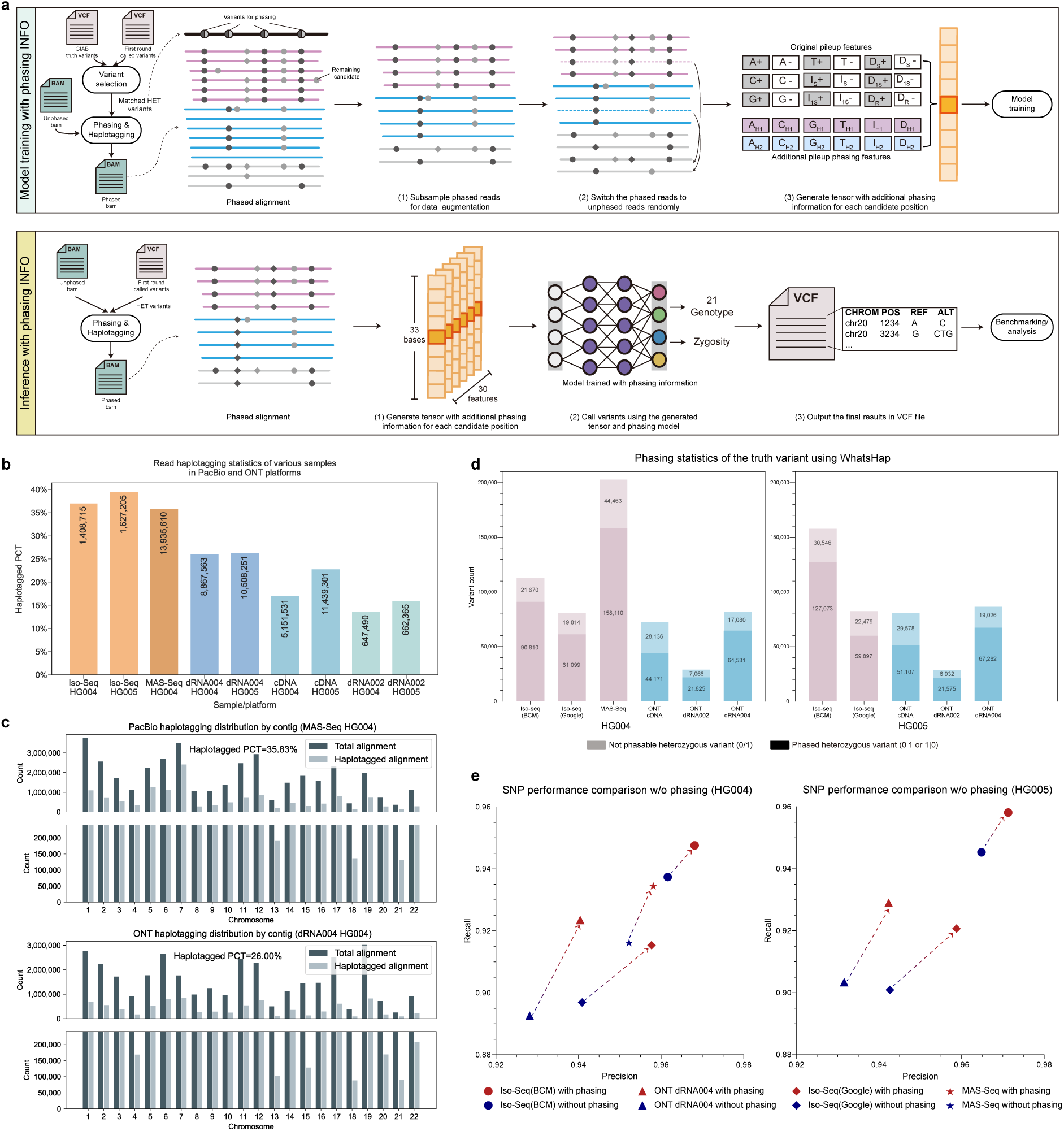
Phasing result analysis in lrRNA-seq. (a) This figure illustrates the workflow of Clair3-RNA for model training and inference incorporating phasing information. During the training phase, variants were phased using the first round of called variants in conjunction with GIAB truths, and the alignments were haplotagged based on these phased variants. The alignments are color-coded, with pink representing haplotype 1, blue for haplotype 2, and grey indicating unknown haplotypes. The circle and rhombus in the alignments represent SNPs and indels, respectively. Twelve phasing features were concatenated with the original eighteen features and subsequently input into the neural network for model training. In the inference phase, heterozygous variants from the first round of called variants of Clair3-RNA were utilized for variant phasing. (b) This panel presents the read haplotagging statistics for various samples across PacBio and ONT platforms. The value on y axis refers to the percentage of haplotagged reads, and the numbers displayed in each histogram represent the total count of haplotagged reads within the respective subset. (c) This panel displays read haplotagging statistics, stratified by chromosomes, for PacBio MAS-Seq HG004 and ONT dRNA004 HG004 samples. (d) This panel presents variant phasing statistics for truth variants analyzed using WhatsHap across various PacBio and ONT samples. (e) The precision-recall curve of the SNP performance across various samples, comparing the performance with and without phasing information in the neural network.

For read haplotagging, we computed the percentage of haplotagged reads against all sequencing reads, while excluding supplementary reads, secondary reads, and reads with a mapping quality lower than 5, as recommended by WhatsHap[30]. **Figure 6b** and **Figure 6c** display the haplotagged percentages of all datasets and the number of haplotagged reads - for PacBio MAS-Seq HG004 and ONT dRNA004 HG004 datasets. The ONT dRNA004 achieved an average haplotagging rate of 26.17%, with PacBio MAS-Seq 35.83%, and Iso-Seq 38.22%. The haplotagging rates of lrRNA-seq are significantly higher than those of the short-read datasets, which is typically less than 10%. This suggests the potential for enhancing variant detection by incorporating phasing information.

We trained the Clair3-RNA pileup model with phasing information from both the PacBio and ONT platforms using the GIAB HG002 dataset. Subsequently, we evaluated the trained models on HG004 and HG005 datasets. As shown in **Figure 6e** and **Supplementary Table 8**, both SNP and Indel precision and recall improved across all datasets. Benchmarking results for the ONT and PacBio datasets indicate an overall improvement of 1.52% in the F1-score for SNP. Specifically, using the PacBio MAS-Seq HG004 dataset, the F1-score increased by 1.23% (94.61% versus 93.38%). The improvement was more significant in the ONT dRNA004 dataset, with an average 2.01% F1-score increase. For HG004, the F1-score increased from 91.73% to 93.56%, and for HG005, it increased from 91.00% to 93.19%. With phasing information, the F1-score reached 98.59% for PacBio MAS-Seq, 98.24% for Iso-Seq, and 97.16% for ONT dRNA004, with at least 10 supporting reads and disregarding zygosity switches.

The improvement in Indel detection is more significant compared to SNP, achieving an average 5.86% F1-score improvement across all benchmarking datasets. PacBio Iso-Seq demonstrates greater improvement than MAS-Seq (6.54% versus 5.01%) and ONT dRNA004 (6.54% versus 4.92%). The Indel performance reached 85.61% F1-score in the PacBio Iso-Seq HG005 dataset, representing a 4.21% improvement. ONT dRNA004 HG005 achieved a 70.63% F1-score with a 4.58% improvement. By increasing the minimum thresholds to DP 10, AD 4, and disregarding zygosity switches, the average Indel F1-score can reach 89.42% for PacBio and 72.56% for ONT. Notably, the highest Indel F1-score achieved was 95.17% for PacBio Iso-Seq HG005 and 75.51% for ONT dRNA004 HG005.

### GENCODE annotated gene analysis

We evaluated the Clair3-RNA variant calls across gene regions to assess its applicability for functional studies in lrRNA-seq. We used the annotated genes from the GENCODE (version 46) database [31]. For gene region analysis, we selected those genes that overlapped the defined callable regions specified in the **Methods** - **Benchmarking metrics** section.

We first categorized all callable genes into three distinct groups based on their overlap with callable high-confidence regions. This overlap ratio *H* was computed how many percent of a gene is in callable high-confidence regions, defined as the intersected regions of the callable regions and GIAB high-confidence regions. More details are given in the **Methods** - **Analysis of overlap and error statistics for GENCODE annotated gene** section. Genes were classified as wholly (*H*=100%), partially (75%<*H*<100%), or not (*H*=0%). We then analyzed how many genes are with variants covered and has no calling mistake in each dataset. Furthermore, we compared these genes to the phasesets computed by WhatsHap to determine how many of them are wholly phased.

Figure 7a presents the number of genes covered by callable high-confidence regions for both the PacBio MAS-Seq HG004 and ONT dRNA004 HG004 datasets. For PacBio, a total of 3,859 genes were fully covered, 3,589 were partially covered, and 1,135 were not included in defined callable high-confidence regions. In contrast, the ONT platform showed 2,199 fully covered genes, 1,870 partially covered, and 876 not included in callable high-confidence regions. Further analysis revealed the number of genes without SNP error. In the PacBio dataset, 1,279 genes (33.1% of fully covered) and 2,404 genes (67.0% of partially covered) were without SNP error. For the ONT platform, 735 genes (33.4% of fully covered) and 1,167 genes (62.4% of partially covered) were without SNP error. Note that the SNP errors in the "not in callable high-confidence region" category could not be determined, because they fall outside the GIAB high-confidence region.

**Figure 7.**
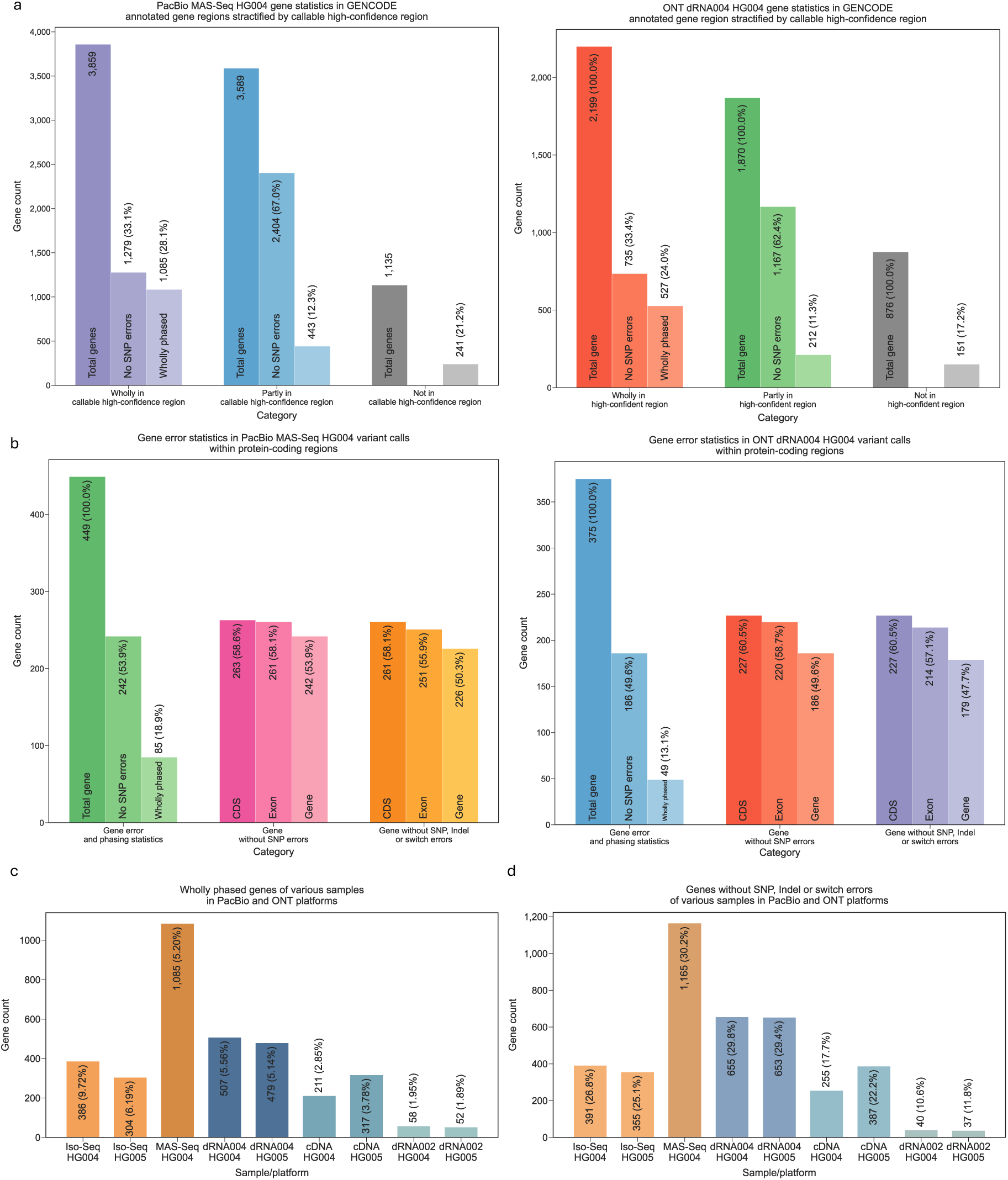
GENCODE annotated gene analysis. (a) Gene statistics for PacBio MAS-Seq HG004 and ONT dRNA004 HG004 are presented, stratified by GENCODE annotated gene regions. The data is categorized into wholly phased genes and those with no SNP errors, based on their overlap with callable high-confidence regions, with percentages reflecting the proportion of each category relative to its predecessor. (b) This panel shows gene error statistics for variant calls within protein-coding regions for both PacBio MAS-Seq HG004 and ONT dRNA004 HG004 datasets. The percentages are calculated relative to the total number of genes in the subset. (d) The number of wholly phased genes across various samples from PacBio and ONT platforms is illustrated, with percentages relative to the total number of annotated genes within the callable high-confidence regions. (e) This panel depicts genes without SNP, Indel, or switch errors from various samples in PacBio and ONT platforms, with percentages again relative to the total annotated genes in the callable high-confidence regions.

**Figure 8.**
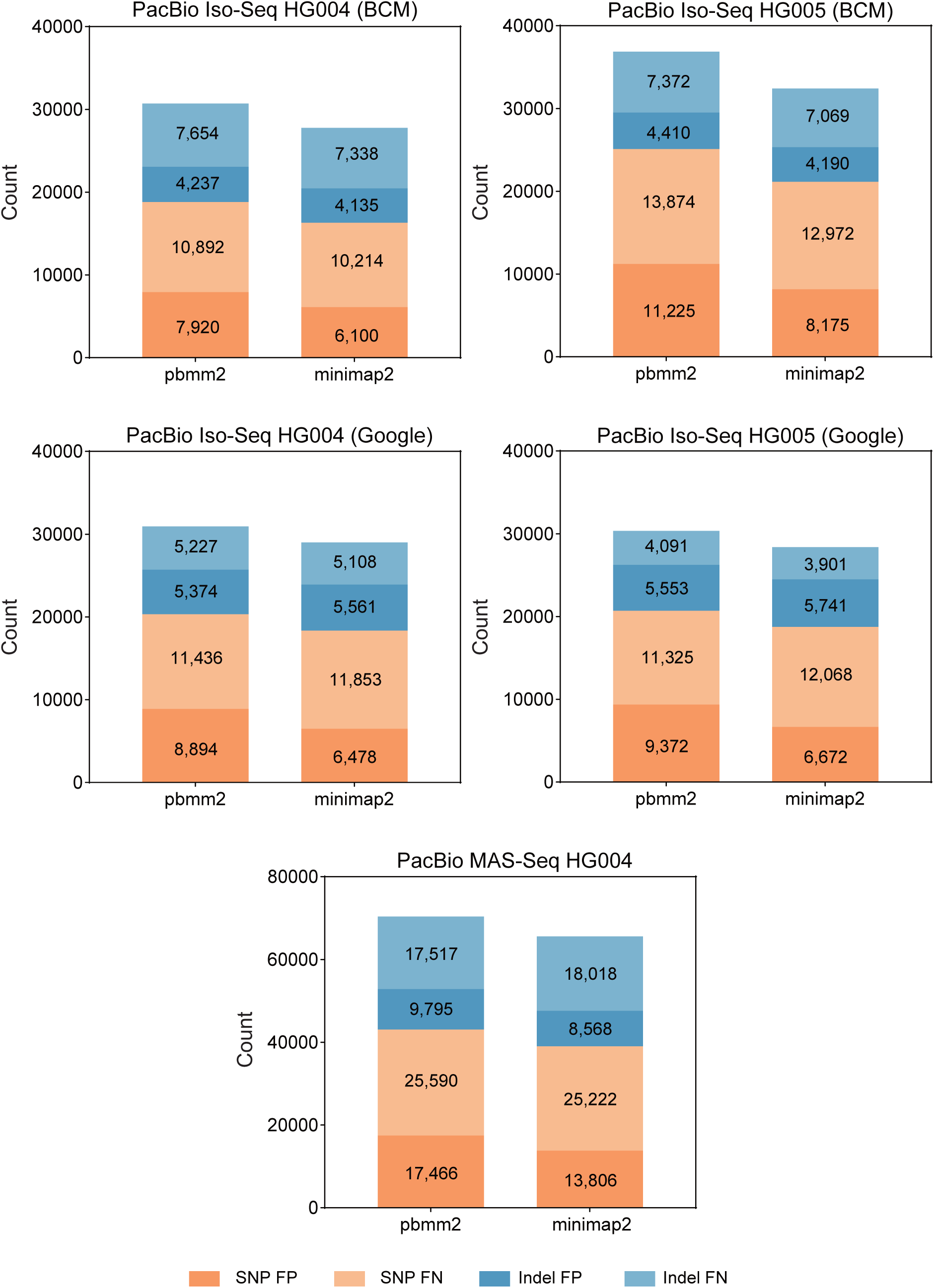
Clair3-RNA PacBio performance using different aligners. This is a comparison of false positive (FP) and false negative (FN) using different alignment methods on PacBio datasets. minimap2 aligner exhibits lower FPs and FNs than pbmm2 in benchmarking datasets. The alignment parameter settings for pbmm2 and minimap2 are described in the **Supplementary Notes − Command line used** section.

We also summarized the numbers by looking into only those genes annotated as “protein_coding” in the GENCODE database, which are more important for studying protein synthesis and biological functions. As shown in Figure 7b, we identified a total of 449 and 375 protein-coding genes in PacBio and ONT, respectively. Among these, 242 and 186 genes were detected without SNP error, while 85 and 49 genes were wholly phased in PacBio and ONT. When examining only the coding sequence (CDS) regions, we found that 263 genes (58.6% of the protein-coding genes) in PacBio and 227 genes (58.7%) in ONT were free of SNP errors. Considering only the exon regions, 242 genes (53.9% of the protein-coding genes) in PacBio and 220 genes (58.7%) in ONT had no SNP errors. We further restricted our analysis to genes without SNPs, Indels, or switch errors. In this context, we observed that 261 genes (58.1%) in CDS and 251 genes (55.9%) in exons for PacBio, as well as 227 genes (60.0%) in CDS and 214 genes (57.1%) in exons for ONT, met this criterion. These results indicate a promising variant calling accuracy in the protein-coding regions.

Figure 7c presents the number of wholly phased genes across all benchmarking datasets. The MAS-Seq HG004 identified 1,085 wholly phased genes, exceeding 386 and 304 genes reported by the Iso-Seq HG004 and HG005, respectively. Similarly, ONT dRNA004 identified 504 and 479 wholly phased genes in HG004 and HG005, respectively, compared to 211 and 317 in the cDNA datasets and 58 and 52 in the dRNA002 datasets. Figure 7d illustrates the number of genes free from SNPs, Indels, or switch errors, with MAS-Seq leading at 1,165 genes, followed by dRNA004 with 655 in HG004 and 653 in HG005. The substantial numbers of wholly phased and error-free genes highlight the potential for reliable downstream gene analysis, particularly using the PacBio MAS-Seq and ONT dRNA004 technologies.

### Benchmarking runtime and memory usage

As shown in Figure 3e and **Supplementary Table 9**, we evaluated the runtime and memory usage of the Clair3-RNA algorithm and other variant callers. Clair3-RNA demonstrated an outperforming runtime against other variant callers. Specifically, using 16 CPU cores, Clair3-RNA required ∼18 minutes and ∼50 minutes, on HG004 PacBio Iso-Seq and MAS-Seq, and ∼34 minutes and ∼29 minutes on HG004 ONT cDNA and dRNA004 dataset, respectively. This is generally faster than Clair3, LongcallR, and DeepVariant, in which the average runtime ranges from 42 to 80 minutes. Clair3-RNA also showcased uniform memory consumption across all datasets, with a footprint of ∼7 GB. In contrast, DeepVariant required about 26 GB and 24 GB of memory for the PacBio and ONT, respectively. Clair3 displayed a slightly higher memory footprint, on average 49 GB, due to its variant phasing process. In comparison, LongcallR recorded the highest memory consumption at PacBio MAS-Seq, with 83 GB, and averaged 47 GB.

## Discussion

Variant calling in RNA-seq datasets offers more comprehensive insights into the transcriptome analysis. However, RNA-seq comes with higher error rates, making it challenging to differentiate between sequencing artifacts and true variants. RNA reads also display imbalanced coverage distribution across the reference genome, making it difficult to detect low-frequency variants. Additionally, A-to-I RNA editing events present a significant challenge in RNA-seq analysis, often resulting in higher false positives.

In this study, we introduced Clair3-RNA, the first deep learning-based variant caller designed for long-read RNA sequencing. Clair3-RNA utilizes a pileup neural network and incorporates various techniques, such as coverage normalization, variant zygosity switching, and editing site tagging, for accurate variant detection in lrRNA-seq. Across different platforms, including PacBio Iso-Seq, PacBio MAS-Seq, and ONT dRNA004, Clair3-RNA successfully identified ∼90% of true RNA variants with a minimum four reads support, and further improved to 92% when the minimum read coverage was increased to 10 and zygosity information was disregarded, showing its capability for accurate variant detection in lrRNA-seq. By identifying known editing sites and analyzing A-to-G and T-to-C SNP false positives, we also found that Clair3-RNA was capable of distinguishing major A-to-I editing sites from genuine variants, which is challenging for other DNA variant callers. We further investigated the incorporation of phasing into the Clair3-RNA architecture and achieved a ∼2% F1-score improvement in our benchmarking, demonstrating the advantages of haplotype phasing in lrRNA-seq.

Clair3-RNA demonstrates greater SNP accuracy than other variant callers across all platforms and sequencing scenarios. Nevertheless, the accuracy of indel calling remains challenging due to allele coverage fluctuations and increased error rates. Enhancing Indel accuracy could involve incorporating techniques such as local assembly for Indel realignment and local haplotype assembly to differentiate low-frequency Indels from sequencing artifacts, leading to more precise Indel detection. Moreover, although Clair3-RNA successfully identified more than 2k and 5k of the editing sites in the ONT dRNA004 and PacBio Iso-Seq, respectively. As the false positive distribution shown in **Supplementary Figure 3**, the presence of RNA false positives was linked primarily to excessively low allelic fraction RNA editing sites or high allelic fraction sequencing artifacts that the neural network failed to capture. These editing sites exhibited subtle differences against true variants at the base level, making them indistinguishable from genuine A-to-G or T-to-C variants without additional contextual information. Further attempts will involve integrating raw sequencing signal features for editing site detection, especially in dRNA sequencing, as the A-to-I RNA editing event is more distinct at the raw signal level. Dorado also supports outputting modified bases in its latest release [24], where the Inosine will be basecalled as canonical A with additional modification tag, thereby avoiding editing errors in downstream variant calling. Furthermore, in our model architecture design, we utilized a recurrent neural network to encode the pileup features. As more RNA data becomes accessible, a meticulously crafted transformer-based architecture might further improve the performance. The attention module within a transformer-based network exhibits the capacity to capture the connections in a distant flanking window, thereby enhancing accuracy in challenging regions.

## Methods

### Overview of Clair3-RNA

Figure 1 shows an overview of the Clair3-RNA workflow. Starting from the read alignments in BAM/CRAM format of an input sample, Clair3-RNA identifies potential variants with adequate read support and base alteration. Leveraging the pileup methodology inherited from Clair3, the candidates and their corresponding flanking window sites are aggregated to create pileup tensors, which are then processed through a Bi-LSTM-based pileup neural network for inference. The neural network produces probabilistic multi-task output that differentiates variants from editing sites and sequencing artifacts. Identified variants are further refined by the editing tagging module to tag known editing sites. Clair3-RNA is compatible with various platforms, including PacBio Sequel with the Iso-Seq kit, PacBio Revio with the MAS-Seq kit, ONT cDNA, ONT dRNA002, and ONT dRNA004.

### Clair3-RNA’s input and output

#### Input

Clair3-RNA takes the input, which includes two components: 1) a reference genome of the sequencing sample, and 2) a sequencing alignment with the same reference material as 1). Clair3-RNA automates the whole variant calling process to get the **VCF output**, as illustrated below. The pileup tensor consists of 594 integers, covering 33 genome positions, which include the candidate position and 16 flanking positions on both the left and right sides, with 18 features at each position. These 18 pileup features are designated as "A+", "C+", "G+", "T+", "I_S_+", "I_1S_+", "D_S_+", "D_1S_+", "D_R_+", "A-", "C-", "G-", "T-", "I_S_-", "I_1S_-", "D_S_-", "D_1S_-", and "D_R_-". A, C, G, T, I, and D represent the counts of read support for the four nucleotides, insertion and deletion, while the symbols "+" and "-" represent the read from the forward and reverse strands, respectively. The superscripts "S" and "R" represent the start and remaining positions of Indel. A detailed description of the 18 pileup features is provided in the **Supplementary Notes** − **Description of RNA pileup input features** section. All the features above worked with sites with the reads covered, and we assigned a background intensity of 0 for all padding sites. This technique of zero-padding is also utilized in RNA sequencing to account for surrounding RNA splicing regions.

#### Clair3-RNA model architecture

The Clair3-RNA pileup network incorporates two bidirectional long short-term memory (Bi-LSTM) layers and three fully connected layers to encode sequential pileup features. The output of the pileup model encompasses two probabilistic tasks: 21-genotype and zygosity. Further details about the output tasks are provided in the **Supplementary Notes** − **Description of pileup network outputs** section. The identified variants, encompassing both nucleotide alterations and genotypes, were determined by evaluating the combined probabilities from two output tasks. In the case of an Indel variant, the network specifies the variant type, and the length of the Indel is inferred from the supporting read evidence.

#### VCF output

Three categories of variants are shown in Clair3-RNA’s VCF output: 1) variants labelled as "PASS" that passed quality score threshold, 2) labelled as "LowQual" that has low-quality scores (i.e., QUAL < 2 for PacBio and QUAL < 8 for ONT, configurable), and 3) tagged as "RNAEditing" corresponded to known editing sites identified by the REDIportal database. Specifically, variants were tagged as "RNAEditing" if they match both the reference and alternate alleles given in REDIportal. For each variant entry, the reference, alternate variant, genotype, coverage of the reference allele, and coverage of all alternate alleles are shown.

### Training techniques tailored for RNA data

#### Training candidate generation

In DNA model training, the ground truths were established using the golden GIAB truth variants. However, in RNA model training, owing to limited sequencing coverage in specific regions, not all truth variants are suitable for training. Initially, we scanned the training dataset to acquire the read coverage (DP) and the alternate allele (AD) supporting truth variants, focusing on evident variants with adequate read support. For the artifacts, we scanned the alignment site-by-site to identify candidates with DP⩾4, AD⩾2, and exceeding a minor allelic fraction (typically 0.08, which was manually decided). Additionally, we accounted for zygosity switches and editing sites in the training data, as elaborated in the **Methods** - **Variant zygosity switching** section.

#### Variant zygosity switching

Zygosity switching in RNA data can lead to misinterpretations of genetic variants. This occurs when a heterozygous variant appears as a homozygous reference or a homozygous variant due to allelic imbalance. Zygosity switching in RNA data might be due to differential gene expression when one allele dominates in specific regions. Additionally, RNA splicing might create isoforms, altering the variant zygosity across different isoforms [5]. Ignoring zygosity switching in RNA data can have a significant impact on model training. Therefore, before training the model, we computed the alternate allelic fraction of each candidate and adjusted the zygosity obtained from GIAB based on DNA sequencing samples. By defining each observed candidate allelic fraction as VAF_O_ and the truth zygosity as *GT_Truth_*, where *GT*∈{0/1, 1/1, 1/2}, we can determine the inferred zygosity *GT_Inferred_*, as follows:

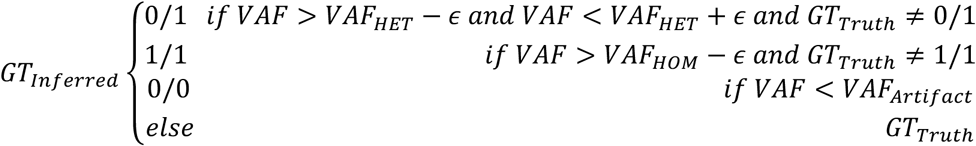

where *GT_Truth_* is the observed ground truth genotype in GIAB truths. *VAF_HET_*, *VAF_HOM_*, and *VAF_Artifact_* represent the expected allelic fraction for heterozygous variants, homozygous variants, and sequencing artifacts were set as 0.5, 1.0, and 0.08, respectively. *є* represents the tolerance epsilon parameter and was manually set as 0.2. For multi-allelic 1/2 genotypes, we considered each allele and inferred the genotype when any of the alternate alleles was missing. Finally, we used *GT_Inferred_* instead of *GT_Truth_* for constructing zygosity and 21-genotype labels during model training.

### Coverage normalization for extremely high-coverage data

RNA data exhibits an uneven distribution of sequencing read coverage, stemming primarily from variations in transcript abundance, non-uniform RNA expression, and RNA splicing events. These factors result in uneven coverage compared to homogeneous DNA sequencing data. The presence of high-coverage sites poses challenges for model inference, as the training data may not adequately represent the full spectrum of coverage distribution. Although Clair3-RNA has implemented techniques like Focal Loss [32] to address data imbalances during model training, excessively high coverage levels can significantly impede the accuracy of model inference.

One potential strategy involves subsampling reads during inference to mitigate the gap; this approach, however, may impact performance due to less alignment included in inferences. Building upon this, we proposed a coverage normalization strategy. Initially, we computed the coverage distribution of the training sample and excluded the top 10% of high-coverage sites, identifying the maximum coverage confidence interval as *C_T_*. In the inference phase, we determined the coverage of each candidate tensor *C_I_* by computing the reads supporting the nucleotide A, C, G, T, and deletions.

Subsequently, for each candidate coverage *C_I_* exceeding *C_T_*, we computed the normalized pileup features *F_pileup_norm_* using the formula:

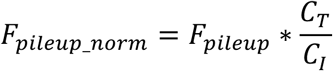

where *F_pileup_* is the total 594 pileup integers used for prediction. In our experiment, using the normalized feature generally improved recall by ∼1%, effectively mitigating the influence of extremely high coverage on the model’s inference.

### Orthogonal high-quality RNA editing benchmark set generation

RNA A-to-I editing is a post-transcriptional modification, where adenosine (A) residues are deaminated to inosine (I) by adenosine deaminases acting on RNA (ADAR) enzymes. The editing process can alter the RNA structure and function, impacting gene expression and protein diversity [16]. In sequence alignments, A-to-I during editing is interpreted as a G by reverse transcription. RNA A-to-I editing is often represented as an A-to-G change on the edited RNA strand and as a T-to-C change on the corresponding DNA template strand. dRNA sequencing detects A-to-I editing directly from RNA data, as it omits the reverse transcription step, offering a precise view of RNA modifications without potential artifacts.

While no definitive editing site reference is available from GIAB, we generated a high-quality benchmark set of editing sites by cross-validating potential editing sites with DNA sequencing data, as shown in Figure 5a. The rationale is that an editing site is determined if present in RNA data but absent in DNA data. Initially, we identified A-to-G and T-to-C candidates from RNA alignments and computed read support in the corresponding DNA sequencing data. If the sequencing coverage (DP) and variant allelic fraction (VAF) supporting either editing allele (G or C) in RNA (R) and DNA (D) is calculated as *DP_R_*, *VAF_R_*, *DP_D_*, and *VAF_D_*, respectively, then the editing site is decided as follows:

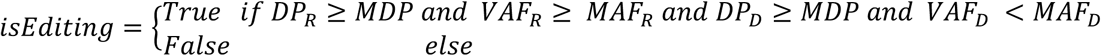

where *MDP* represents minimal coverage depth, and *VAF_D_* and *VAF_R_* represent the VAF observed in DNA and RNA, respectively. *MAF_D_* represents maximal allelic fraction supporting A-to-C or G-to-C in DNA, and *MAF_R_* represents minimal allelic fraction supporting A-to-C or G-to-C in RNA. We set *MDP*, *MAF_D_*, and *MAF_R_* as 8, 0.08, and 0.75, respectively, for both the PacBio and ONT experiments. **Supplementary Figure 1** presents the detected allele frequency distribution of high-quality RNA editing sites across PacBio and ONT datasets.

We incorporated candidates from the GIAB high-confidence regions for model training. These candidates were categorized in RNA BAM files as RNA variants, RNA editing sites, or artifacts. Throughout our experiments, we explored the inclusion of a separate RNA editing site task to identify RNA edits. Additionally, we tried incorporating the editing sites as an additional category within the zygosity task. We found that both approaches failed to accurately identify true RNA variants compared to directly integrating editing sites into the homozygous reference category. This indicates that the model is able to classify the editing sites as references in model training.

### RNA editing site tagging using the REDIportal database

REDIportal is a comprehensive database that consolidates a vast array of RNA editing sites sourced from three main repositories: ATLAS, DARNED, and RADAR. Within REDIportal v2, approximately 40 million RNA editing sites are documented. In Clair3-RNA, variants are tagged as RNA editing sites if they correspond to A-to-G or T-to-C alterations found in the REDIportal database, ensuring a matching alternate representation. REDIportal prioritizes RNA editing sites validated by agreement of at least two of its constituent sources (ATLAS, DARNED, RADAR), with adjustable configuration setting for users’ customs. Clair3-RNA supports REDIportal versions customized for both the GRCh37 and GRCh38 reference genomes, providing a reliable resource for exploring established RNA editing events.

### Incorporating phasing information into neural network

Haplotype assembly, or phasing, is a fundamental technique for reconstructing an individual’s haplotypes [30, 33]. Long-read sequencing generates reads spanning tens of kilobases or more, enabling the capture of haplotype information often lost in short-read sequencing data. This method has been shown to significantly enhance variant calling performance in long-read sequencing data. In comparison to lrDNA-seq, which typically has read lengths of 10-100k for phasing, lrRNA-seq can still achieve read lengths of 1-3k, allowing for phasing in specific regions. However, to date, no methods have investigated the impact of phasing on the performance of lrRNA-seq. In this study, we attempted to incorporate additional phasing information into Clair3-RNA architecture to gain a deeper understanding of its contribution to lrRNA-seq performance.

#### Variant phasing and read haplotagging

As illustrated in Figure 6a, during the training phase, we selected only the heterozygous germline SNP variants identified in both Clair3-RNA and the GIAB truths for variant phasing. This approach was chosen to mitigate the potential introduction of false positives, which could adversely affect phasing accuracy and lead to cumulative phasing errors during model training. Incorporating only GIAB truths might introduce variants that have insufficient read coverage and discrepancies in zygosity. Alignments are then haplotagged using the phased variants using the WhatsHap “haplotag” submodule, which assigns each phased read the "HP:x" tag, where x indicates the haplotype assigned.

During model training, read subsampling is adopted. We first applied variant phasing and read haplotagging to the BAM file with a full-coverage spectrum. The phased full-coverage reads were then subsampled into varying coverage levels for data augmentation. For candidates with phased reads, we randomly switched the phased reads to unphased reads by discarding the “HP” tag. This switching process simulates phasing scenarios for low-coverage BAM files, where the read haplotagging percentage is lower, while avoiding introducing incorrect phasing that could occur when applying phasing to each subsampled coverage. All subsampled candidates and switched candidates were generated with additional phasing features for model training. In the inference phase, the heterozygous variants were acquired from Clair3-RNA’s first round called variants. We enabled the "--distrust-genotypes" option in WhatsHap phasing to allow for switching variant zygosities in the event of incorrectly called zygosities.

#### Pileup phasing feature

We incorporated 12 pileup phasing features for each candidate position, represented as *X*_H1_ and *X*_H2_, where *X*∈{A, C, G, T, I, D}. The phasing features denote the total counts of nucleotides A, C, G, and T, as well as insertions (I) and deletions (D) for each candidate site in the phased paternal and maternal haplotypes, derived from the "HP" tag of the phased reads. The 12 additional pileup phasing features were concatenated with the original 18 pileup features, resulting in a total of 30 features that were input into the neural network for model training and inference.

### Analysis of overlap and error statistics for GENCODE annotated gene

The gene list was obtained from the GENCODE database (version 46), focusing specifically on the "gene" category for analysis. We categorized the genes based on their overlap with callable high-confidence regions, defined as the intersection of the callable region with read support and the GIAB high-confidence regions. Giving each gene coordinate and all its overlapping high-confidence regions (*hc*), we calculate the intersection size using the formula:

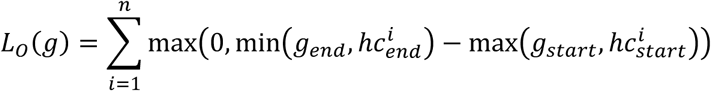

Where *n* represents the number of overlapping regions. *g* and *hc* refer to gene and callable high-confidence region, respectively. The overlap ratio score *H*(*g*) for each gene is then computed as follows:

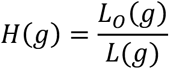

Where *L*(*g*) denotes the gene size, a gene is categorized as “wholly covered” if *H*(*g*) equals 1. Conversely, it is classified as “partially covered” if the total overlap length *H*(*g*) is greater than or equal to *α*, where *α* is a threshold percentage for partial overlapping and is manually set to 0.75. If *H*(*g*) equals 0, the gene is classified as “not covered”.

A gene is considered to have no SNP or Indel errors if true positives are present within the specified gene coordinates, with no false positives or false negatives in the same region, based on benchmarking results obtained from hap.py. A gene is considered to have no switch errors if all heterozygous variants in the given phasesets are phased correctly. Switch errors [30] were assessed using the WhatsHap “compare” submodule within the given overlapping phasesets of lrDNA-seq and lrRNA-seq. The editing distance of the phased heterozygous variants in these two groups was computed as switch errors. The command used for this analysis is given in the **Supplementary Notes** – **Command line used** section.

### Data preparation

#### PacBio mapping with pbmm2 and minimap2

PacBio Iso-Seq and MAS-Seq reads were aligned with pbmm2 or minimap2 with the configurations provided in the **Supplementary Notes**. ONT reads were aligned with minimap2 splice mode. As discussed in the **Results − Performance on PacBio** section, we found miniamp2 alignment output performed in RNA data alignment, even though pbmm2 was optimized for PacBio data.

#### ONT dRNA004 base-calling and mapping

The ONT dRNA002 and cDNA reads were basecalled using Guppy (version 6.4.6), and the ONT dRNA004 reads were basecalled using Dorado (version 3.2.4). Basecalled reads with low read quality scores (“qs” < 10) were labelled as “FAILED” and discarded, as recommended by the ONT protocol. Subsequently, the basecalled reads were mapped to the GRCh38 human reference genome using minimap2 for evaluation.

#### ONT SQK-RNA004 kit library preparation and sequencing

The RNA sequencing libraries were constructed following the ONT’s SQK-RNA004 protocol. RNA sourced from GIAB HG002, HG004, and HG005 samples was obtained from the Coriell Institute. The total RNA underwent assessment for concentration, purity, and integrity using Nanodrop and Qubit analyses. Subsequently, the RNA fragments were ligated with a sequencing adapter utilizing ONT’s latest SQK-RNA004 direct RNA sequencing kit. Finally, the RNAs were sequenced on the PromethION flow cell using our in-house PromethION 2 Solo sequencing device, running MinKNOW software version 1.18.02 for 96 hours.

#### Model training and evaluation

Model training was conducted using the known variants in the GIAB HG002 dataset, encompassing the full spectrum of read coverage, along with four subsampled coverage levels. For all pre-trained models described in the manuscript, we consistently used variants from chromosomes 1 to 19, 21, and 22 for training, with a random selection of 90% of the data for training and the remaining 10% for evaluation, while always holding out chromosome 20 in training. Performance evaluation was conducted in callable regions defined in the **Methods** - **Benchmarking metrics** section. Clair3-RNA trains the model for either 30 epochs or until the validation loss does not decrease for five consecutive epochs, employing an exponentially decaying learning rate until convergence. The epoch that performs best on the validation set will be selected for benchmarking and as the production model. The training process utilizes the Rectified Adam optimizer [34] and includes an initial weight warm-up. An optimized cross-entropy loss function, specifically Focal Loss [32], is used during model training to enhance performance.

### Benchmarking metrics

RNA sequences represent transcribed segments of the genome, encompassing mainly the exon regions. Notably, the number of total variants in RNA was considerably lower than that in DNA, where there were ∼27,000 to ∼318,000 variants, depending on the sequencing platform, RNA throughput, and specific gene expression. In contrast, ∼4 million true variants were available for DNA data. Figure 3a shows the benchmarking workflow for RNA variant evaluation. We utilized Clair3-RNA’s "get_rna_bed" submodule to identify callable regions with minimum read support (default set at 4, adjustable through options). These callable regions were then cross-referenced with the GIAB high-confidence regions to establish reliable benchmarking regions. Within the benchmark VCF from GIAB, variants with less than two reads supporting the alternate allele were excluded. We used hap.py [35] to compare the variant-called variants against the truth variants in callable high-confidence regions. We used Clair3-RNA’s "calculate_overall_metrics" submodule to generate three metrics, "Precision", "Recall", and "F1-score", for five categories: "Overall", "SNP", "Indel", "Insertion", and "Deletion".

### Computational performance

Clair3-RNA was implemented with Python and TensorFlow and leveraged PyPy for speedup. RNA variant calling using Clair3-RNA requires only CPU resources. Using one replicate sequencing data, Clair3-RNA takes ∼30 minutes for ONT dRNA004 or PacBio MAS-Seq, using two 12-core Intel Xeon Silver 4116 processors. The memory consumption was capped at 500 MB per process. Model training requires a high-end graphics processing unit (GPU). We used TensorFlow to train models on an Nvidia GeForce RTX™ 3090 Ti and 4090 Ti GPU. We used the GNU time command to benchmark the runtime and memory usage.

## Code availability

Clair3-RNA is open source and available at https://github.com/HKU-BAL/Clair3-RNA under the BSD 3-Clause license. A public Docker image <u>hkubal/clair3-rna:latest</u> that can evaluate Clair3-RNA’s evaluation pipeline is also available for simplicity. Users can run the workflow seamlessly in one command with the input RNA BAM file and reference file. Clair3-RNA provides user interface flexibility that supports parallel computing, user-customized parameter setting, and workflow refinement.

## Data availability

The links to the reference genomes, truth variants, benchmarking materials, and ONT & PacBio dataset, and the benchmarked commands and parameters used in this study are available in the **Supplementary Notes**. All analysis output, including the VCFs and running logs, are available at http://www.bio8.cs.hku.hk/clair3_rna/analysis_result. The ONT dRNA004 GIAB HG002, HG004, and HG005 sequencing data using the dRNA SQK-RNA004 kit generated in this study was deposited in the NCBI archive with accession ID PRJNA1169852.

## Acknowledgements

We would like to thank the GIAB community for providing us an early access to the lrRNA-seq datasets. R.L. was supported by Hong Kong Research Grants Council grants GRF (17113721) and TRS (T21-705/20-N), the Shenzhen Municipal Government General Program (JCYJ20210324134405015), the URC fund at HKU and Oxford Nanopore Technologies. R.K.K. and F.S. are supported in part by NIH grant (1U01HG011758-01).

## Author contributions

R.L. and F.S. conceived the study. Z.Z., X.Y., and R.L. designed the algorithms, implemented Clair3-RNA, and wrote the paper. F.S. and J.M. supported datasets for benchmarking and evaluation. Y.L., C.L., X.C., R.K.K., F.S., and J.M. evaluated the benchmarking results. All authors revised the manuscript.

