## Supplemental Notes for "Clair3-RNA: A deep learning-based small variant caller for long-read RNA sequencing data"

### Supplementary Notes

#### Supplementary Figures

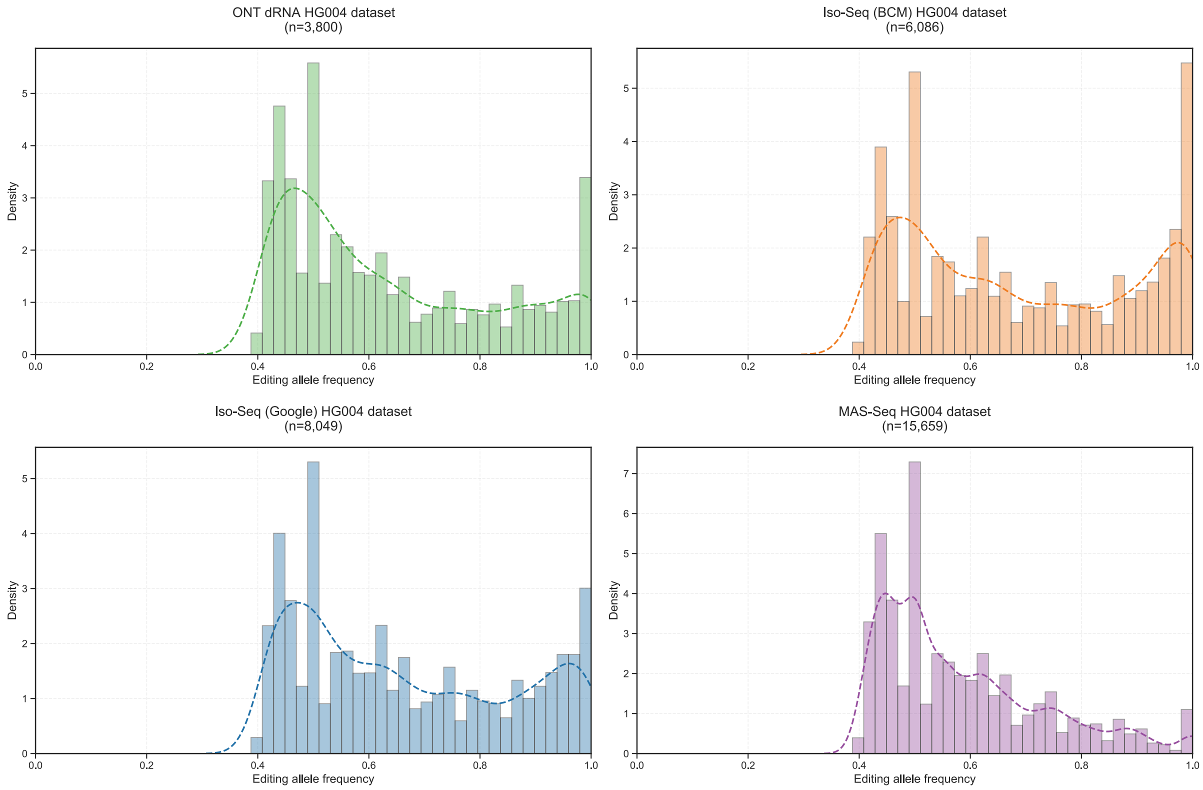


**Supplementary Figure 1. Allele frequency distribution of benchmark RNA editing sites in various datasets.**

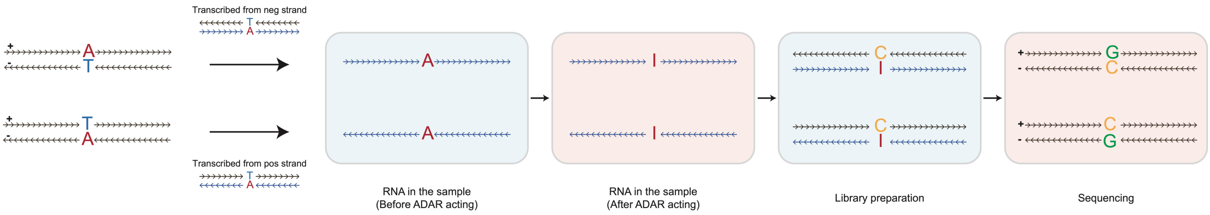


**Supplementary Figure 2. Mechanism of RNA transcription in single-stranded or double-stranded contexts.**

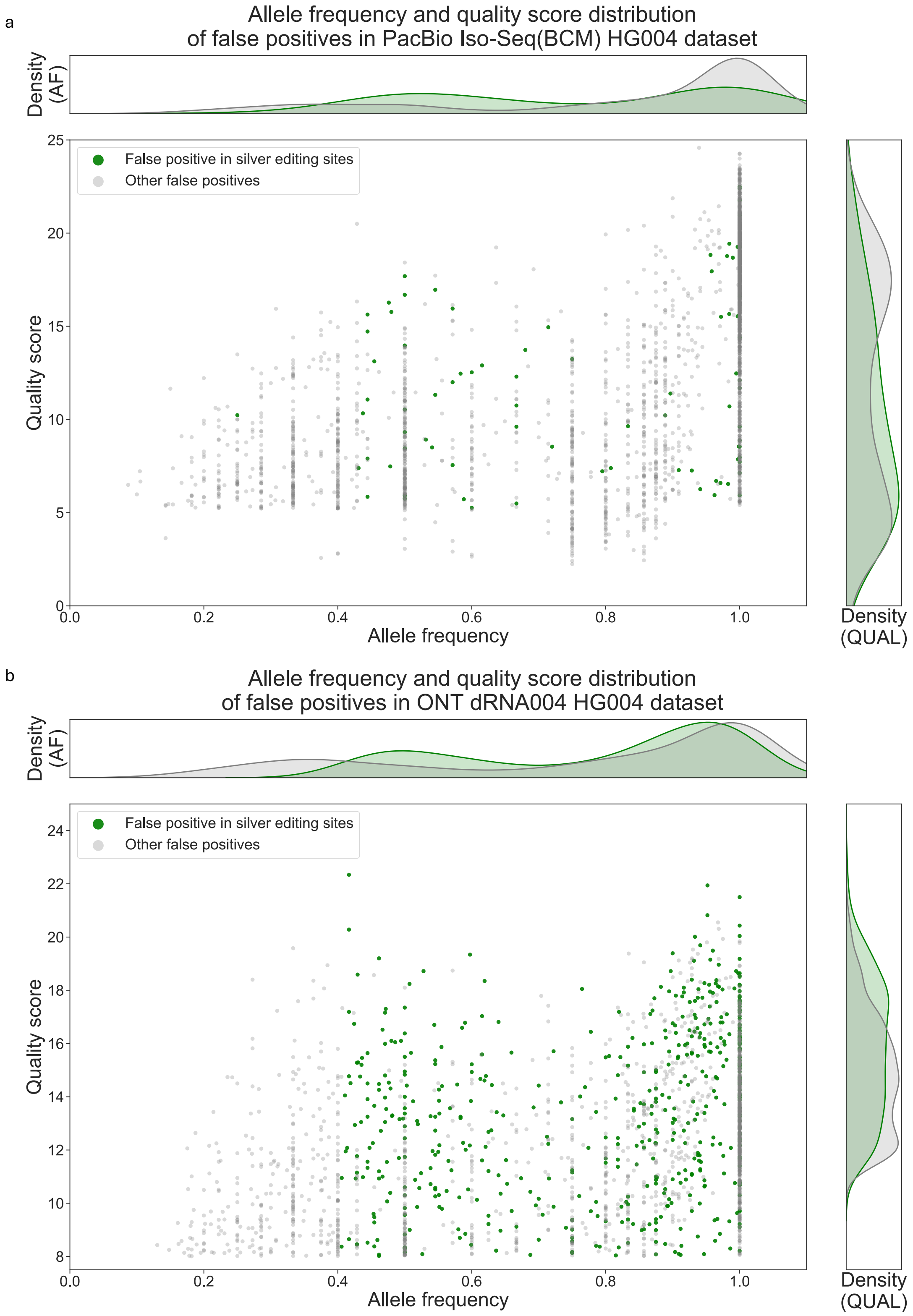


**Supplementary Figure 3. The quality score and allele frequency distribution of true variant in different datasets.**

(a) The allele frequency and quality score distribution of PacBio Iso-Seq HG004 dataset. (b) The distribution of ONT dRNA004 HG004 dataset.

### Supplementary Tables

**Supplementary Table 1. Summary of the datasets used for model training and performance evaluation.**

**BCM**: Baylor College of Medicine; **HKU**: The University of Hong Kong; **NEU**: Northeastern University.

| Platform | Chemistry /Instruments/ Basecaller | Sequen-cing type | Sample | Source | Data size | Average error rate | Average read length | Used in training | Used for testing |
|---|---|---|---|---|---|---|---|---|---|
| PacBio | Iso-Seq | cDNA | HG002 | BCM | 15G | 1.1% | 2,875 | ✓ | |
| | | | HG004 | BCM | 4.1G | 0.6% | 2,499 | | ✓ |
| | | | HG005 | BCM | 5.7G | 0.8% | 2,910 | | ✓ |
| | | | HG002 | Google | 11G | 3.7% | 2,807 | ✓ | |
| | | | HG004 | Google | 3.3G | 2.8% | 2,908 | | ✓ |
| | | | HG005 | Google | 3.2G | 2.9% | 2,752 | | ✓ |
| | MAS-Seq | | HG002 | BCM | 93G | 0.8% | 1,680 | ✓ | |
| | | | HG004 | BCM | 29G | 0.4% | 1,731 | | ✓ |
| ONT | SQK-RNA004 kit, Dorado | dRNA | HG002 | HKU | 54G | 1.7% | 1,176 | ✓ | |
| | | | HG004 | HKU | 36G | 1.8% | 1,404 | | ✓ |
| | | | HG005 | HKU | 37G | 1.8% | 1,180 | | ✓ |
| | SQK-RNA002 kit, Guppy | | HG002 | NEU | 14G | 10.2% | 1,106 | ✓ | |
| | | | HG004 | NEU | 3.3G | 9.7% | 969 | | ✓ |
| | | | HG005 | NEU | 2.9G | 9.6% | 1,020 | | ✓ |
| | R9.4.1, Guppy | cDNA | HG002 | NEU | 127G | 4.7% | 1,137 | ✓ | |
| | | | HG004 | NEU | 44G | 4.3% | 1,154 | | ✓ |
| | | | HG005 | NEU | 61G | 4.2% | 1,239 | | ✓ |

**Supplementary Table 2. ONT performance of different callers.**

(a) Performance on ONT dRNA004

| Read Coverage | Allele depth | Dataset | Sample | Caller | Regarding zygosity | | | | | | Disregarding zygosity | | | | | |
|---|---|---|---|---|---|---|---|---|---|---|---|---|---|---|---|---|
| | | | | | SNP | | | Indel | | | SNP | | | Indel | | |
| | | | | | Precision | Recall | F1-score | Precision | Recall | F1-score | Precision | Recall | F1-score | Precision | Recall | F1-score |
| DP≥4 | AD≥2 | dRNA004 | HG004 | Clair3-RNA | 92.82% | 89.26% | 91.00% | 83.61% | 47.19% | 60.33% | 95.63% | 91.97% | 93.76% | 89.39% | 50.46% | 64.50% |
| | | | HG004 | LongcallR | 94.49% | 45.11% | 61.07% | \ | \ | \ | 95.59% | 45.63% | 61.77% | \ | \ | \ |
| | | | HG004 | Clair3 | 66.93% | 84.89% | 74.85% | 41.23% | 45.32% | 43.18% | 69.38% | 87.99% | 77.59% | 45.98% | 50.63% | 48.19% |
| | | | HG004 | DeepVariant | 86.53% | 43.55% | 57.94% | 46.60% | 26.79% | 34.02% | 92.89% | 46.75% | 62.19% | 52.97% | 30.52% | 38.73% |
| | | | HG005 | Clair3-RNA | 93.16% | 90.34% | 91.73% | 85.64% | 53.76% | 66.05% | 95.30% | 92.42% | 93.84% | 90.52% | 56.85% | 69.84% |
| | | | HG005 | LongcallR | 94.89% | 49.67% | 65.20% | \ | \ | \ | 95.76% | 50.12% | 65.80% | \ | \ | \ |
| | | | HG005 | Clair3 | 70.33% | 85.58% | 77.21% | 50.96% | 49.50% | 50.22% | 72.14% | 87.78% | 79.20% | 55.45% | 53.96% | 54.70% |
| | | | HG005 | DeepVariant | 87.14% | 42.31% | 56.96% | 54.85% | 29.04% | 37.98% | 93.20% | 45.25% | 60.92% | 62.01% | 32.91% | 43.00% |
| DP≥8 | AD≥2 | dRNA004 | HG004 | Clair3-RNA | 94.42% | 92.49% | 93.45% | 84.02% | 47.85% | 60.98% | 96.18% | 94.21% | 95.19% | 88.59% | 50.46% | 64.30% |
| | | | HG004 | LongcallR | 93.98% | 54.95% | 69.35% | \ | \ | \ | 94.96% | 55.53% | 70.08% | \ | \ | \ |
| | | | HG004 | Clair3 | 70.46% | 88.40% | 78.41% | 41.84% | 45.66% | 43.67% | 72.24% | 90.64% | 80.40% | 46.19% | 50.53% | 48.26% |
| | | | HG004 | DeepVariant | 89.60% | 51.67% | 65.54% | 46.23% | 27.93% | 34.82% | 93.28% | 53.79% | 68.24% | 51.14% | 30.96% | 38.57% |
| | | | HG005 | Clair3-RNA | 94.63% | 93.39% | 94.01% | 85.92% | 54.85% | 66.95% | 95.78% | 94.52% | 95.15% | 89.90% | 57.42% | 70.08% |
| | | | HG005 | LongcallR | 94.11% | 58.30% | 71.99% | \ | \ | \ | 94.90% | 58.78% | 72.59% | \ | \ | \ |
| | | | HG005 | Clair3 | 73.59% | 90.55% | 81.20% | 52.58% | 51.23% | 51.90% | 74.61% | 91.80% | 82.31% | 56.55% | 55.20% | 55.86% |
| | | | HG005 | DeepVariant | 90.20% | 49.67% | 64.07% | 55.30% | 30.30% | 39.15% | 93.44% | 51.46% | 66.37% | 60.90% | 33.45% | 43.18% |
| DP≥10 | AD≥2 | dRNA004 | HG004 | Clair3-RNA | 94.75% | 93.28% | 94.01% | 84.34% | 47.80% | 61.02% | 96.37% | 94.87% | 95.62% | 88.57% | 50.21% | 64.09% |
| | | | HG004 | LongcallR | 93.74% | 58.27% | 71.87% | \ | \ | \ | 94.68% | 58.86% | 72.59% | \ | \ | \ |
| | | | HG004 | Clair3 | 72.88% | 88.44% | 79.91% | 42.55% | 45.24% | 43.85% | 74.65% | 90.59% | 81.85% | 46.94% | 50.04% | 48.44% |
| | | | HG004 | DeepVariant | 90.29% | 54.54% | 68.01% | 45.75% | 28.51% | 35.13% | 93.56% | 56.52% | 70.47% | 50.29% | 31.40% | 38.66% |
| | | | HG005 | Clair3-RNA | 94.85% | 94.20% | 94.52% | 86.07% | 55.00% | 67.12% | 95.87% | 95.21% | 95.54% | 89.78% | 57.40% | 70.03% |
| | | | HG005 | LongcallR | 93.81% | 61.15% | 74.04% | \ | \ | \ | 94.54% | 61.62% | 74.61% | \ | \ | \ |
| | | | HG005 | Clair3 | 75.88% | 90.82% | 82.68% | 53.86% | 51.19% | 52.49% | 76.84% | 91.96% | 83.72% | 57.79% | 55.03% | 56.37% |
| | | | HG005 | DeepVariant | 90.68% | 52.33% | 66.37% | 55.12% | 30.94% | 39.64% | 93.54% | 53.98% | 68.46% | 60.26% | 33.90% | 43.39% |
| DP≥10 | AD≥4 | dRNA004 | HG004 | Clair3-RNA | 94.98% | 94.51% | 94.75% | 84.86% | 50.57% | 63.37% | 96.62% | 96.14% | 96.38% | 89.04% | 53.07% | 66.50% |
| | | | HG004 | LongcallR | 93.69% | 59.02% | 72.42% | \ | \ | \ | 94.63% | 59.61% | 73.14% | \ | \ | \ |
| | | | HG004 | Clair3 | 81.88% | 89.35% | 85.45% | 48.25% | 47.60% | 47.92% | 83.88% | 91.53% | 87.54% | 52.94% | 52.37% | 52.65% |
| | | | HG004 | DeepVariant | 90.53% | 55.61% | 68.90% | 46.23% | 30.06% | 36.43% | 93.81% | 57.63% | 71.40% | 50.74% | 33.05% | 40.03% |
| | | | HG005 | Clair3-RNA | 95.06% | 95.28% | 95.17% | 86.69% | 58.02% | 69.51% | 96.10% | 96.31% | 96.21% | 90.23% | 60.41% | 72.37% |
| | | | HG005 | LongcallR | 93.75% | 61.70% | 74.42% | \ | \ | \ | 94.48% | 62.17% | 74.99% | \ | \ | \ |
| | | | HG005 | Clair3 | 84.06% | 91.64% | 87.68% | 59.69% | 53.75% | 56.56% | 85.12% | 92.80% | 88.80% | 63.93% | 57.67% | 60.64% |
| | | | HG005 | DeepVariant | 90.80% | 53.17% | 67.07% | 55.59% | 32.61% | 41.11% | 93.68% | 54.85% | 69.19% | 60.72% | 35.67% | 44.94% |

(b) Performance on ONT cDNA

| Read Coverage | Allele depth | Dataset | Sample | Caller | Regarding zygosity | | | | | | Disregarding zygosity | | | | | |
|---|---|---|---|---|---|---|---|---|---|---|---|---|---|---|---|---|
| | | | | | SNP | | | Indel | | | SNP | | | Indel | | |
| | | | | | Precision | Recall | F1-score | Precision | Recall | F1-score | Precision | Recall | F1-score | Precision | Recall | F1-score |
| DP≥4 | AD≥2 | cDNA | HG004 | Clair3-RNA | 69.53% | 67.69% | 68.60% | 45.16% | 36.11% | 40.13% | 80.86% | 78.73% | 79.78% | 54.41% | 43.50% | 48.35% |
| | | | HG004 | LongcallR | 88.17% | 31.26% | 46.16% | \ | \ | \ | 91.60% | 32.48% | 47.95% | \ | \ | \ |
| | | | HG004 | Clair3 | 38.86% | 73.13% | 50.75% | 32.92% | 38.43% | 35.46% | 44.33% | 83.41% | 57.89% | 38.12% | 44.69% | 41.14% |
| | | | HG004 | DeepVariant | 60.34% | 42.69% | 50.00% | 30.93% | 22.64% | 26.14% | 72.18% | 51.07% | 59.82% | 39.41% | 29.06% | 33.46% |
| | | | HG005 | Clair3-RNA | 87.19% | 73.63% | 79.84% | 65.39% | 43.77% | 52.44% | 92.18% | 77.85% | 84.41% | 72.49% | 48.53% | 58.14% |
| | | | HG005 | LongcallR | 94.15% | 31.99% | 47.75% | \ | \ | \ | 96.03% | 32.63% | 48.70% | \ | \ | \ |
| | | | HG005 | Clair3 | 48.48% | 80.97% | 60.65% | 48.39% | 47.41% | 47.89% | 50.72% | 84.71% | 63.45% | 51.86% | 50.96% | 51.41% |
| | | | HG005 | DeepVariant | 76.65% | 35.85% | 48.85% | 45.36% | 22.05% | 29.68% | 84.60% | 39.57% | 53.92% | 52.65% | 25.71% | 34.55% |
| DP≥8 | AD≥2 | cDNA | HG004 | Clair3-RNA | 79.61% | 79.70% | 79.66% | 49.35% | 36.73% | 42.12% | 84.47% | 84.57% | 84.52% | 55.43% | 41.26% | 47.31% |
| | | | HG004 | LongcallR | 88.17% | 53.71% | 66.76% | \ | \ | \ | 91.60% | 55.81% | 69.36% | \ | \ | \ |
| | | | HG004 | Clair3 | 51.02% | 85.90% | 64.02% | 37.02% | 40.29% | 38.58% | 53.85% | 90.67% | 67.57% | 40.51% | 44.24% | 42.30% |
| | | | HG004 | DeepVariant | 75.83% | 54.17% | 63.20% | 39.67% | 22.76% | 28.92% | 81.71% | 58.37% | 68.09% | 46.67% | 26.92% | 34.15% |
| | | | HG005 | Clair3-RNA | 91.05% | 83.46% | 87.09% | 65.13% | 45.03% | 53.25% | 93.19% | 85.43% | 89.14% | 70.67% | 48.88% | 57.79% |
| | | | HG005 | LongcallR | 94.15% | 51.98% | 66.98% | \ | \ | \ | 96.03% | 53.02% | 68.32% | \ | \ | \ |
| | | | HG005 | Clair3 | 59.06% | 91.10% | 71.66% | 48.55% | 50.20% | 49.36% | 60.17% | 92.80% | 73.00% | 51.27% | 53.16% | 52.20% |
| | | | HG005 | DeepVariant | 83.86% | 46.93% | 60.18% | 51.23% | 23.86% | 32.55% | 88.14% | 49.32% | 63.25% | 57.52% | 26.90% | 36.65% |
| DP≥10 | AD≥2 | cDNA | HG004 | Clair3-RNA | 83.39% | 82.24% | 82.81% | 50.99% | 36.49% | 42.54% | 86.91% | 85.71% | 86.31% | 56.59% | 40.50% | 47.21% |
| | | | HG004 | LongcallR | 88.17% | 63.00% | 73.49% | \ | \ | \ | 91.60% | 65.44% | 76.34% | \ | \ | \ |
| | | | HG004 | Clair3 | 56.89% | 87.59% | 68.97% | 39.33% | 39.84% | 39.58% | 59.19% | 91.14% | 71.77% | 42.68% | 43.37% | 43.02% |
| | | | HG004 | DeepVariant | 79.88% | 58.06% | 67.25% | 42.13% | 22.72% | 29.52% | 84.56% | 61.46% | 71.19% | 48.88% | 26.51% | 34.38% |
| | | | HG005 | Clair3-RNA | 92.15% | 85.21% | 88.54% | 65.30% | 45.22% | 53.44% | 93.85% | 86.78% | 90.18% | 70.52% | 48.84% | 57.71% |
| | | | HG005 | LongcallR | 94.16% | 60.49% | 73.66% | \ | \ | \ | 96.03% | 61.69% | 75.12% | \ | \ | \ |
| | | | HG005 | Clair3 | 63.11% | 92.14% | 74.91% | 48.59% | 50.08% | 49.33% | 64.06% | 93.53% | 76.04% | 51.23% | 52.97% | 52.08% |
| | | | HG005 | DeepVariant | 85.71% | 50.95% | 63.91% | 52.40% | 24.55% | 33.43% | 89.31% | 53.09% | 66.60% | 58.22% | 27.38% | 37.25% |
| DP≥10 | AD≥4 | cDNA | HG004 | Clair3-RNA | 85.32% | 84.76% | 85.04% | 53.68% | 39.97% | 45.82% | 88.94% | 88.36% | 88.65% | 59.31% | 44.16% | 50.63% |
| | | | HG004 | LongcallR | 88.16% | 65.43% | 75.11% | \ | \ | \ | 91.59% | 67.97% | 78.03% | \ | \ | \ |
| | | | HG004 | Clair3 | 65.43% | 89.33% | 75.54% | 41.41% | 42.88% | 42.13% | 68.13% | 93.00% | 78.64% | 44.78% | 46.52% | 45.63% |
| | | | HG004 | DeepVariant | 82.53% | 60.22% | 69.64% | 43.85% | 25.00% | 31.84% | 87.38% | 63.76% | 73.72% | 50.53% | 28.96% | 36.82% |
| | | | HG005 | Clair3-RNA | 93.16% | 87.45% | 90.21% | 67.75% | 48.73% | 56.69% | 94.90% | 89.07% | 91.89% | 72.75% | 52.34% | 60.88% |
| | | | HG005 | LongcallR | 94.15% | 62.53% | 75.15% | \ | \ | \ | 96.02% | 63.77% | 76.64% | \ | \ | \ |
| | | | HG005 | Clair3 | 70.02% | 93.61% | 80.11% | 51.29% | 53.43% | 52.34% | 71.08% | 95.03% | 81.33% | 53.87% | 56.26% | 55.04% |
| | | | HG005 | DeepVariant | 86.80% | 52.71% | 65.59% | 53.47% | 26.60% | 35.53% | 90.44% | 54.93% | 68.35% | 59.22% | 29.57% | 39.45% |

(c) Performance on ONT dRNA002

| Read Coverage | Allele depth | Dataset | Sample | Caller | Regarding zygosity | | | | | | Disregarding zygosity | | | | | |
|---|---|---|---|---|---|---|---|---|---|---|---|---|---|---|---|---|
| | | | | | SNP | | | Indel | | | SNP | | | Indel | | |
| | | | | | Precision | Recall | F1-score | Precision | Recall | F1-score | Precision | Recall | F1-score | Precision | Recall | F1-score |
| DP≥4 | AD≥2 | dRNA002 | HG004 | Clair3-RNA | 66.14% | 71.27% | 68.61% | 36.34% | 14.93% | 21.16% | 68.33% | 73.63% | 70.88% | 40.02% | 16.44% | 23.31% |
| | | | HG004 | LongcallR | 80.85% | 32.44% | 46.31% | \ | \ | \ | 84.15% | 33.77% | 48.20% | \ | \ | \ |
| | | | HG004 | Clair3 | 16.30% | 73.82% | 26.70% | 1.64% | 31.74% | 3.12% | 17.26% | 78.19% | 28.28% | 1.79% | 34.74% | 3.41% |
| | | | HG004 | DeepVariant | 65.78% | 42.07% | 51.32% | 1.54% | 18.55% | 2.85% | 76.17% | 48.71% | 59.42% | 1.87% | 22.56% | 3.46% |
| | | | HG005 | Clair3-RNA | 66.57% | 73.75% | 69.98% | 37.14% | 18.18% | 24.41% | 68.28% | 75.65% | 71.77% | 39.09% | 19.13% | 25.69% |
| | | | HG005 | LongcallR | 81.03% | 33.36% | 47.26% | \ | \ | \ | 84.18% | 34.65% | 49.10% | \ | \ | \ |
| | | | HG005 | Clair3 | 16.86% | 76.57% | 27.63% | 1.83% | 40.54% | 3.50% | 17.54% | 79.67% | 28.75% | 1.96% | 43.40% | 3.74% |
| | | | HG005 | DeepVariant | 69.02% | 45.54% | 54.87% | 1.65% | 21.87% | 3.07% | 77.17% | 50.91% | 61.35% | 1.97% | 26.14% | 3.66% |
| DP≥8 | AD≥2 | dRNA002 | HG004 | Clair3-RNA | 66.52% | 78.40% | 71.97% | 36.02% | 15.08% | 21.26% | 67.84% | 79.96% | 73.40% | 39.79% | 16.66% | 23.48% |
| | | | HG004 | LongcallR | 80.85% | 42.85% | 56.01% | \ | \ | \ | 84.15% | 44.60% | 58.30% | \ | \ | \ |
| | | | HG004 | Clair3 | 16.45% | 76.98% | 27.10% | 1.53% | 31.04% | 2.91% | 17.06% | 79.82% | 28.11% | 1.67% | 34.01% | 3.19% |
| | | | HG004 | DeepVariant | 67.18% | 46.27% | 54.80% | 1.46% | 18.15% | 2.71% | 76.66% | 52.80% | 62.53% | 1.75% | 21.77% | 3.24% |
| | | | HG005 | Clair3-RNA | 66.80% | 79.95% | 72.78% | 36.88% | 18.01% | 24.20% | 67.83% | 81.18% | 73.90% | 38.96% | 19.02% | 25.56% |
| | | | HG005 | LongcallR | 81.03% | 43.63% | 56.72% | \ | \ | \ | 84.18% | 45.33% | 58.92% | \ | \ | \ |
| | | | HG005 | Clair3 | 17.11% | 79.53% | 28.16% | 1.66% | 39.42% | 3.19% | 17.54% | 81.54% | 28.87% | 1.78% | 42.21% | 3.42% |
| | | | HG005 | DeepVariant | 70.78% | 49.68% | 58.38% | 1.54% | 21.42% | 2.88% | 78.05% | 54.79% | 64.38% | 1.83% | 25.46% | 3.41% |
| DP≥10 | AD≥2 | dRNA002 | HG004 | Clair3-RNA | 66.90% | 80.03% | 72.88% | 35.37% | 14.91% | 20.98% | 68.04% | 81.39% | 74.12% | 39.16% | 16.51% | 23.23% |
| | | | HG004 | LongcallR | 80.85% | 47.85% | 60.12% | \ | \ | \ | 84.15% | 49.80% | 62.57% | \ | \ | \ |
| | | | HG004 | Clair3 | 17.00% | 77.24% | 27.87% | 1.54% | 30.72% | 2.94% | 17.56% | 79.79% | 28.78% | 1.68% | 33.51% | 3.20% |
| | | | HG004 | DeepVariant | 67.67% | 47.17% | 55.59% | 1.46% | 18.16% | 2.70% | 77.10% | 53.74% | 63.33% | 1.74% | 21.74% | 3.22% |
| | | | HG005 | Clair3-RNA | 67.32% | 81.47% | 73.72% | 35.89% | 17.43% | 23.47% | 68.23% | 82.57% | 74.71% | 37.95% | 18.43% | 24.81% |
| | | | HG005 | LongcallR | 81.03% | 48.73% | 60.86% | \ | \ | \ | 84.18% | 50.62% | 63.23% | \ | \ | \ |
| | | | HG005 | Clair3 | 17.76% | 80.06% | 29.07% | 1.66% | 38.66% | 3.19% | 18.15% | 81.84% | 29.71% | 1.78% | 41.47% | 3.42% |
| | | | HG005 | DeepVariant | 71.37% | 50.52% | 59.16% | 1.50% | 20.88% | 2.80% | 78.47% | 55.55% | 65.05% | 1.79% | 24.95% | 3.33% |
| DP≥10 | AD≥4 | dRNA002 | HG004 | Clair3-RNA | 67.85% | 84.01% | 75.07% | 37.07% | 16.82% | 23.14% | 69.00% | 85.43% | 76.34% | 40.91% | 18.56% | 25.53% |
| | | | HG004 | LongcallR | 80.85% | 50.80% | 62.40% | \ | \ | \ | 84.15% | 52.88% | 64.94% | \ | \ | \ |
| | | | HG004 | Clair3 | 22.68% | 80.09% | 35.36% | 1.59% | 34.17% | 3.04% | 23.43% | 82.71% | 36.51% | 1.72% | 36.95% | 3.28% |
| | | | HG004 | DeepVariant | 67.78% | 50.16% | 57.66% | 1.47% | 20.49% | 2.74% | 77.24% | 57.16% | 65.70% | 1.75% | 24.48% | 3.26% |
| | | | HG005 | Clair3-RNA | 68.21% | 84.70% | 75.57% | 36.78% | 19.70% | 25.65% | 69.14% | 85.86% | 76.60% | 38.90% | 20.84% | 27.14% |
| | | | HG005 | LongcallR | 81.04% | 51.29% | 62.82% | \ | \ | \ | 84.18% | 53.27% | 65.25% | \ | \ | \ |
| | | | HG005 | Clair3 | 23.62% | 82.33% | 36.70% | 1.69% | 42.89% | 3.24% | 24.13% | 84.13% | 37.50% | 1.79% | 45.51% | 3.44% |
| | | | HG005 | DeepVariant | 71.48% | 53.29% | 61.06% | 1.51% | 23.93% | 2.84% | 78.58% | 58.58% | 67.12% | 1.79% | 28.45% | 3.36% |

**Supplementary Table 3. PacBio performance of different callers.**

(a) Performance on Iso-Seq (BCM)

| Read Coverage | Allele depth | Dataset | Sample | Caller | Regarding zygosity | | | | | | Disregarding zygosity | | | | | |
|---|---|---|---|---|---|---|---|---|---|---|---|---|---|---|---|---|
| | | | | | SNP | | | Indel | | | SNP | | | Indel | | |
| | | | | | Precision | Recall | F1-score | Precision | Recall | F1-score | Precision | Recall | F1-score | Precision | Recall | F1-score |
| DP≥4 | AD≥2 | Iso-Seq (BCM) | HG004 | Clair3-RNA | 96.16% | 93.73% | 94.93% | 80.74% | 70.21% | 75.11% | 98.89% | 96.39% | 97.62% | 90.96% | 79.13% | 84.64% |
| | | | HG004 | LongcallR | 94.51% | 64.67% | 76.79% | \ | \ | \ | 95.89% | 65.61% | 77.92% | \ | \ | \ |
| | | | HG004 | Clair3 | 68.73% | 96.22% | 80.18% | 57.17% | 69.35% | 62.67% | 70.70% | 98.98% | 82.48% | 62.75% | 76.28% | 68.86% |
| | | | HG004 | DeepVariant | 76.03% | 43.18% | 55.08% | 55.63% | 39.68% | 46.32% | 91.80% | 52.14% | 66.50% | 69.77% | 49.95% | 58.22% |
| | | | HG005 | Clair3-RNA | 96.48% | 94.53% | 95.50% | 85.47% | 77.69% | 81.40% | 98.49% | 96.49% | 97.48% | 93.45% | 84.98% | 89.02% |
| | | | HG005 | LongcallR | 94.45% | 67.80% | 78.93% | \ | \ | \ | 95.60% | 68.61% | 79.89% | \ | \ | \ |
| | | | HG005 | Clair3 | 66.80% | 97.15% | 79.16% | 64.31% | 76.18% | 69.75% | 68.21% | 99.21% | 80.84% | 68.83% | 81.63% | 74.68% |
| | | | HG005 | DeepVariant | 72.74% | 39.83% | 51.48% | 63.80% | 39.08% | 48.47% | 90.96% | 49.80% | 64.37% | 77.51% | 47.61% | 58.99% |
| DP≥8 | AD≥2 | Iso-Seq (BCM) | HG004 | Clair3-RNA | 98.19% | 95.30% | 96.72% | 83.56% | 73.68% | 78.31% | 99.13% | 96.22% | 97.66% | 91.35% | 80.58% | 85.62% |
| | | | HG004 | LongcallR | 94.35% | 90.74% | 92.51% | \ | \ | \ | 95.24% | 91.59% | 93.38% | \ | \ | \ |
| | | | HG004 | Clair3 | 68.48% | 97.48% | 80.45% | 51.50% | 73.12% | 60.43% | 69.25% | 98.57% | 81.35% | 56.15% | 79.96% | 65.97% |
| | | | HG004 | DeepVariant | 82.58% | 50.63% | 62.77% | 60.99% | 38.79% | 47.42% | 93.03% | 57.03% | 70.71% | 70.29% | 44.89% | 54.79% |
| | | | HG005 | Clair3-RNA | 98.22% | 95.84% | 97.01% | 88.49% | 81.53% | 84.87% | 98.86% | 96.46% | 97.65% | 94.08% | 86.71% | 90.24% |
| | | | HG005 | LongcallR | 94.32% | 89.83% | 92.02% | \ | \ | \ | 95.02% | 90.49% | 92.70% | \ | \ | \ |
| | | | HG005 | Clair3 | 67.26% | 98.38% | 79.90% | 60.57% | 80.29% | 69.05% | 67.78% | 99.13% | 80.51% | 64.25% | 85.30% | 73.30% |
| | | | HG005 | DeepVariant | 77.66% | 44.27% | 56.39% | 70.00% | 37.25% | 48.62% | 91.97% | 52.42% | 66.78% | 78.59% | 41.94% | 54.69% |
| DP≥10 | AD≥2 | Iso-Seq (BCM) | HG004 | Clair3-RNA | 98.49% | 95.37% | 96.90% | 84.42% | 74.48% | 79.14% | 99.27% | 96.13% | 97.67% | 91.63% | 80.89% | 85.93% |
| | | | HG004 | LongcallR | 93.81% | 91.22% | 92.49% | \ | \ | \ | 94.48% | 91.86% | 93.16% | \ | \ | \ |
| | | | HG004 | Clair3 | 68.74% | 97.39% | 80.59% | 49.51% | 73.44% | 59.15% | 69.44% | 98.38% | 81.41% | 54.08% | 80.51% | 64.69% |
| | | | HG004 | DeepVariant | 83.89% | 53.55% | 65.38% | 62.93% | 38.78% | 47.99% | 93.32% | 59.57% | 72.72% | 70.66% | 43.74% | 54.03% |
| | | | HG005 | Clair3-RNA | 98.58% | 96.06% | 97.30% | 89.30% | 82.49% | 85.76% | 99.07% | 96.53% | 97.78% | 94.38% | 87.22% | 90.66% |
| | | | HG005 | LongcallR | 93.74% | 90.31% | 92.00% | \ | \ | \ | 94.30% | 90.84% | 92.54% | \ | \ | \ |
| | | | HG005 | Clair3 | 68.09% | 98.41% | 80.49% | 59.51% | 81.16% | 68.67% | 68.53% | 99.04% | 81.01% | 63.02% | 86.10% | 72.78% |
| | | | HG005 | DeepVariant | 78.83% | 46.55% | 58.53% | 71.76% | 36.84% | 48.69% | 92.52% | 54.63% | 68.70% | 79.14% | 40.75% | 53.80% |
| DP≥10 | AD≥4 | Iso-Seq (BCM) | HG004 | Clair3-RNA | 98.71% | 96.08% | 97.38% | 86.55% | 79.41% | 82.83% | 99.51% | 96.85% | 98.16% | 93.40% | 85.75% | 89.42% |
| | | | HG004 | LongcallR | 93.68% | 92.00% | 92.84% | \ | \ | \ | 94.37% | 92.67% | 93.51% | \ | \ | \ |
| | | | HG004 | Clair3 | 75.43% | 97.66% | 85.12% | 52.27% | 77.39% | 62.39% | 76.16% | 98.60% | 85.94% | 56.70% | 84.25% | 67.79% |
| | | | HG004 | DeepVariant | 84.61% | 54.42% | 66.24% | 70.29% | 40.94% | 51.74% | 94.23% | 60.61% | 73.77% | 77.45% | 45.29% | 57.16% |
| | | | HG005 | Clair3-RNA | 98.91% | 96.71% | 97.80% | 90.67% | 85.95% | 88.25% | 99.41% | 97.20% | 98.29% | 95.43% | 90.49% | 92.90% |
| | | | HG005 | LongcallR | 93.65% | 91.08% | 92.34% | \ | \ | \ | 94.21% | 91.61% | 92.90% | \ | \ | \ |
| | | | HG005 | Clair3 | 76.27% | 98.67% | 86.03% | 62.18% | 84.05% | 71.48% | 76.72% | 99.25% | 86.54% | 65.58% | 88.79% | 75.44% |
| | | | HG005 | DeepVariant | 79.46% | 47.06% | 59.11% | 76.82% | 37.92% | 50.77% | 93.44% | 55.34% | 69.51% | 83.66% | 41.42% | 55.41% |

(b) Performance on Iso-Seq (Google)

| Read Coverage | Allele depth | Dataset | Sample | Caller | Regarding zygosity | | | | | | Disregarding zygosity | | | | | |
|---|---|---|---|---|---|---|---|---|---|---|---|---|---|---|---|---|
| | | | | | SNP | | | Indel | | | SNP | | | Indel | | |
| | | | | | Precision | Recall | F1-score | Precision | Recall | F1-score | Precision | Recall | F1-score | Precision | Recall | F1-score |
| DP≥4 | AD≥2 | Iso-Seq (Google) | HG004 | Clair3-RNA | 94.09% | 89.69% | 91.84% | 63.08% | 65.02% | 64.03% | 97.81% | 93.24% | 95.47% | 72.51% | 74.76% | 73.62% |
| | | | HG004 | LongcallR | 94.35% | 57.21% | 71.23% | \ | \ | \ | 95.47% | 57.89% | 72.08% | \ | \ | \ |
| | | | HG004 | Clair3 | 54.65% | 94.23% | 69.18% | 35.56% | 67.84% | 46.66% | 56.87% | 98.05% | 71.98% | 40.28% | 77.01% | 52.89% |
| | | | HG004 | DeepVariant | 73.91% | 12.71% | 21.69% | 41.33% | 13.94% | 20.85% | 96.46% | 16.58% | 28.30% | 56.75% | 19.20% | 28.69% |
| | | | HG005 | Clair3-RNA | 94.27% | 90.09% | 92.13% | 61.90% | 70.49% | 65.92% | 97.39% | 93.08% | 95.19% | 69.82% | 79.53% | 74.36% |
| | | | HG005 | LongcallR | 94.63% | 64.78% | 76.91% | \ | \ | \ | 95.40% | 65.31% | 77.54% | \ | \ | \ |
| | | | HG005 | Clair3 | 53.25% | 95.07% | 68.27% | 36.22% | 73.19% | 48.46% | 55.07% | 98.31% | 70.59% | 40.21% | 81.37% | 53.83% |
| | | | HG005 | DeepVariant | 71.53% | 11.79% | 20.25% | 44.52% | 13.83% | 21.11% | 96.27% | 15.87% | 27.25% | 59.39% | 18.49% | 28.21% |
| DP≥8 | AD≥2 | Iso-Seq (Google) | HG004 | Clair3-RNA | 96.80% | 91.90% | 94.29% | 67.35% | 67.67% | 67.51% | 98.37% | 93.39% | 95.81% | 75.11% | 75.48% | 75.29% |
| | | | HG004 | LongcallR | 92.94% | 72.09% | 81.20% | \ | \ | \ | 93.87% | 72.81% | 82.01% | \ | \ | \ |
| | | | HG004 | Clair3 | 53.25% | 95.70% | 68.42% | 32.05% | 71.48% | 44.26% | 54.34% | 97.68% | 69.83% | 35.93% | 80.37% | 49.66% |
| | | | HG004 | DeepVariant | 75.23% | 16.45% | 27.00% | 54.78% | 13.15% | 21.21% | 97.52% | 21.33% | 35.00% | 68.77% | 16.59% | 26.73% |
| | | | HG005 | Clair3-RNA | 96.75% | 91.89% | 94.26% | 66.46% | 73.56% | 69.83% | 97.97% | 93.04% | 95.44% | 72.53% | 80.31% | 76.22% |
| | | | HG005 | LongcallR | 92.84% | 75.83% | 83.48% | \ | \ | \ | 93.47% | 76.34% | 84.04% | \ | \ | \ |
| | | | HG005 | Clair3 | 52.08% | 96.54% | 67.66% | 33.05% | 77.28% | 46.30% | 52.89% | 98.05% | 68.71% | 36.01% | 84.34% | 50.47% |
| | | | HG005 | DeepVariant | 72.16% | 15.00% | 24.84% | 62.86% | 12.65% | 21.07% | 97.28% | 20.22% | 33.49% | 76.50% | 15.45% | 25.70% |
| DP≥10 | AD≥2 | Iso-Seq (Google) | HG004 | Clair3-RNA | 97.34% | 92.34% | 94.77% | 70.29% | 68.52% | 69.39% | 98.63% | 93.57% | 96.03% | 77.61% | 75.67% | 76.63% |
| | | | HG004 | LongcallR | 92.32% | 76.31% | 83.56% | \ | \ | \ | 93.14% | 76.98% | 84.29% | \ | \ | \ |
| | | | HG004 | Clair3 | 53.99% | 95.69% | 69.03% | 31.50% | 72.02% | 43.83% | 55.00% | 97.49% | 70.33% | 35.26% | 80.92% | 49.12% |
| | | | HG004 | DeepVariant | 75.42% | 17.78% | 28.78% | 59.60% | 13.11% | 21.50% | 97.89% | 23.08% | 37.35% | 73.15% | 16.18% | 26.50% |
| | | | HG005 | Clair3-RNA | 97.39% | 92.26% | 94.76% | 68.95% | 74.28% | 71.52% | 98.38% | 93.20% | 95.72% | 74.97% | 80.79% | 77.77% |
| | | | HG005 | LongcallR | 92.00% | 78.84% | 84.91% | \ | \ | \ | 92.56% | 79.31% | 85.42% | \ | \ | \ |
| | | | HG005 | Clair3 | 52.89% | 96.60% | 68.36% | 32.76% | 78.27% | 46.18% | 53.61% | 97.90% | 69.28% | 35.58% | 85.17% | 50.19% |
| | | | HG005 | DeepVariant | 72.18% | 16.06% | 26.27% | 67.88% | 12.21% | 20.70% | 97.88% | 21.77% | 35.62% | 81.83% | 14.79% | 25.05% |
| DP≥10 | AD≥4 | Iso-Seq (Google) | HG004 | Clair3-RNA | 97.92% | 93.53% | 95.67% | 76.70% | 73.69% | 75.16% | 99.23% | 94.79% | 96.96% | 84.00% | 80.72% | 82.33% |
| | | | HG004 | LongcallR | 92.18% | 77.13% | 83.99% | \ | \ | \ | 93.01% | 77.82% | 84.74% | \ | \ | \ |
| | | | HG004 | Clair3 | 63.31% | 96.20% | 76.36% | 35.39% | 76.63% | 48.42% | 64.43% | 97.91% | 77.72% | 39.23% | 85.27% | 53.74% |
| | | | HG004 | DeepVariant | 75.64% | 18.23% | 29.38% | 70.23% | 13.68% | 22.90% | 98.30% | 23.69% | 38.17% | 83.54% | 16.35% | 27.34% |
| | | | HG005 | Clair3-RNA | 98.11% | 93.46% | 95.73% | 75.05% | 78.11% | 76.55% | 99.11% | 94.42% | 96.71% | 81.18% | 84.52% | 82.82% |
| | | | HG005 | LongcallR | 91.87% | 79.53% | 85.25% | \ | \ | \ | 92.42% | 80.00% | 85.76% | \ | \ | \ |
| | | | HG005 | Clair3 | 62.97% | 97.18% | 76.42% | 37.16% | 81.24% | 51.00% | 63.71% | 98.33% | 77.32% | 40.15% | 87.95% | 55.13% |
| | | | HG005 | DeepVariant | 72.31% | 16.44% | 26.79% | 75.95% | 12.62% | 21.64% | 98.14% | 22.31% | 36.35% | 89.87% | 15.00% | 25.71% |

(c) Performance on MAS-Seq

| Read Coverage | Allele depth | Dataset | Sample | Caller | Regarding zygosity | | | | | | Disregarding zygosity | | | | | |
|---|---|---|---|---|---|---|---|---|---|---|---|---|---|---|---|---|
| | | | | | SNP | | | Indel | | | SNP | | | Indel | | |
| | | | | | Precision | Recall | F1-score | Precision | Recall | F1-score | Precision | Recall | F1-score | Precision | Recall | F1-score |
| DP≥4 | AD≥2 | MAS-Seq | HG004 | Clair3-RNA | 95.23% | 91.61% | 93.38% | 76.48% | 60.71% | 67.69% | 98.31% | 94.58% | 96.41% | 87.34% | 69.36% | 77.32% |
| | | | HG004 | LongcallR | 91.76% | 72.08% | 80.74% | \ | \ | \ | 93.50% | 73.45% | 82.27% | \ | \ | \ |
| | | | HG004 | Clair3 | 75.38% | 92.77% | 83.17% | 74.24% | 45.07% | 56.09% | 78.00% | 95.98% | 86.06% | 83.22% | 50.62% | 62.95% |
| | | | HG004 | DeepVariant | 72.04% | 28.26% | 40.59% | 43.15% | 23.22% | 30.19% | 87.35% | 34.27% | 49.22% | 55.71% | 30.08% | 39.07% |
| DP≥8 | AD≥2 | MAS-Seq | HG004 | Clair3-RNA | 97.63% | 93.89% | 95.72% | 78.46% | 62.17% | 69.37% | 98.93% | 95.14% | 97.00% | 87.33% | 69.22% | 77.23% |
| | | | HG004 | LongcallR | 91.34% | 92.19% | 91.77% | \ | \ | \ | 92.57% | 93.42% | 92.99% | \ | \ | \ |
| | | | HG004 | Clair3 | 81.62% | 94.10% | 87.42% | 74.25% | 44.31% | 55.50% | 83.05% | 95.74% | 88.95% | 82.89% | 49.60% | 62.06% |
| | | | HG004 | DeepVariant | 78.38% | 31.41% | 44.84% | 51.26% | 20.85% | 29.64% | 89.97% | 36.05% | 51.48% | 60.60% | 24.77% | 35.16% |
| DP≥10 | AD≥2 | MAS-Seq | HG004 | Clair3-RNA | 97.99% | 94.24% | 96.08% | 79.25% | 62.23% | 69.72% | 99.12% | 95.33% | 97.19% | 87.55% | 68.78% | 77.04% |
| | | | HG004 | LongcallR | 90.62% | 92.84% | 91.71% | \ | \ | \ | 91.64% | 93.88% | 92.75% | \ | \ | \ |
| | | | HG004 | Clair3 | 83.74% | 94.16% | 88.64% | 73.42% | 43.53% | 54.66% | 85.10% | 95.70% | 90.09% | 82.29% | 48.94% | 61.38% |
| | | | HG004 | DeepVariant | 80.23% | 33.20% | 46.96% | 54.32% | 20.54% | 29.80% | 90.93% | 37.62% | 53.22% | 62.61% | 23.80% | 34.48% |
| DP≥10 | AD≥4 | MAS-Seq | HG004 | Clair3-RNA | 98.29% | 95.30% | 96.77% | 81.55% | 66.20% | 73.08% | 99.44% | 96.41% | 97.90% | 89.32% | 72.53% | 80.05% |
| | | | HG004 | LongcallR | 90.47% | 93.83% | 92.12% | \ | \ | \ | 91.51% | 94.91% | 93.18% | \ | \ | \ |
| | | | HG004 | Clair3 | 87.41% | 95.04% | 91.06% | 76.10% | 46.05% | 57.38% | 88.85% | 96.61% | 92.57% | 84.92% | 51.55% | 64.16% |
| | | | HG004 | DeepVariant | 81.68% | 33.70% | 47.72% | 60.95% | 21.29% | 31.56% | 92.72% | 38.26% | 54.16% | 69.11% | 24.29% | 35.95% |

**Supplementary Table 4. Performance w/o tagged by REDIportal database.**

(a) Performance with variants tagged by REDIportal

| Read Coverage | Allele depth | Disregarding zygosity | Platform | Dataset | Sample | SNP performance (apply REDIportal tagging ✓) | | | | | | | SNP performance (apply REDIportal tagging ✗) | | | | | | |
|---|---|---|---|---|---|---|---|---|---|---|---|---|---|---|---|---|---|---|---|
| | | | | | | TRUTH. FP | TRUTH. FN | TRUTH. TP | QUERY. TP | Precision | Recall | F1-score | TRUTH. FP | TRUTH. FN | TRUTH. TP | QUERY. TP | Precision | Recall | F1-score |
| DP≥4 | AD≥2 | √ | ONT | cDNA | HG004 | 22,735 | 24,758 | 51,874 | 51,875 | 69.53% | 67.69% | 68.60% | 26,938 | 24,711 | 51,921 | 51922 | 65.84% | 67.75% | 66.78% |
| | | | | cDNA | HG005 | 10,337 | 25,190 | 70,338 | 70,338 | 87.19% | 73.63% | 79.84% | 15,055 | 25,129 | 70,399 | 70399 | 82.38% | 73.69% | 77.80% |
| | | | | dRNA002 | HG004 | 7,275 | 5,729 | 14,210 | 14,210 | 66.14% | 71.27% | 68.61% | 7,531 | 5,707 | 14,232 | 14232 | 65.40% | 71.38% | 68.26% |
| | | | | dRNA002 | HG005 | 7,293 | 5,167 | 14,520 | 14,520 | 66.57% | 73.75% | 69.98% | 7,620 | 5,147 | 14,540 | 14540 | 65.61% | 73.86% | 69.49% |
| | | | | dRNA004 | HG004 | 6,887 | 10,711 | 89,017 | 89,018 | 92.82% | 89.26% | 91.00% | 8,785 | 10,621 | 89,107 | 89108 | 91.03% | 89.35% | 90.18% |
| | | | | dRNA004 | HG005 | 7,349 | 10,701 | 100,093 | 100,093 | 93.16% | 90.34% | 91.73% | 9,440 | 10,607 | 100,187 | 100187 | 91.39% | 90.43% | 90.91% |
| | | | PacBio | MAS-Seq | HG004 | 13,806 | 25,222 | 275,369 | 275,387 | 95.23% | 91.61% | 93.38% | 14,718 | 25,143 | 275,448 | 275466 | 94.93% | 91.64% | 93.25% |
| | | | | Iso-Seq (BCM) | HG004 | 6,100 | 10,214 | 152,810 | 152,818 | 96.16% | 93.73% | 94.93% | 7,133 | 10,155 | 152,869 | 152877 | 95.54% | 93.77% | 94.65% |
| | | | | Iso-Seq (BCM) | HG005 | 8,175 | 12,972 | 224,196 | 224,215 | 96.48% | 94.53% | 95.50% | 10,098 | 12,916 | 224,252 | 224271 | 95.69% | 94.55% | 95.12% |
| | | | | Iso-Seq (Google) | HG004 | 6,478 | 11,853 | 103,120 | 103,124 | 94.09% | 89.69% | 91.84% | 7,388 | 11,797 | 103,176 | 103180 | 93.32% | 89.74% | 91.49% |
| | | | | Iso-Seq (Google) | HG005 | 6,672 | 12,068 | 109,705 | 109,712 | 94.27% | 90.09% | 92.13% | 7,841 | 12,015 | 109,758 | 109765 | 93.33% | 90.13% | 91.71% |
| DP≥8 | AD≥2 | √ | ONT | cDNA | HG004 | 9,087 | 9,038 | 35,485 | 35,486 | 79.61% | 79.70% | 79.66% | 10,869 | 9,003 | 35,519 | 35520 | 76.57% | 79.78% | 78.14% |
| | | | | cDNA | HG005 | 4,818 | 9,708 | 48,994 | 48,994 | 91.05% | 83.46% | 87.09% | 7,378 | 9,654 | 49,048 | 49048 | 86.92% | 83.55% | 85.21% |
| | | | | dRNA002 | HG004 | 5,954 | 3,259 | 11,828 | 11,828 | 66.52% | 78.40% | 71.97% | 6,129 | 3,245 | 11,842 | 11842 | 65.90% | 78.49% | 71.64% |
| | | | | dRNA002 | HG005 | 5,978 | 3,017 | 12,027 | 12,027 | 66.80% | 79.95% | 72.78% | 6,213 | 3,005 | 12,039 | 12039 | 65.96% | 80.03% | 72.32% |
| | | | | dRNA004 | HG004 | 3,894 | 5,356 | 65,944 | 65,945 | 94.42% | 92.49% | 93.45% | 5,030 | 5,284 | 66,016 | 66017 | 92.92% | 92.59% | 92.75% |
| | | | | dRNA004 | HG005 | 4,114 | 5,127 | 72,474 | 72,474 | 94.63% | 93.39% | 94.01% | 5,326 | 5,049 | 72,552 | 72552 | 93.16% | 93.49% | 93.33% |
| | | | PacBio | MAS-Seq | HG004 | 4,489 | 12,017 | 184,661 | 184,675 | 97.63% | 93.89% | 95.72% | 5,038 | 11,942 | 184,736 | 184750 | 97.35% | 93.93% | 95.61% |
| | | | | Iso-Seq (BCM) | HG004 | 1,601 | 4,273 | 86,697 | 86,702 | 98.19% | 95.30% | 96.72% | 2,067 | 4,233 | 86,737 | 86742 | 97.67% | 95.35% | 96.50% |
| | | | | Iso-Seq (BCM) | HG005 | 2,521 | 6,028 | 138,855 | 138,872 | 98.22% | 95.84% | 97.01% | 3,548 | 5,984 | 138,899 | 138916 | 97.51% | 95.87% | 96.68% |
| | | | | Iso-Seq (Google) | HG004 | 2,017 | 5,374 | 61,008 | 61,010 | 96.80% | 91.90% | 94.29% | 2,468 | 5,332 | 61,050 | 61052 | 96.11% | 91.97% | 94.00% |
| | | | | Iso-Seq (Google) | HG005 | 2,217 | 5,832 | 66,059 | 66,066 | 96.75% | 91.89% | 94.26% | 2,832 | 5,791 | 66,100 | 66107 | 95.89% | 91.94% | 93.88% |
| DP≥10 | AD≥2 | √ | ONT | cDNA | HG004 | 6,216 | 6,740 | 31,208 | 31,208 | 83.39% | 82.24% | 82.81% | 7,506 | 6,709 | 31,239 | 31239 | 80.63% | 82.32% | 81.47% |
| | | | | cDNA | HG005 | 3,661 | 7,456 | 42,947 | 42,947 | 92.15% | 85.21% | 88.54% | 5,694 | 7,407 | 42,996 | 42996 | 88.31% | 85.30% | 86.78% |
| | | | | dRNA002 | HG004 | 5,348 | 2,697 | 10,811 | 10,811 | 66.90% | 80.03% | 72.88% | 5,504 | 2,683 | 10,825 | 10825 | 66.29% | 80.14% | 72.56% |
| | | | | dRNA002 | HG005 | 5,327 | 2,496 | 10,972 | 10,972 | 67.32% | 81.47% | 73.72% | 5,526 | 2,485 | 10,983 | 10983 | 66.53% | 81.55% | 73.28% |
| | | | | dRNA004 | HG004 | 3,284 | 4,271 | 59,239 | 59,240 | 94.75% | 93.28% | 94.01% | 4,238 | 4,200 | 59,310 | 59311 | 93.33% | 93.39% | 93.36% |
| | | | | dRNA004 | HG005 | 3,527 | 4,000 | 64,920 | 64,920 | 94.85% | 94.20% | 94.52% | 4,505 | 3,924 | 64,996 | 64996 | 93.52% | 94.31% | 93.91% |
| | | | PacBio | MAS-Seq | HG004 | 3,229 | 9,600 | 157,111 | 157,124 | 97.99% | 94.24% | 96.08% | 3,668 | 9,528 | 157,183 | 157196 | 97.72% | 94.28% | 95.97% |
| | | | | Iso-Seq (BCM) | HG004 | 1,073 | 3,390 | 69,805 | 69,809 | 98.49% | 95.37% | 96.90% | 1,408 | 3,359 | 69,836 | 69840 | 98.02% | 95.41% | 96.70% |
| | | | | Iso-Seq (BCM) | HG005 | 1,657 | 4,711 | 114,725 | 114,740 | 98.58% | 96.06% | 97.30% | 2,416 | 4,674 | 114,762 | 114777 | 97.94% | 96.09% | 97.00% |
| | | | | Iso-Seq (Google) | HG004 | 1,399 | 4,246 | 51,187 | 51,188 | 97.34% | 92.34% | 94.77% | 1,735 | 4,208 | 51,225 | 51226 | 96.72% | 92.41% | 94.52% |
| | | | | Iso-Seq (Google) | HG005 | 1,475 | 4,612 | 55,002 | 55,009 | 97.39% | 92.26% | 94.76% | 1,933 | 4,574 | 55,040 | 55047 | 96.61% | 92.33% | 94.42% |
| DP≥10 | AD≥4 | √ | ONT | cDNA | HG004 | 5,319 | 5,558 | 30,913 | 30,913 | 85.32% | 84.76% | 85.04% | 6,557 | 5,527 | 30,944 | 30944 | 82.52% | 84.85% | 83.66% |
| | | | | cDNA | HG005 | 3,124 | 6,109 | 42,555 | 42,555 | 93.16% | 87.45% | 90.21% | 5,055 | 6,060 | 42,604 | 42604 | 89.39% | 87.55% | 88.46% |
| | | | | dRNA002 | HG004 | 5,062 | 2,034 | 10,684 | 10,684 | 67.85% | 84.01% | 75.07% | 5,195 | 2,021 | 10,697 | 10697 | 67.31% | 84.11% | 74.78% |
| | | | | dRNA002 | HG005 | 5,048 | 1,956 | 10,832 | 10,832 | 68.21% | 84.70% | 75.57% | 5,229 | 1,945 | 10,843 | 10843 | 67.47% | 84.79% | 75.14% |
| | | | | dRNA004 | HG004 | 3,099 | 3,410 | 58,680 | 58,681 | 94.98% | 94.51% | 94.75% | 4,004 | 3,339 | 58,751 | 58752 | 93.62% | 94.62% | 94.12% |
| | | | | dRNA004 | HG005 | 3,340 | 3,188 | 64,338 | 64,338 | 95.06% | 95.28% | 95.17% | 4,281 | 3,112 | 64,414 | 64414 | 93.77% | 95.39% | 94.57% |
| | | | PacBio | MAS-Seq | HG004 | 2,682 | 7,616 | 154,359 | 154,371 | 98.29% | 95.30% | 96.77% | 3,096 | 7,544 | 154,431 | 154443 | 98.03% | 95.34% | 96.67% |
| | | | | Iso-Seq (BCM) | HG004 | 891 | 2,783 | 68,241 | 68,245 | 98.71% | 96.08% | 97.38% | 1,174 | 2,752 | 68,272 | 68276 | 98.31% | 96.13% | 97.21% |
| | | | | Iso-Seq (BCM) | HG005 | 1,241 | 3,830 | 112,630 | 112,643 | 98.91% | 96.71% | 97.80% | 1,915 | 3,793 | 112,667 | 112680 | 98.33% | 96.74% | 97.53% |
| | | | | Iso-Seq (Google) | HG004 | 1,068 | 3,475 | 50,242 | 50,242 | 97.92% | 93.53% | 95.67% | 1,370 | 3,437 | 50,280 | 50280 | 97.35% | 93.60% | 95.44% |
| | | | | Iso-Seq (Google) | HG005 | 1,043 | 3,788 | 54,170 | 54,177 | 98.11% | 93.46% | 95.73% | 1,466 | 3,750 | 54,208 | 54215 | 97.37% | 93.53% | 95.41% |
| DP≥4 | AD≥2 | × | ONT | cDNA | HG004 | 14,277 | 16,300 | 60,332 | 60,333 | 80.86% | 78.73% | 79.78% | 18,471 | 16,244 | 60,388 | 60389 | 76.58% | 78.80% | 77.67% |
| | | | | cDNA | HG005 | 6,308 | 21,162 | 74,366 | 74,367 | 92.18% | 77.85% | 84.41% | 11,022 | 21,097 | 74,431 | 74432 | 87.10% | 77.92% | 82.25% |
| | | | | dRNA002 | HG004 | 6,804 | 5,258 | 14,681 | 14,681 | 68.33% | 73.63% | 70.88% | 7,059 | 5,235 | 14,704 | 14704 | 67.56% | 73.74% | 70.52% |
| | | | | dRNA002 | HG005 | 6,920 | 4,794 | 14,893 | 14,893 | 68.28% | 75.65% | 71.77% | 7,247 | 4,774 | 14,913 | 14913 | 67.30% | 75.75% | 71.27% |
| | | | | dRNA004 | HG004 | 4,188 | 8,013 | 91,715 | 91,717 | 95.63% | 91.97% | 93.76% | 6,083 | 7,920 | 91,808 | 91810 | 93.79% | 92.06% | 92.91% |
| | | | | dRNA004 | HG005 | 5,046 | 8,401 | 102,393 | 102,396 | 95.30% | 92.42% | 93.84% | 7,135 | 8,305 | 102,489 | 102492 | 93.49% | 92.50% | 93.00% |
| | | | PacBio | MAS-Seq | HG004 | 4,878 | 16,304 | 284,287 | 284,315 | 98.31% | 94.58% | 96.41% | 5,788 | 16,223 | 284,368 | 284396 | 98.01% | 94.60% | 96.27% |
| | | | | Iso-Seq (BCM) | HG004 | 1,766 | 5,883 | 157,141 | 157,152 | 98.89% | 96.39% | 97.62% | 2,796 | 5,821 | 157,203 | 157214 | 98.25% | 96.43% | 97.33% |
| | | | | Iso-Seq (BCM) | HG005 | 3,517 | 8,321 | 228,847 | 228,873 | 98.49% | 96.49% | 97.48% | 5,438 | 8,263 | 228,905 | 228931 | 97.68% | 96.52% | 97.09% |
| | | | | Iso-Seq (Google) | HG004 | 2,397 | 7,774 | 107,199 | 107,205 | 97.81% | 93.24% | 95.47% | 3,306 | 7,717 | 107,256 | 107262 | 97.01% | 93.29% | 95.11% |
| | | | | Iso-Seq (Google) | HG005 | 3,032 | 8,429 | 113,344 | 113,352 | 97.39% | 93.08% | 95.19% | 4,200 | 8,375 | 113,398 | 113406 | 96.43% | 93.12% | 94.75% |
| DP≥8 | AD≥2 | × | ONT | cDNA | HG004 | 6,921 | 6,872 | 37,651 | 37,652 | 84.47% | 84.57% | 84.52% | 8,700 | 6,834 | 37,688 | 37689 | 81.25% | 84.65% | 82.91% |
| | | | | cDNA | HG005 | 3,663 | 8,554 | 50,148 | 50,149 | 93.19% | 85.43% | 89.14% | 6,220 | 8,497 | 50,205 | 50206 | 88.98% | 85.53% | 87.22% |
| | | | | dRNA002 | HG004 | 5,719 | 3,024 | 12,063 | 12,063 | 67.84% | 79.96% | 73.40% | 5,894 | 3,010 | 12,077 | 12077 | 67.20% | 80.05% | 73.07% |
| | | | | dRNA002 | HG005 | 5,793 | 2,832 | 12,212 | 12,212 | 67.83% | 81.18% | 73.90% | 6,028 | 2,820 | 12,224 | 12224 | 66.97% | 81.26% | 73.43% |
| | | | | dRNA004 | HG004 | 2,665 | 4,128 | 67,172 | 67,174 | 96.18% | 94.21% | 95.19% | 3,799 | 4,054 | 67,246 | 67248 | 94.65% | 94.31% | 94.48% |
| | | | | dRNA004 | HG005 | 3,233 | 4,249 | 73,352 | 73,355 | 95.78% | 94.52% | 95.15% | 4,443 | 4,169 | 73,432 | 73435 | 94.29% | 94.63% | 94.46% |
| | | | PacBio | MAS-Seq | HG004 | 2,023 | 9,559 | 187,119 | 187,141 | 98.93% | 95.14% | 97.00% | 2,571 | 9,483 | 187,195 | 187217 | 98.65% | 95.18% | 96.88% |
| | | | | Iso-Seq (BCM) | HG004 | 764 | 3,438 | 87,532 | 87,539 | 99.13% | 96.22% | 97.66% | 1,228 | 3,396 | 87,574 | 87581 | 98.62% | 96.27% | 97.43% |
| | | | | Iso-Seq (BCM) | HG005 | 1,611 | 5,124 | 139,759 | 139,782 | 98.86% | 96.46% | 97.65% | 2,637 | 5,079 | 139,804 | 139827 | 98.15% | 96.49% | 97.31% |
| | | | | Iso-Seq (Google) | HG004 | 1,030 | 4,387 | 61,995 | 61,997 | 98.37% | 93.39% | 95.81% | 1,480 | 4,344 | 62,038 | 62040 | 97.67% | 93.46% | 95.52% |
| | | | | Iso-Seq (Google) | HG005 | 1,388 | 5,004 | 66,887 | 66,895 | 97.97% | 93.04% | 95.44% | 2,003 | 4,963 | 66,928 | 66936 | 97.09% | 93.10% | 95.05% |
| DP≥10 | AD≥2 | × | ONT | cDNA | HG004 | 4,899 | 5,423 | 32,525 | 32,525 | 86.91% | 85.71% | 86.31% | 6,186 | 5,389 | 32,559 | 32559 | 84.03% | 85.80% | 84.91% |
| | | | | cDNA | HG005 | 2,866 | 6,662 | 43,741 | 43,742 | 93.85% | 86.78% | 90.18% | 4,898 | 6,612 | 43,791 | 43792 | 89.94% | 86.88% | 88.38% |
| | | | | dRNA002 | HG004 | 5,165 | 2,514 | 10,994 | 10,994 | 68.04% | 81.39% | 74.12% | 5,321 | 2,500 | 11,008 | 11008 | 67.41% | 81.49% | 73.79% |
| | | | | dRNA002 | HG005 | 5,179 | 2,348 | 11,120 | 11,120 | 68.23% | 82.57% | 74.71% | 5,378 | 2,337 | 11,131 | 11131 | 67.42% | 82.65% | 74.26% |
| | | | | dRNA004 | HG004 | 2,269 | 3,257 | 60,253 | 60,255 | 96.37% | 94.87% | 95.62% | 3,222 | 3,185 | 60,325 | 60327 | 94.93% | 94.99% | 94.96% |
| | | | | dRNA004 | HG005 | 2,824 | 3,300 | 65,620 | 65,623 | 95.87% | 95.21% | 95.54% | 3,800 | 3,222 | 65,698 | 65701 | 94.53% | 95.33% | 94.93% |
| | | | PacBio | MAS-Seq | HG004 | 1,410 | 7,789 | 158,922 | 158,943 | 99.12% | 95.33% | 97.19% | 1,848 | 7,716 | 158,995 | 159016 | 98.85% | 95.37% | 97.08% |
| | | | | Iso-Seq (BCM) | HG004 | 517 | 2,836 | 70,359 | 70,365 | 99.27% | 96.13% | 97.67% | 851 | 2,804 | 70,391 | 70397 | 98.81% | 96.17% | 97.47% |
| | | | | Iso-Seq (BCM) | HG005 | 1,082 | 4,142 | 115,294 | 115,315 | 99.07% | 96.53% | 97.78% | 1,840 | 4,104 | 115,332 | 115353 | 98.43% | 96.56% | 97.49% |
| | | | | Iso-Seq (Google) | HG004 | 720 | 3,567 | 51,866 | 51,867 | 98.63% | 93.57% | 96.03% | 1,055 | 3,528 | 51,905 | 51906 | 98.01% | 93.64% | 95.77% |
| | | | | Iso-Seq (Google) | HG005 | 914 | 4,052 | 55,562 | 55,570 | 98.38% | 93.20% | 95.72% | 1,372 | 4,014 | 55,600 | 55608 | 97.59% | 93.27% | 95.38% |
| DP≥10 | AD≥4 | × | ONT | cDNA | HG004 | 4,006 | 4,245 | 32,226 | 32,226 | 88.94% | 88.36% | 88.65% | 5,241 | 4,211 | 32,260 | 32260 | 86.02% | 88.45% | 87.22% |
| | | | | cDNA | HG005 | 2,331 | 5,317 | 43,347 | 43,348 | 94.90% | 89.07% | 91.89% | 4,261 | 5,267 | 43,397 | 43398 | 91.06% | 89.18% | 90.11% |
| | | | | dRNA002 | HG004 | 4,881 | 1,853 | 10,865 | 10,865 | 69.00% | 85.43% | 76.34% | 5,014 | 1,840 | 10,878 | 10878 | 68.45% | 85.53% | 76.04% |
| | | | | dRNA002 | HG005 | 4,900 | 1,808 | 10,980 | 10,980 | 69.14% | 85.86% | 76.60% | 5,081 | 1,797 | 10,991 | 10991 | 68.39% | 85.95% | 76.17% |
| | | | | dRNA004 | HG004 | 2,087 | 2,399 | 59,691 | 59,693 | 96.62% | 96.14% | 96.38% | 2,991 | 2,327 | 59,763 | 59765 | 95.23% | 96.25% | 95.74% |
| | | | | dRNA004 | HG005 | 2,640 | 2,490 | 65,036 | 65,038 | 96.10% | 96.31% | 96.21% | 3,579 | 2,412 | 65,114 | 65116 | 94.79% | 96.43% | 95.60% |
| | | | PacBio | MAS-Seq | HG004 | 873 | 5,813 | 156,162 | 156,180 | 99.44% | 96.41% | 97.90% | 1,286 | 5,740 | 156,235 | 156253 | 99.18% | 96.46% | 97.80% |
| | | | | Iso-Seq (BCM) | HG004 | 341 | 2,235 | 68,789 | 68,795 | 99.51% | 96.85% | 98.16% | 623 | 2,203 | 68,821 | 68827 | 99.10% | 96.90% | 97.99% |
| | | | | Iso-Seq (BCM) | HG005 | 670 | 3,263 | 113,197 | 113,214 | 99.41% | 97.20% | 98.29% | 1,343 | 3,225 | 113,235 | 113252 | 98.83% | 97.23% | 98.02% |
| | | | | Iso-Seq (Google) | HG004 | 393 | 2,800 | 50,917 | 50,917 | 99.23% | 94.79% | 96.96% | 694 | 2,761 | 50,956 | 50956 | 98.66% | 94.86% | 96.72% |
| | | | | Iso-Seq (Google) | HG005 | 489 | 3,235 | 54,723 | 54,731 | 99.11% | 94.42% | 96.71% | 912 | 3,197 | 54,761 | 54769 | 98.36% | 94.48% | 96.38% |

(b) Performance without variants tagged by REDIportal

| Read Coverage | Allele depth | Disregarding zygosity | Platform | Dataset | Sample | SNP performance (apply REDIportal tagging ✓) | | | | | | | SNP performance (apply REDIportal tagging ✗) | | | | | | |
|---|---|---|---|---|---|---|---|---|---|---|---|---|---|---|---|---|---|---|
| | | | | | | TRUTH. FP | TRUTH. FN | TRUTH. TP | QUERY. TP | Precision | Recall | F1-score | TRUTH. FP | TRUTH. FN | TRUTH. TP | QUERY. TP | Precision | Recall | F1-score |
| DP≥4 | AD≥2 | √ | ONT | cDNA | HG004 | 22,735 | 24,758 | 51,874 | 51,875 | 69.53% | 67.69% | 68.60% | 26,938 | 24,711 | 51,921 | 51922 | 65.84% | 67.75% | 66.78% |
| | | | | cDNA | HG005 | 10,337 | 25,190 | 70,338 | 70,338 | 87.19% | 73.63% | 79.84% | 15,055 | 25,129 | 70,399 | 70399 | 82.38% | 73.69% | 77.80% |
| | | | | dRNA002 | HG004 | 7,275 | 5,729 | 14,210 | 14,210 | 66.14% | 71.27% | 68.61% | 7,531 | 5,707 | 14,232 | 14232 | 65.40% | 71.38% | 68.26% |
| | | | | dRNA002 | HG005 | 7,293 | 5,167 | 14,520 | 14,520 | 66.57% | 73.75% | 69.98% | 7,620 | 5,147 | 14,540 | 14540 | 65.61% | 73.86% | 69.49% |
| | | | | dRNA004 | HG004 | 6,887 | 10,711 | 89,017 | 89,018 | 92.82% | 89.26% | 91.00% | 8,785 | 10,621 | 89,107 | 89108 | 91.03% | 89.35% | 90.18% |
| | | | | dRNA004 | HG005 | 7,349 | 10,701 | 100,093 | 100,093 | 93.16% | 90.34% | 91.73% | 9,440 | 10,607 | 100,187 | 100187 | 91.39% | 90.43% | 90.91% |
| | | | PacBio | MAS-Seq | HG004 | 13,806 | 25,222 | 275,369 | 275,387 | 95.23% | 91.61% | 93.38% | 14,718 | 25,143 | 275,448 | 275466 | 94.93% | 91.64% | 93.25% |
| | | | | Iso-Seq (BCM) | HG004 | 6,100 | 10,214 | 152,810 | 152,818 | 96.16% | 93.73% | 94.93% | 7,133 | 10,155 | 152,869 | 152877 | 95.54% | 93.77% | 94.65% |
| | | | | Iso-Seq (BCM) | HG005 | 8,175 | 12,972 | 224,196 | 224,215 | 96.48% | 94.53% | 95.50% | 10,098 | 12,916 | 224,252 | 224271 | 95.69% | 94.55% | 95.12% |
| | | | | Iso-Seq (Google) | HG004 | 6,478 | 11,853 | 103,120 | 103,124 | 94.09% | 89.69% | 91.84% | 7,388 | 11,797 | 103,176 | 103180 | 93.32% | 89.74% | 91.49% |
| | | | | Iso-Seq (Google) | HG005 | 6,672 | 12,068 | 109,705 | 109,712 | 94.27% | 90.09% | 92.13% | 7,841 | 12,015 | 109,758 | 109765 | 93.33% | 90.13% | 91.71% |
| DP≥8 | AD≥2 | √ | ONT | cDNA | HG004 | 9,087 | 9,038 | 35,485 | 35,486 | 79.61% | 79.70% | 79.66% | 10,869 | 9,003 | 35,519 | 35520 | 76.57% | 79.78% | 78.14% |
| | | | | cDNA | HG005 | 4,818 | 9,708 | 48,994 | 48,994 | 91.05% | 83.46% | 87.09% | 7,378 | 9,654 | 49,048 | 49048 | 86.92% | 83.55% | 85.21% |
| | | | | dRNA002 | HG004 | 5,954 | 3,259 | 11,828 | 11,828 | 66.52% | 78.40% | 71.97% | 6,129 | 3,245 | 11,842 | 11842 | 65.90% | 78.49% | 71.64% |
| | | | | dRNA002 | HG005 | 5,978 | 3,017 | 12,027 | 12,027 | 66.80% | 79.95% | 72.78% | 6,213 | 3,005 | 12,039 | 12039 | 65.96% | 80.03% | 72.32% |
| | | | | dRNA004 | HG004 | 3,894 | 5,356 | 65,944 | 65,945 | 94.42% | 92.49% | 93.45% | 5,030 | 5,284 | 66,016 | 66017 | 92.92% | 92.59% | 92.75% |
| | | | | dRNA004 | HG005 | 4,114 | 5,127 | 72,474 | 72,474 | 94.63% | 93.39% | 94.01% | 5,326 | 5,049 | 72,552 | 72552 | 93.16% | 93.49% | 93.33% |
| | | | PacBio | MAS-Seq | HG004 | 4,489 | 12,017 | 184,661 | 184,675 | 97.63% | 93.89% | 95.72% | 5,038 | 11,942 | 184,736 | 184750 | 97.35% | 93.93% | 95.61% |
| | | | | Iso-Seq (BCM) | HG004 | 1,601 | 4,273 | 86,697 | 86,702 | 98.19% | 95.30% | 96.72% | 2,067 | 4,233 | 86,737 | 86742 | 97.67% | 95.35% | 96.50% |
| | | | | Iso-Seq (BCM) | HG005 | 2,521 | 6,028 | 138,855 | 138,872 | 98.22% | 95.84% | 97.01% | 3,548 | 5,984 | 138,899 | 138916 | 97.51% | 95.87% | 96.68% |
| | | | | Iso-Seq (Google) | HG004 | 2,017 | 5,374 | 61,008 | 61,010 | 96.80% | 91.90% | 94.29% | 2,468 | 5,332 | 61,050 | 61052 | 96.11% | 91.97% | 94.00% |
| | | | | Iso-Seq (Google) | HG005 | 2,217 | 5,832 | 66,059 | 66,066 | 96.75% | 91.89% | 94.26% | 2,832 | 5,791 | 66,100 | 66107 | 95.89% | 91.94% | 93.88% |
| DP≥10 | AD≥2 | √ | ONT | cDNA | HG004 | 6,216 | 6,740 | 31,208 | 31,208 | 83.39% | 82.24% | 82.81% | 7,506 | 6,709 | 31,239 | 31239 | 80.63% | 82.32% | 81.47% |
| | | | | cDNA | HG005 | 3,661 | 7,456 | 42,947 | 42,947 | 92.15% | 85.21% | 88.54% | 5,694 | 7,407 | 42,996 | 42996 | 88.31% | 85.30% | 86.78% |
| | | | | dRNA002 | HG004 | 5,348 | 2,697 | 10,811 | 10,811 | 66.90% | 80.03% | 72.88% | 5,504 | 2,683 | 10,825 | 10825 | 66.29% | 80.14% | 72.56% |
| | | | | dRNA002 | HG005 | 5,327 | 2,496 | 10,972 | 10,972 | 67.32% | 81.47% | 73.72% | 5,526 | 2,485 | 10,983 | 10983 | 66.53% | 81.55% | 73.28% |
| | | | | dRNA004 | HG004 | 3,284 | 4,271 | 59,239 | 59,240 | 94.75% | 93.28% | 94.01% | 4,238 | 4,200 | 59,310 | 59311 | 93.33% | 93.39% | 93.36% |
| | | | | dRNA004 | HG005 | 3,527 | 4,000 | 64,920 | 64,920 | 94.85% | 94.20% | 94.52% | 4,505 | 3,924 | 64,996 | 64996 | 93.52% | 94.31% | 93.91% |
| | | | PacBio | MAS-Seq | HG004 | 3,229 | 9,600 | 157,111 | 157,124 | 97.99% | 94.24% | 96.08% | 3,668 | 9,528 | 157,183 | 157196 | 97.72% | 94.28% | 95.97% |
| | | | | Iso-Seq (BCM) | HG004 | 1,073 | 3,390 | 69,805 | 69,809 | 98.49% | 95.37% | 96.90% | 1,408 | 3,359 | 69,836 | 69840 | 98.02% | 95.41% | 96.70% |
| | | | | Iso-Seq (BCM) | HG005 | 1,657 | 4,711 | 114,725 | 114,740 | 98.58% | 96.06% | 97.30% | 2,416 | 4,674 | 114,762 | 114777 | 97.94% | 96.09% | 97.00% |
| | | | | Iso-Seq (Google) | HG004 | 1,399 | 4,246 | 51,187 | 51,188 | 97.34% | 92.34% | 94.77% | 1,735 | 4,208 | 51,225 | 51226 | 96.72% | 92.41% | 94.52% |
| | | | | Iso-Seq (Google) | HG005 | 1,475 | 4,612 | 55,002 | 55,009 | 97.39% | 92.26% | 94.76% | 1,933 | 4,574 | 55,040 | 55047 | 96.61% | 92.33% | 94.42% |
| DP≥10 | AD≥4 | √ | ONT | cDNA | HG004 | 5,319 | 5,558 | 30,913 | 30,913 | 85.32% | 84.76% | 85.04% | 6,557 | 5,527 | 30,944 | 30944 | 82.52% | 84.85% | 83.66% |
| | | | | cDNA | HG005 | 3,124 | 6,109 | 42,555 | 42,555 | 93.16% | 87.45% | 90.21% | 5,055 | 6,060 | 42,604 | 42604 | 89.39% | 87.55% | 88.46% |
| | | | | dRNA002 | HG004 | 5,062 | 2,034 | 10,684 | 10,684 | 67.85% | 84.01% | 75.07% | 5,195 | 2,021 | 10,697 | 10697 | 67.31% | 84.11% | 74.78% |
| | | | | dRNA002 | HG005 | 5,048 | 1,956 | 10,832 | 10,832 | 68.21% | 84.70% | 75.57% | 5,229 | 1,945 | 10,843 | 10843 | 67.47% | 84.79% | 75.14% |
| | | | | dRNA004 | HG004 | 3,099 | 3,410 | 58,680 | 58,681 | 94.98% | 94.51% | 94.75% | 4,004 | 3,339 | 58,751 | 58752 | 93.62% | 94.62% | 94.12% |
| | | | | dRNA004 | HG005 | 3,340 | 3,188 | 64,338 | 64,338 | 95.06% | 95.28% | 95.17% | 4,281 | 3,112 | 64,414 | 64414 | 93.77% | 95.39% | 94.57% |
| | | | PacBio | MAS-Seq | HG004 | 2,682 | 7,616 | 154,359 | 154,371 | 98.29% | 95.30% | 96.77% | 3,096 | 7,544 | 154,431 | 154443 | 98.03% | 95.34% | 96.67% |
| | | | | Iso-Seq (BCM) | HG004 | 891 | 2,783 | 68,241 | 68,245 | 98.71% | 96.08% | 97.38% | 1,174 | 2,752 | 68,272 | 68276 | 98.31% | 96.13% | 97.21% |
| | | | | Iso-Seq (BCM) | HG005 | 1,241 | 3,830 | 112,630 | 112,643 | 98.91% | 96.71% | 97.80% | 1,915 | 3,793 | 112,667 | 112680 | 98.33% | 96.74% | 97.53% |
| | | | | Iso-Seq (Google) | HG004 | 1,068 | 3,475 | 50,242 | 50,242 | 97.92% | 93.53% | 95.67% | 1,370 | 3,437 | 50,280 | 50280 | 97.35% | 93.60% | 95.44% |
| | | | | Iso-Seq (Google) | HG005 | 1,043 | 3,788 | 54,170 | 54,177 | 98.11% | 93.46% | 95.73% | 1,466 | 3,750 | 54,208 | 54215 | 97.37% | 93.53% | 95.41% |
| DP≥4 | AD≥2 | × | ONT | cDNA | HG004 | 14,277 | 16,300 | 60,332 | 60,333 | 80.86% | 78.73% | 79.78% | 18,471 | 16,244 | 60,388 | 60389 | 76.58% | 78.80% | 77.67% |
| | | | | cDNA | HG005 | 6,308 | 21,162 | 74,366 | 74,367 | 92.18% | 77.85% | 84.41% | 11,022 | 21,097 | 74,431 | 74432 | 87.10% | 77.92% | 82.25% |
| | | | | dRNA002 | HG004 | 6,804 | 5,258 | 14,681 | 14,681 | 68.33% | 73.63% | 70.88% | 7,059 | 5,235 | 14,704 | 14704 | 67.56% | 73.74% | 70.52% |
| | | | | dRNA002 | HG005 | 6,920 | 4,794 | 14,893 | 14,893 | 68.28% | 75.65% | 71.77% | 7,247 | 4,774 | 14,913 | 14913 | 67.30% | 75.75% | 71.27% |
| | | | | dRNA004 | HG004 | 4,188 | 8,013 | 91,715 | 91,717 | 95.63% | 91.97% | 93.76% | 6,083 | 7,920 | 91,808 | 91810 | 93.79% | 92.06% | 92.91% |
| | | | | dRNA004 | HG005 | 5,046 | 8,401 | 102,393 | 102,396 | 95.30% | 92.42% | 93.84% | 7,135 | 8,305 | 102,489 | 102492 | 93.49% | 92.50% | 93.00% |
| | | | PacBio | MAS-Seq | HG004 | 4,878 | 16,304 | 284,287 | 284,315 | 98.31% | 94.58% | 96.41% | 5,788 | 16,223 | 284,368 | 284396 | 98.01% | 94.60% | 96.27% |
| | | | | Iso-Seq (BCM) | HG004 | 1,766 | 5,883 | 157,141 | 157,152 | 98.89% | 96.39% | 97.62% | 2,796 | 5,821 | 157,203 | 157214 | 98.25% | 96.43% | 97.33% |
| | | | | Iso-Seq (BCM) | HG005 | 3,517 | 8,321 | 228,847 | 228,873 | 98.49% | 96.49% | 97.48% | 5,438 | 8,263 | 228,905 | 228931 | 97.68% | 96.52% | 97.09% |
| | | | | Iso-Seq (Google) | HG004 | 2,397 | 7,774 | 107,199 | 107,205 | 97.81% | 93.24% | 95.47% | 3,306 | 7,717 | 107,256 | 107262 | 97.01% | 93.29% | 95.11% |
| | | | | Iso-Seq (Google) | HG005 | 3,032 | 8,429 | 113,344 | 113,352 | 97.39% | 93.08% | 95.19% | 4,200 | 8,375 | 113,398 | 113406 | 96.43% | 93.12% | 94.75% |
| DP≥8 | AD≥2 | × | ONT | cDNA | HG004 | 6,921 | 6,872 | 37,651 | 37,652 | 84.47% | 84.57% | 84.52% | 8,700 | 6,834 | 37,688 | 37689 | 81.25% | 84.65% | 82.91% |
| | | | | cDNA | HG005 | 3,663 | 8,554 | 50,148 | 50,149 | 93.19% | 85.43% | 89.14% | 6,220 | 8,497 | 50,205 | 50206 | 88.98% | 85.53% | 87.22% |
| | | | | dRNA002 | HG004 | 5,719 | 3,024 | 12,063 | 12,063 | 67.84% | 79.96% | 73.40% | 5,894 | 3,010 | 12,077 | 12077 | 67.20% | 80.05% | 73.07% |
| | | | | dRNA002 | HG005 | 5,793 | 2,832 | 12,212 | 12,212 | 67.83% | 81.18% | 73.90% | 6,028 | 2,820 | 12,224 | 12224 | 66.97% | 81.26% | 73.43% |
| | | | | dRNA004 | HG004 | 2,665 | 4,128 | 67,172 | 67,174 | 96.18% | 94.21% | 95.19% | 3,799 | 4,054 | 67,246 | 67248 | 94.65% | 94.31% | 94.48% |
| | | | | dRNA004 | HG005 | 3,233 | 4,249 | 73,352 | 73,355 | 95.78% | 94.52% | 95.15% | 4,443 | 4,169 | 73,432 | 73435 | 94.29% | 94.63% | 94.46% |
| | | | PacBio | MAS-Seq | HG004 | 2,023 | 9,559 | 187,119 | 187,141 | 98.93% | 95.14% | 97.00% | 2,571 | 9,483 | 187,195 | 187217 | 98.65% | 95.18% | 96.88% |
| | | | | Iso-Seq (BCM) | HG004 | 764 | 3,438 | 87,532 | 87,539 | 99.13% | 96.22% | 97.66% | 1,228 | 3,396 | 87,574 | 87581 | 98.62% | 96.27% | 97.43% |
| | | | | Iso-Seq (BCM) | HG005 | 1,611 | 5,124 | 139,759 | 139,782 | 98.86% | 96.46% | 97.65% | 2,637 | 5,079 | 139,804 | 139827 | 98.15% | 96.49% | 97.31% |
| | | | | Iso-Seq (Google) | HG004 | 1,030 | 4,387 | 61,995 | 61,997 | 98.37% | 93.39% | 95.81% | 1,480 | 4,344 | 62,038 | 62040 | 97.67% | 93.46% | 95.52% |
| | | | | Iso-Seq (Google) | HG005 | 1,388 | 5,004 | 66,887 | 66,895 | 97.97% | 93.04% | 95.44% | 2,003 | 4,963 | 66,928 | 66936 | 97.09% | 93.10% | 95.05% |
| DP≥10 | AD≥2 | × | ONT | cDNA | HG004 | 4,899 | 5,423 | 32,525 | 32,525 | 86.91% | 85.71% | 86.31% | 6,186 | 5,389 | 32,559 | 32559 | 84.03% | 85.80% | 84.91% |
| | | | | cDNA | HG005 | 2,866 | 6,662 | 43,741 | 43,742 | 93.85% | 86.78% | 90.18% | 4,898 | 6,612 | 43,791 | 43792 | 89.94% | 86.88% | 88.38% |
| | | | | dRNA002 | HG004 | 5,165 | 2,514 | 10,994 | 10,994 | 68.04% | 81.39% | 74.12% | 5,321 | 2,500 | 11,008 | 11008 | 67.41% | 81.49% | 73.79% |
| | | | | dRNA002 | HG005 | 5,179 | 2,348 | 11,120 | 11,120 | 68.23% | 82.57% | 74.71% | 5,378 | 2,337 | 11,131 | 11131 | 67.42% | 82.65% | 74.26% |
| | | | | dRNA004 | HG004 | 2,269 | 3,257 | 60,253 | 60,255 | 96.37% | 94.87% | 95.62% | 3,222 | 3,185 | 60,325 | 60327 | 94.93% | 94.99% | 94.96% |
| | | | | dRNA004 | HG005 | 2,824 | 3,300 | 65,620 | 65,623 | 95.87% | 95.21% | 95.54% | 3,800 | 3,222 | 65,698 | 65701 | 94.53% | 95.33% | 94.93% |
| | | | PacBio | MAS-Seq | HG004 | 1,410 | 7,789 | 158,922 | 158,943 | 99.12% | 95.33% | 97.19% | 1,848 | 7,716 | 158,995 | 159016 | 98.85% | 95.37% | 97.08% |
| | | | | Iso-Seq (BCM) | HG004 | 517 | 2,836 | 70,359 | 70,365 | 99.27% | 96.13% | 97.67% | 851 | 2,804 | 70,391 | 70397 | 98.81% | 96.17% | 97.47% |
| | | | | Iso-Seq (BCM) | HG005 | 1,082 | 4,142 | 115,294 | 115,315 | 99.07% | 96.53% | 97.78% | 1,840 | 4,104 | 115,332 | 115353 | 98.43% | 96.56% | 97.49% |
| | | | | Iso-Seq (Google) | HG004 | 720 | 3,567 | 51,866 | 51,867 | 98.63% | 93.57% | 96.03% | 1,055 | 3,528 | 51,905 | 51906 | 98.01% | 93.64% | 95.77% |
| | | | | Iso-Seq (Google) | HG005 | 914 | 4,052 | 55,562 | 55,570 | 98.38% | 93.20% | 95.72% | 1,372 | 4,014 | 55,600 | 55608 | 97.59% | 93.27% | 95.38% |
| DP≥10 | AD≥4 | × | ONT | cDNA | HG004 | 4,006 | 4,245 | 32,226 | 32,226 | 88.94% | 88.36% | 88.65% | 5,241 | 4,211 | 32,260 | 32260 | 86.02% | 88.45% | 87.22% |
| | | | | cDNA | HG005 | 2,331 | 5,317 | 43,347 | 43,348 | 94.90% | 89.07% | 91.89% | 4,261 | 5,267 | 43,397 | 43398 | 91.06% | 89.18% | 90.11% |
| | | | | dRNA002 | HG004 | 4,881 | 1,853 | 10,865 | 10,865 | 69.00% | 85.43% | 76.34% | 5,014 | 1,840 | 10,878 | 10878 | 68.45% | 85.53% | 76.04% |
| | | | | dRNA002 | HG005 | 4,900 | 1,808 | 10,980 | 10,980 | 69.14% | 85.86% | 76.60% | 5,081 | 1,797 | 10,991 | 10991 | 68.39% | 85.95% | 76.17% |
| | | | | dRNA004 | HG004 | 2,087 | 2,399 | 59,691 | 59,693 | 96.62% | 96.14% | 96.38% | 2,991 | 2,327 | 59,763 | 59765 | 95.23% | 96.25% | 95.74% |
| | | | | dRNA004 | HG005 | 2,640 | 2,490 | 65,036 | 65,038 | 96.10% | 96.31% | 96.21% | 3,579 | 2,412 | 65,114 | 65116 | 94.79% | 96.43% | 95.60% |
| | | | PacBio | MAS-Seq | HG004 | 873 | 5,813 | 156,162 | 156,180 | 99.44% | 96.41% | 97.90% | 1,286 | 5,740 | 156,235 | 156253 | 99.18% | 96.46% | 97.80% |
| | | | | Iso-Seq (BCM) | HG004 | 341 | 2,235 | 68,789 | 68,795 | 99.51% | 96.85% | 98.16% | 623 | 2,203 | 68,821 | 68827 | 99.10% | 96.90% | 97.99% |
| | | | | Iso-Seq (BCM) | HG005 | 670 | 3,263 | 113,197 | 113,214 | 99.41% | 97.20% | 98.29% | 1,343 | 3,225 | 113,235 | 113252 | 98.83% | 97.23% | 98.02% |
| | | | | Iso-Seq (Google) | HG004 | 393 | 2,800 | 50,917 | 50,917 | 99.23% | 94.79% | 96.96% | 694 | 2,761 | 50,956 | 50956 | 98.66% | 94.86% | 96.72% |
| | | | | Iso-Seq (Google) | HG005 | 489 | 3,235 | 54,723 | 54,731 | 99.11% | 94.42% | 96.71% | 912 | 3,197 | 54,761 | 54769 | 98.36% | 94.48% | 96.38% |

**Supplementary Table 5. PacBio performance using miniamp2 and pbmm2 aligners.**

| Read Coverage | Allele depth | Dataset | Aligner | Sample | Regarding zygosity | | | | | | Disregarding zygosity | | | | | |
|---|---|---|---|---|---|---|---|---|---|---|---|---|---|---|---|---|
| | | | | | SNP | | | Indel | | | SNP | | | Indel | | |
| | | | | | Precision | Recall | F1-score | Precision | Recall | F1-score | Precision | Recall | F1-score | Precision | Recall | F1-score |
| DP≥4 | AD≥2 | Iso-Seq (BCM) | pbmm2 | HG004 | 95.07% | 93.34% | 94.20% | 80.30% | 69.26% | 74.37% | 98.28% | 96.49% | 97.38% | 90.68% | 78.24% | 84.00% |
| | | | | HG005 | 95.23% | 94.17% | 94.70% | 84.76% | 76.88% | 80.62% | 97.70% | 96.61% | 97.15% | 93.02% | 84.38% | 88.49% |
| | | | minimap2 | HG004 | 96.16% | 93.73% | 94.93% | 80.74% | 70.21% | 75.11% | 98.89% | 96.39% | 97.62% | 90.96% | 79.13% | 84.64% |
| | | | | HG005 | 96.48% | 94.53% | 95.50% | 85.47% | 77.69% | 81.40% | 98.49% | 96.49% | 97.48% | 93.45% | 84.98% | 89.02% |
| | | Iso-Seq (Google) | pbmm2 | HG004 | 92.20% | 90.19% | 91.18% | 64.35% | 64.96% | 64.66% | 96.30% | 94.19% | 95.23% | 73.85% | 74.58% | 74.21% |
| | | | | HG005 | 92.29% | 90.83% | 91.55% | 62.75% | 69.55% | 65.98% | 95.82% | 94.31% | 95.06% | 71.40% | 79.15% | 75.08% |
| | | | minimap2 | HG004 | 94.09% | 89.69% | 91.84% | 63.08% | 65.02% | 64.03% | 97.81% | 93.24% | 95.47% | 72.51% | 74.76% | 73.62% |
| | | | | HG005 | 94.27% | 90.09% | 92.13% | 61.90% | 70.49% | 65.92% | 97.39% | 93.08% | 95.19% | 69.82% | 79.53% | 74.36% |
| | | MAS-Seq | pbmm2 | HG004 | 94.04% | 91.51% | 92.76% | 74.48% | 61.99% | 67.67% | 97.84% | 95.20% | 96.50% | 85.45% | 71.15% | 77.65% |
| | | | minimap2 | HG004 | 95.23% | 91.61% | 93.38% | 76.48% | 60.71% | 67.69% | 98.31% | 94.58% | 96.41% | 87.34% | 69.36% | 77.32% |
| DP≥8 | AD≥2 | Iso-Seq (BCM) | pbmm2 | HG004 | 97.27% | 94.98% | 96.11% | 83.57% | 72.82% | 77.83% | 98.68% | 96.37% | 97.51% | 91.16% | 79.46% | 84.91% |
| | | | | HG005 | 97.17% | 95.71% | 96.43% | 87.98% | 80.94% | 84.31% | 98.20% | 96.72% | 97.45% | 93.68% | 86.21% | 89.79% |
| | | | minimap2 | HG004 | 98.19% | 95.30% | 96.72% | 83.56% | 73.68% | 78.31% | 99.13% | 96.22% | 97.66% | 91.35% | 80.58% | 85.62% |
| | | | | HG005 | 98.22% | 95.84% | 97.01% | 88.49% | 81.53% | 84.87% | 98.86% | 96.46% | 97.65% | 94.08% | 86.71% | 90.24% |
| | | Iso-Seq (Google) | pbmm2 | HG004 | 95.33% | 92.61% | 93.95% | 69.78% | 67.82% | 68.79% | 97.17% | 94.40% | 95.77% | 77.58% | 75.44% | 76.50% |
| | | | | HG005 | 95.14% | 92.96% | 94.03% | 68.14% | 73.15% | 70.56% | 96.63% | 94.41% | 95.51% | 75.06% | 80.59% | 77.72% |
| | | | minimap2 | HG004 | 96.80% | 91.90% | 94.29% | 67.35% | 67.67% | 67.51% | 98.37% | 93.39% | 95.81% | 75.11% | 75.48% | 75.29% |
| | | | | HG005 | 96.75% | 91.89% | 94.26% | 66.46% | 73.56% | 69.83% | 97.97% | 93.04% | 95.44% | 72.53% | 80.31% | 76.22% |
| | | MAS-Seq | pbmm2 | HG004 | 96.76% | 93.56% | 95.13% | 77.10% | 63.18% | 69.45% | 98.62% | 95.37% | 96.97% | 86.17% | 70.65% | 77.64% |
| | | | minimap2 | HG004 | 97.63% | 93.89% | 95.72% | 78.46% | 62.17% | 69.37% | 98.93% | 95.14% | 97.00% | 87.33% | 69.22% | 77.23% |
| DP≥10 | AD≥2 | Iso-Seq (BCM) | pbmm2 | HG004 | 97.83% | 95.32% | 96.56% | 84.65% | 73.90% | 78.91% | 98.82% | 96.28% | 97.53% | 91.55% | 79.96% | 85.36% |
| | | | | HG005 | 97.83% | 96.17% | 96.99% | 88.87% | 82.19% | 85.40% | 98.54% | 96.86% | 97.70% | 93.99% | 86.94% | 90.33% |
| | | | minimap2 | HG004 | 98.49% | 95.37% | 96.90% | 84.42% | 74.48% | 79.14% | 99.27% | 96.13% | 97.67% | 91.63% | 80.89% | 85.93% |
| | | | | HG005 | 98.58% | 96.06% | 97.30% | 89.30% | 82.49% | 85.76% | 99.07% | 96.53% | 97.78% | 94.38% | 87.22% | 90.66% |
| | | Iso-Seq (Google) | pbmm2 | HG004 | 96.18% | 93.03% | 94.58% | 72.50% | 68.58% | 70.49% | 97.68% | 94.48% | 96.05% | 79.95% | 75.66% | 77.75% |
| | | | | HG005 | 96.28% | 93.43% | 94.83% | 70.94% | 74.07% | 72.47% | 97.43% | 94.55% | 95.97% | 77.56% | 80.98% | 79.23% |
| | | | minimap2 | HG004 | 97.34% | 92.34% | 94.77% | 70.29% | 68.52% | 69.39% | 98.63% | 93.57% | 96.03% | 77.61% | 75.67% | 76.63% |
| | | | | HG005 | 97.39% | 92.26% | 94.76% | 68.95% | 74.28% | 71.52% | 98.38% | 93.20% | 95.72% | 74.97% | 80.79% | 77.77% |
| | | MAS-Seq | pbmm2 | HG004 | 97.48% | 94.12% | 95.77% | 78.22% | 63.09% | 69.85% | 98.87% | 95.46% | 97.14% | 86.72% | 69.99% | 77.46% |
| | | | minimap2 | HG004 | 97.99% | 94.24% | 96.08% | 79.25% | 62.23% | 69.72% | 99.12% | 95.33% | 97.19% | 87.55% | 68.78% | 77.04% |
| DP≥10 | AD≥4 | Iso-Seq (BCM) | pbmm2 | HG004 | 98.13% | 95.97% | 97.04% | 87.07% | 79.31% | 83.01% | 99.14% | 96.96% | 98.04% | 93.24% | 84.96% | 88.90% |
| | | | | HG005 | 98.31% | 96.79% | 97.54% | 90.16% | 85.84% | 87.95% | 99.03% | 97.50% | 98.26% | 94.87% | 90.33% | 92.55% |
| | | | minimap2 | HG004 | 98.71% | 96.08% | 97.38% | 86.55% | 79.41% | 82.83% | 99.51% | 96.85% | 98.16% | 93.40% | 85.75% | 89.42% |
| | | | | HG005 | 98.91% | 96.71% | 97.80% | 90.67% | 85.95% | 88.25% | 99.41% | 97.20% | 98.29% | 95.43% | 90.49% | 92.90% |
| | | Iso-Seq (Google) | pbmm2 | HG004 | 96.99% | 94.17% | 95.56% | 77.84% | 74.31% | 76.03% | 98.52% | 95.65% | 97.07% | 84.86% | 81.07% | 82.92% |
| | | | | HG005 | 97.31% | 94.56% | 95.92% | 76.09% | 78.38% | 77.22% | 98.49% | 95.71% | 97.08% | 82.53% | 85.03% | 83.76% |
| | | | minimap2 | HG004 | 97.92% | 93.53% | 95.67% | 76.70% | 73.69% | 75.16% | 99.23% | 94.79% | 96.96% | 84.00% | 80.72% | 82.33% |
| | | | | HG005 | 98.11% | 93.46% | 95.73% | 75.05% | 78.11% | 76.55% | 99.11% | 94.42% | 96.71% | 81.18% | 84.52% | 82.82% |
| | | MAS-Seq | pbmm2 | HG004 | 97.86% | 95.13% | 96.48% | 80.77% | 67.07% | 73.28% | 99.28% | 96.50% | 97.87% | 88.50% | 73.52% | 80.32% |
| | | | minimap2 | HG004 | 98.29% | 95.30% | 96.77% | 81.55% | 66.20% | 73.08% | 99.44% | 96.41% | 97.90% | 89.32% | 72.53% | 80.05% |

**Supplementary Table 6. Performance by genomic context on PacBio and ONT datasets.**

(a) Performance on PacBio

| Dataset | Stratification type | Stratification subtype | Clair3-RNA SNP performance | | | | | | LongcallR SNP performance | | | | | |
|---|---|---|---|---|---|---|---|---|---|---|---|---|---|---|
| | | | Precision | Recall | F1-score | FP | FN | TP | Precision | Recall | F1-score | FP | FN | TP |
| PacBio Iso-Seq (BCM) HG004 | Low complexity | Homopol 4-6bp | 91.49% | 95.68% | 87.65% | 1,747 | 5,455 | 38,726 | 73.18% | 94.63% | 59.66% | 1,495 | 17,824 | 26,356 |
| | | Homopol 7-11bp | 85.22% | 91.15% | 80.01% | 313 | 805 | 3,221 | 61.03% | 91.64% | 45.75% | 168 | 2,184 | 1,842 |
| | | Homopol gt11bp | 74.76% | 83.67% | 67.57% | 122 | 300 | 625 | 37.71% | 68.97% | 25.95% | 108 | 685 | 240 |
| | | Imp Homopol gt10bp | 80.44% | 87.96% | 74.10% | 245 | 625 | 1,788 | 51.63% | 83.01% | 37.46% | 185 | 1,509 | 904 |
| | | TR lt51bp | 88.29% | 93.49% | 83.64% | 109 | 304 | 1,554 | 71.03% | 92.64% | 57.59% | 85 | 788 | 1,070 |
| | | TR51-220bp | 89.42% | 95.05% | 84.41% | 47 | 166 | 899 | 72.62% | 92.20% | 59.91% | 54 | 427 | 638 |
| | | TR201-10kbp | 87.32% | 93.33% | 82.04% | 24 | 74 | 338 | 70.17% | 96.60% | 55.10% | 8 | 185 | 227 |
| | | TR gt100bp | 87.27% | 93.70% | 81.66% | 43 | 144 | 641 | 70.47% | 95.63% | 55.80% | 20 | 347 | 438 |
| | | TR and Homopol | 85.94% | 91.90% | 80.70% | 621 | 1,682 | 7,035 | 63.76% | 90.71% | 49.16% | 439 | 4,432 | 4,285 |
| | Segmental duplications | ChainSelf | 88.54% | 93.06% | 84.43% | 784 | 1,938 | 10,512 | 73.82% | 91.58% | 61.83% | 708 | 4,752 | 7,698 |
| | | ChainSelf gt10kb | 84.06% | 88.83% | 79.78% | 391 | 787 | 3,106 | 63.49% | 90.74% | 48.83% | 194 | 1,992 | 1,901 |
| | | SegDups | 86.90% | 91.41% | 82.81% | 617 | 1,362 | 6,561 | 70.89% | 93.40% | 57.12% | 320 | 3,397 | 4,526 |
| | | SegDups gt10kb | 85.25% | 90.17% | 80.84% | 536 | 1,165 | 4,915 | 66.45% | 92.36% | 51.89% | 261 | 2,925 | 3,155 |
| | Low mappability | LowMAD | 85.30% | 88.20% | 82.58% | 733 | 1,155 | 5,477 | 64.97% | 91.51% | 50.36% | 310 | 3,292 | 3,340 |
| | Other difficult regions | L1H | 89.68% | 90.00% | 89.36% | 14 | 15 | 126 | 68.84% | 99.92% | 52.48% | 0 | 67 | 74 |
| | | MHC | 76.90% | 94.37% | 64.89% | 110 | 997 | 1,843 | 47.37% | 96.33% | 31.41% | 34 | 1,948 | 892 |
| | Functional regions | CDS | 95.23% | 98.37% | 92.27% | 141 | 713 | 8,516 | 91.75% | 98.87% | 85.58% | 90 | 1,331 | 7,898 |
| PacBio MAS-Seq HG004 | Low complexity | Homopol_4-6bp | 89.81% | 94.90% | 85.24% | 3,665 | 11,816 | 68,214 | 78.09% | 94.03% | 66.78% | 3,394 | 26,589 | 53,441 |
| | | Homopol_7-11bp | 84.96% | 90.78% | 79.83% | 551 | 1,370 | 5,422 | 67.33% | 86.61% | 55.06% | 578 | 3,052 | 3,740 |
| | | Homopol_gt11bp | 73.94% | 81.78% | 67.47% | 209 | 448 | 929 | 49.26% | 68.16% | 38.56% | 248 | 846 | 531 |
| | | Imp_Homopol_gt10bp | 81.47% | 88.47% | 75.50% | 392 | 972 | 2,996 | 62.30% | 81.15% | 50.55% | 466 | 1,962 | 2,006 |
| | | TR_lt51bp | 85.83% | 92.24% | 80.25% | 231 | 673 | 2,734 | 76.23% | 92.91% | 64.63% | 168 | 1,205 | 2,202 |
| | | TR51-220bp | 86.55% | 92.87% | 81.03% | 132 | 402 | 1,717 | 78.37% | 93.82% | 67.30% | 94 | 693 | 1,426 |
| | | TR201-10kbp | 85.79% | 90.99% | 81.16% | 64 | 150 | 646 | 75.46% | 89.77% | 65.08% | 59 | 278 | 518 |
| | | TR_gt100bp | 85.73% | 91.56% | 80.59% | 113 | 295 | 1,225 | 76.22% | 90.97% | 65.59% | 99 | 523 | 997 |
| | | TR_and_Homopol | 85.11% | 91.34% | 79.68% | 1,162 | 3,118 | 12,228 | 70.66% | 88.54% | 58.78% | 1,168 | 6,325 | 9,021 |
| | Segmental duplications | ChainSelf | 86.40% | 91.29% | 82.01% | 1,700 | 3,906 | 17,806 | 75.99% | 89.57% | 65.99% | 1,669 | 7,385 | 14,327 |
| | | ChainSelf_gt10kb | 81.37% | 85.05% | 77.99% | 922 | 1,480 | 5,243 | 68.62% | 87.02% | 56.64% | 568 | 2,915 | 3,808 |
| | | SegDups | 84.16% | 88.80% | 79.99% | 1,452 | 2,878 | 11,503 | 72.89% | 90.57% | 60.99% | 913 | 5,610 | 8,771 |
| | | SegDups_gt10kb | 82.88% | 87.11% | 79.04% | 1,341 | 2,401 | 9,055 | 70.12% | 89.03% | 57.83% | 816 | 4,831 | 6,625 |
| | Low mappability | LowMAD | 83.41% | 87.28% | 79.88% | 1,479 | 2,555 | 10,141 | 69.34% | 88.62% | 56.95% | 928 | 5,466 | 7,230 |
| | Other difficult regions | L1H | 87.47% | 96.22% | 80.18% | 7 | 44 | 178 | 52.98% | 99.92% | 36.04% | 0 | 142 | 80 |
| | | MHC | 75.91% | 90.25% | 65.49% | 370 | 1,804 | 3,424 | 52.31% | 92.92% | 36.40% | 145 | 3,325 | 1,903 |
| | Functional regions | CDS | 92.13% | 97.50% | 87.33% | 264 | 1,491 | 10,277 | 93.00% | 98.23% | 88.29% | 187 | 1,378 | 10,390 |

(b) Performance on ONT

| Dataset | Stratification type | Stratification subtype | Clair3-RNA SNP performance | | | | | | LongcallR SNP performance | | | | | |
|---|---|---|---|---|---|---|---|---|---|---|---|---|---|---|
| | | | Precision | Recall | F1-score | FP | FN | TP | Precision | Recall | F1-score | FP | FN | TP |
| ONT dRNA004 HG004 | Low complexity | Homopol_4-6bp | 86.84% | 92.27% | 82.01% | 1,809 | 4,735 | 21,584 | 58.99% | 94.41% | 42.89% | 668 | 15,030 | 11,288 |
| | | Homopol_7-11bp | 74.52% | 89.55% | 63.81% | 144 | 700 | 1,234 | 50.57% | 93.19% | 34.69% | 49 | 1,263 | 671 |
| | | Homopol_gt11bp | 61.80% | 82.13% | 49.54% | 47 | 219 | 215 | 29.86% | 68.03% | 19.12% | 39 | 351 | 83 |
| | | Imp_Homopol_gt10bp | 70.59% | 86.36% | 59.69% | 114 | 487 | 721 | 40.28% | 83.99% | 26.49% | 61 | 888 | 320 |
| | | TR_lt51bp | 83.23% | 88.63% | 78.44% | 119 | 255 | 928 | 52.94% | 91.30% | 37.28% | 42 | 742 | 441 |
| | | TR51-220bp | 85.16% | 91.50% | 79.64% | 65 | 179 | 700 | 49.26% | 88.50% | 34.13% | 39 | 579 | 300 |
| | | TR201-10kbp | 87.92% | 93.05% | 83.33% | 23 | 62 | 310 | 60.22% | 97.59% | 43.55% | 4 | 210 | 162 |
| | | TR_gt100bp | 86.85% | 92.99% | 81.48% | 43 | 130 | 572 | 54.10% | 96.35% | 37.61% | 10 | 438 | 264 |
| | | TR_and_Homopol | 79.61% | 90.28% | 71.20% | 387 | 1,454 | 3,594 | 50.50% | 91.02% | 34.94% | 174 | 3,284 | 1,764 |
| | Segmental duplications | ChainSelf | 83.19% | 84.48% | 81.94% | 1,771 | 2,125 | 9,643 | 62.16% | 92.63% | 46.77% | 438 | 6,264 | 5,504 |
| | | ChainSelf_gt10kb | 82.08% | 87.40% | 77.37% | 349 | 708 | 2,421 | 53.98% | 92.20% | 38.16% | 101 | 1,935 | 1,194 |
| | | SegDups | 79.89% | 80.94% | 78.88% | 1,263 | 1,436 | 5,362 | 59.36% | 94.18% | 43.34% | 182 | 3,852 | 2,946 |
| | | SegDups_gt10kb | 80.90% | 85.60% | 76.69% | 638 | 1,153 | 3,793 | 53.63% | 93.06% | 37.67% | 139 | 3,083 | 1,863 |
| | Low mappability | LowMAD | 76.61% | 75.96% | 77.27% | 1,093 | 1,016 | 3,453 | 49.88% | 90.12% | 34.48% | 169 | 2,928 | 1,541 |
| | Other difficult regions | L1H | 86.57% | 87.88% | 85.29% | 4 | 5 | 29 | 64.00% | 99.92% | 47.06% | 0 | 18 | 16 |
| | | MHC | 75.90% | 85.89% | 67.99% | 411 | 1,178 | 2,502 | 34.40% | 96.01% | 20.95% | 32 | 2,909 | 771 |
| | Functional regions | CDS | 94.59% | 95.95% | 93.26% | 434 | 742 | 10,272 | 82.51% | 98.88% | 70.79% | 88 | 3,217 | 7,797 |

**Supplementary Table 7. Annotation results of variant and editing sites across various functional regions.**

| Region | PacBio variant | PacBio editing | ONT variant | ONT editing |
|---|---|---|---|---|
| Intron | 160,450 | 2,915 | 75,496 | 1,805 |
| 3'UTR | 18,805 | 804 | 25,920 | 525 |
| 3'Flank | 7,797 | 205 | 12,943 | 516 |
| 5'Flank | 5,897 | 58 | 9,742 | 255 |
| Targeted_Region | 405 | 327 | 518 | 180 |
| RNA | 1,951 | 37 | 3,716 | 133 |
| Splice_Site | 41 | 847 | 155 | 107 |
| Splice_Region | 360 | 822 | 620 | 102 |
| IGR | 1,838 | 48 | 1,664 | 73 |
| Missense_Mutation | 4,134 | 12 | 5,962 | 53 |
| Silent | 5,090 | 6 | 6,774 | 41 |
| 5'UTR | 1,922 | 5 | 2,754 | 10 |
| Others | 170 | 0 | 371 | 0 |
| Total | 208,860 | 6,086 | 146,635 | 3,800 |

**Supplementary Table 8. Performance w/o phasing information.**

(a) Performance on SNP

| Read coverage | Allele depth | Disregarding zygosity | Platform | Dataset | Sample | SNP performance (apply phasing √) | | | | | | | SNP performance (apply phasing ✗) | | | | | | |
|---|---|---|---|---|---|---|---|---|---|---|---|---|---|---|---|---|---|---|---|
| | | | | | | TRUTH.FP | TRUTH.FN | TRUTH.TP | QUERY.TP | Precision | Recall | F1-score | TRUTH.FP | TRUTH.FN | TRUTH.TP | QUERY.TP | Precision | Recall | F1-score |
| DP≥4 | AD≥2 | × | PacBio | Iso-Seq(BCM) | HG004 | 5,085 | 8,547 | 154,478 | 154,493 | 96.81% | 94.76% | 95.77% | 6,100 | 10,214 | 152,810 | 152,818 | 96.16% | 93.73% | 94.93% |
| | | | | Iso-Seq(BCM) | HG005 | 6,714 | 9,935 | 227,229 | 227,251 | 97.13% | 95.81% | 96.47% | 8,175 | 12,972 | 224,196 | 224,215 | 96.48% | 94.53% | 95.50% |
| | | | | Iso-Seq(Google) | HG004 | 4,649 | 9,733 | 105,243 | 105,252 | 95.77% | 91.53% | 93.60% | 6,478 | 11,853 | 103,120 | 103,124 | 94.09% | 89.69% | 91.84% |
| | | | | Iso-Seq(Google) | HG005 | 4,822 | 9,658 | 112,116 | 112,123 | 95.88% | 92.07% | 93.93% | 6,672 | 12,068 | 109,705 | 109,712 | 94.27% | 90.09% | 92.13% |
| | | | | MAS-Seq | HG004 | 12,283 | 19,717 | 280,873 | 280,895 | 95.81% | 93.44% | 94.61% | 13,806 | 25,222 | 275,369 | 275,387 | 95.23% | 91.61% | 93.38% |
| | | | ONT | ONT dRNA004 | HG004 | 5,834 | 7,630 | 92,096 | 92,098 | 94.04% | 92.35% | 93.19% | 6,887 | 10,711 | 89,017 | 89,018 | 92.82% | 89.26% | 91.00% |
| | | | | ONT dRNA004 | HG005 | 6,297 | 7,863 | 102,930 | 102,929 | 94.23% | 92.90% | 93.56% | 7,349 | 10,701 | 100,093 | 100,093 | 93.16% | 90.34% | 91.73% |
| DP≥8 | AD≥2 | × | PacBio | Iso-Seq(BCM) | HG004 | 1,144 | 3,404 | 87,568 | 87,579 | 98.71% | 96.26% | 97.47% | 1,601 | 4,273 | 86,697 | 86,702 | 98.19% | 95.30% | 96.72% |
| | | | | Iso-Seq(BCM) | HG005 | 1,713 | 3,994 | 140,884 | 140,901 | 98.80% | 97.24% | 98.01% | 2,521 | 6,028 | 138,855 | 138,872 | 98.22% | 95.84% | 97.01% |
| | | | | Iso-Seq(Google) | HG004 | 1,254 | 4,056 | 62,321 | 62,328 | 98.03% | 93.89% | 95.91% | 2,017 | 5,374 | 61,008 | 61,010 | 96.80% | 91.90% | 94.29% |
| | | | | Iso-Seq(Google) | HG005 | 1,391 | 3,944 | 67,949 | 67,956 | 97.99% | 94.51% | 96.22% | 2,217 | 5,832 | 66,059 | 66,066 | 96.75% | 91.89% | 94.26% |
| | | | | MAS-Seq | HG004 | 3,673 | 8,138 | 188,532 | 188,548 | 98.09% | 95.86% | 96.96% | 4,489 | 12,017 | 184,661 | 184,675 | 97.63% | 93.89% | 95.72% |
| | | | ONT | ONT dRNA004 | HG004 | 3,027 | 3,570 | 67,724 | 67,726 | 95.72% | 94.99% | 95.36% | 3,894 | 5,356 | 65,944 | 65,945 | 94.42% | 92.49% | 93.45% |
| | | | | ONT dRNA004 | HG005 | 3,434 | 3,333 | 74,264 | 74,263 | 95.58% | 95.70% | 95.64% | 4,114 | 5,127 | 72,474 | 72,474 | 94.63% | 93.39% | 94.01% |
| DP≥10 | AD≥2 | × | PacBio | Iso-Seq(BCM) | HG004 | 784 | 2,728 | 70,468 | 70,478 | 98.90% | 96.27% | 97.57% | 1,073 | 3,390 | 69,805 | 69,809 | 98.49% | 95.37% | 96.90% |
| | | | | Iso-Seq(BCM) | HG005 | 1,121 | 3,103 | 116,329 | 116,345 | 99.05% | 97.40% | 98.22% | 1,657 | 4,711 | 114,725 | 114,740 | 98.58% | 96.06% | 97.30% |
| | | | | Iso-Seq(Google) | HG004 | 878 | 3,254 | 52,172 | 52,179 | 98.35% | 94.13% | 96.19% | 1,399 | 4,246 | 51,187 | 51,188 | 97.34% | 92.34% | 94.77% |
| | | | | Iso-Seq(Google) | HG005 | 919 | 3,053 | 56,567 | 56,574 | 98.40% | 94.88% | 96.61% | 1,475 | 4,612 | 55,002 | 55,009 | 97.39% | 92.26% | 94.76% |
| | | | | MAS-Seq | HG004 | 2,653 | 6,364 | 160,337 | 160,353 | 98.37% | 96.18% | 97.27% | 3,229 | 9,600 | 157,111 | 157,124 | 97.99% | 94.24% | 96.08% |
| | | | ONT | ONT dRNA004 | HG004 | 2,506 | 2,928 | 60,575 | 60,577 | 96.03% | 95.39% | 95.71% | 3,284 | 4,271 | 59,239 | 59,240 | 94.75% | 93.28% | 94.01% |
| | | | | ONT dRNA004 | HG005 | 2,904 | 2,676 | 66,238 | 66,237 | 95.80% | 96.12% | 95.96% | 3,527 | 4,000 | 64,920 | 64,920 | 94.85% | 94.20% | 94.52% |
| DP≥10 | AD≥4 | × | PacBio | Iso-Seq(BCM) | HG004 | 705 | 2,325 | 68,698 | 68,708 | 98.98% | 96.73% | 97.84% | 891 | 2,783 | 68,241 | 68,245 | 98.71% | 96.08% | 97.38% |
| | | | | Iso-Seq(BCM) | HG005 | 957 | 2,628 | 113,829 | 113,845 | 99.17% | 97.74% | 98.45% | 1,241 | 3,830 | 112,630 | 112,643 | 98.91% | 96.71% | 97.80% |
| | | | | Iso-Seq(Google) | HG004 | 816 | 2,737 | 50,972 | 50,976 | 98.42% | 94.90% | 96.63% | 1,068 | 3,475 | 50,242 | 50,242 | 97.92% | 93.53% | 95.67% |
| | | | | Iso-Seq(Google) | HG005 | 829 | 2,492 | 55,471 | 55,478 | 98.53% | 95.70% | 97.09% | 1,043 | 3,788 | 54,170 | 54,177 | 98.11% | 93.46% | 95.73% |
| | | | | MAS-Seq | HG004 | 2,339 | 5,171 | 156,795 | 156,810 | 98.53% | 96.81% | 97.66% | 2,682 | 7,616 | 154,359 | 154,371 | 98.29% | 95.30% | 96.77% |
| | | | ONT | ONT dRNA004 | HG004 | 2,358 | 2,462 | 59,619 | 59,621 | 96.20% | 96.03% | 96.11% | 3,099 | 3,410 | 58,680 | 58,681 | 94.98% | 94.51% | 94.75% |
| | | | | ONT dRNA004 | HG005 | 2,761 | 2,216 | 65,301 | 65,301 | 95.94% | 96.72% | 96.33% | 3,340 | 3,188 | 64,338 | 64,338 | 95.06% | 95.28% | 95.17% |
| DP≥4 | AD≥2 | √ | PacBio | Iso-Seq(BCM) | HG004 | 1,278 | 4,743 | 158,282 | 158,300 | 99.20% | 97.09% | 98.13% | 1,766 | 5,883 | 157,141 | 157,152 | 98.89% | 96.39% | 97.62% |
| | | | | Iso-Seq(BCM) | HG005 | 2,641 | 5,869 | 231,295 | 231,324 | 98.87% | 97.53% | 98.19% | 3,517 | 8,321 | 228,847 | 228,873 | 98.49% | 96.49% | 97.48% |
| | | | | Iso-Seq(Google) | HG004 | 1,289 | 6,375 | 108,601 | 108,612 | 98.83% | 94.46% | 96.59% | 2,397 | 7,774 | 107,199 | 107,205 | 97.81% | 93.24% | 95.47% |
| | | | | Iso-Seq(Google) | HG005 | 1,814 | 6,653 | 115,121 | 115,131 | 98.45% | 94.54% | 96.45% | 3,032 | 8,429 | 113,344 | 113,352 | 97.39% | 93.08% | 95.19% |
| | | | | MAS-Seq | HG004 | 3,923 | 11,368 | 289,222 | 289,255 | 98.66% | 96.22% | 97.42% | 4,878 | 16,304 | 284,287 | 284,315 | 98.31% | 94.58% | 96.41% |
| | | | ONT | ONT dRNA004 | HG004 | 3,788 | 5,585 | 94,141 | 94,144 | 96.13% | 94.40% | 95.26% | 4,188 | 8,013 | 91,715 | 91,717 | 95.63% | 91.97% | 93.76% |
| | | | | ONT dRNA004 | HG005 | 4,512 | 6,082 | 104,711 | 104,714 | 95.87% | 94.51% | 95.19% | 5,046 | 8,401 | 102,393 | 102,396 | 95.30% | 92.42% | 93.84% |
| DP≥8 | AD≥2 | √ | PacBio | Iso-Seq(BCM) | HG004 | 453 | 2,715 | 88,257 | 88,270 | 99.49% | 97.02% | 98.24% | 764 | 3,438 | 87,532 | 87,539 | 99.13% | 96.22% | 97.66% |
| | | | | Iso-Seq(BCM) | HG005 | 1,002 | 3,289 | 141,589 | 141,612 | 99.30% | 97.73% | 98.51% | 1,611 | 5,124 | 139,759 | 139,782 | 98.86% | 96.46% | 97.65% |
| | | | | Iso-Seq(Google) | HG004 | 541 | 3,343 | 63,034 | 63,041 | 99.15% | 94.96% | 97.01% | 1,030 | 4,387 | 61,995 | 61,997 | 98.37% | 93.39% | 95.81% |
| | | | | Iso-Seq(Google) | HG005 | 786 | 3,342 | 68,551 | 68,561 | 98.87% | 95.35% | 97.08% | 1,388 | 5,004 | 66,887 | 66,895 | 97.97% | 93.04% | 95.44% |
| | | | | MAS-Seq | HG004 | 1,545 | 6,018 | 190,652 | 190,676 | 99.20% | 96.94% | 98.06% | 2,023 | 9,559 | 187,119 | 187,141 | 98.93% | 95.14% | 97.00% |
| | | | ONT | ONT dRNA004 | HG004 | 2,156 | 2,700 | 68,594 | 68,597 | 96.95% | 96.21% | 96.58% | 2,665 | 4,128 | 67,172 | 67,174 | 96.18% | 94.21% | 95.19% |
| | | | | ONT dRNA004 | HG005 | 2,761 | 2,664 | 74,933 | 74,936 | 96.45% | 96.57% | 96.51% | 3,233 | 4,249 | 73,352 | 73,355 | 95.78% | 94.52% | 95.15% |
| DP≥10 | AD≥2 | √ | PacBio | Iso-Seq(BCM) | HG004 | 331 | 2,277 | 70,919 | 70,931 | 99.54% | 96.89% | 98.19% | 517 | 2,836 | 70,359 | 70,365 | 99.27% | 96.13% | 97.67% |
| | | | | Iso-Seq(BCM) | HG005 | 677 | 2,665 | 116,767 | 116,789 | 99.42% | 97.77% | 98.59% | 1,082 | 4,142 | 115,294 | 115,315 | 99.07% | 96.53% | 97.78% |
| | | | | Iso-Seq(Google) | HG004 | 399 | 2,775 | 52,651 | 52,658 | 99.25% | 94.99% | 97.07% | 720 | 3,567 | 51,866 | 51,867 | 98.63% | 93.57% | 96.03% |
| | | | | Iso-Seq(Google) | HG005 | 525 | 2,662 | 56,958 | 56,968 | 99.09% | 95.54% | 97.28% | 914 | 4,052 | 55,562 | 55,570 | 98.38% | 93.20% | 95.72% |
| | | | | MAS-Seq | HG004 | 1,149 | 4,868 | 161,833 | 161,857 | 99.30% | 97.08% | 98.18% | 1,410 | 7,789 | 158,922 | 158,943 | 99.12% | 95.33% | 97.19% |
| | | | ONT | ONT dRNA004 | HG004 | 1,807 | 2,230 | 61,273 | 61,276 | 97.14% | 96.49% | 96.81% | 2,269 | 3,257 | 60,253 | 60,255 | 96.37% | 94.87% | 95.62% |
| | | | | ONT dRNA004 | HG005 | 2,392 | 2,167 | 66,747 | 66,749 | 96.54% | 96.86% | 96.70% | 2,824 | 3,300 | 65,620 | 65,623 | 95.87% | 95.21% | 95.54% |
| DP≥10 | AD≥4 | √ | PacBio | Iso-Seq(BCM) | HG004 | 255 | 1,876 | 69,147 | 69,158 | 99.63% | 97.36% | 98.48% | 341 | 2,235 | 68,789 | 68,795 | 99.51% | 96.85% | 98.16% |
| | | | | Iso-Seq(BCM) | HG005 | 515 | 2,190 | 114,267 | 114,287 | 99.55% | 98.12% | 98.83% | 670 | 3,263 | 113,197 | 113,214 | 99.41% | 97.20% | 98.29% |
| | | | | Iso-Seq(Google) | HG004 | 339 | 2,260 | 51,449 | 51,453 | 99.35% | 95.79% | 97.54% | 393 | 2,800 | 50,917 | 50,917 | 99.23% | 94.79% | 96.96% |
| | | | | Iso-Seq(Google) | HG005 | 438 | 2,104 | 55,859 | 55,869 | 99.22% | 96.37% | 97.78% | 489 | 3,235 | 54,723 | 54,731 | 99.11% | 94.42% | 96.71% |
| | | | | MAS-Seq | HG004 | 840 | 3,678 | 158,288 | 158,309 | 99.47% | 97.73% | 98.59% | 873 | 5,813 | 156,162 | 156,180 | 99.44% | 96.41% | 97.90% |
| | | | ONT | ONT dRNA004 | HG004 | 1,660 | 1,765 | 60,316 | 60,319 | 97.32% | 97.16% | 97.24% | 2,087 | 2,399 | 59,691 | 59,693 | 96.62% | 96.14% | 96.38% |
| | | | | ONT dRNA004 | HG005 | 2,251 | 1,709 | 65,808 | 65,811 | 96.69% | 97.47% | 97.08% | 2,640 | 2,490 | 65,036 | 65,038 | 96.10% | 96.31% | 96.21% |

(a) Performance on Indel

| Read coverage | Allele depth | Disregarding zygosity | Platform | Dataset | Sample | Indel performance (apply phasing √) | | | | | | | Indel performance (apply phasing ✗) | | | | | | |
|---|---|---|---|---|---|---|---|---|---|---|---|---|---|---|---|---|---|---|---|
| | | | | | | TRUTH.FP | TRUTH.FN | TRUTH.TP | QUERY.TP | Precision | Recall | F1-score | TRUTH.FP | TRUTH.FN | TRUTH.TP | QUERY.TP | Precision | Recall | F1-score |
| DP≥4 | AD≥2 | × | PacBio | Iso-Seq(BCM) | HG004 | 3,696 | 5,892 | 18,677 | 18,694 | 83.49% | 76.02% | 79.58% | 4,135 | 7,338 | 17,296 | 17,335 | 80.74% | 70.21% | 75.11% |
| | | | | Iso-Seq(BCM) | HG005 | 3,441 | 5,374 | 26,212 | 26,233 | 88.40% | 82.99% | 85.61% | 4,190 | 7,069 | 24,619 | 24,655 | 85.47% | 77.69% | 81.40% |
| | | | | Iso-Seq(Google) | HG004 | 3,619 | 4,268 | 10,254 | 10,261 | 73.93% | 70.61% | 72.23% | 5,561 | 5,108 | 9,493 | 9,500 | 63.08% | 65.02% | 64.03% |
| | | | | Iso-Seq(Google) | HG005 | 3,358 | 3,216 | 9,952 | 9,968 | 74.80% | 75.58% | 75.19% | 5,741 | 3,901 | 9,319 | 9,327 | 61.90% | 70.49% | 65.92% |
| | | | | MAS-Seq | HG004 | 8,173 | 15,010 | 30,859 | 30,880 | 79.07% | 67.28% | 72.70% | 8,568 | 18,018 | 27,839 | 27,854 | 76.48% | 60.71% | 67.69% |
| | | | ONT | ONT dRNA004 | HG004 | 1,052 | 6,571 | 7,263 | 7,273 | 87.36% | 52.50% | 65.59% | 1,280 | 7,306 | 6,529 | 6,531 | 83.61% | 47.19% | 60.33% |
| | | | | ONT dRNA004 | HG005 | 1,029 | 5,581 | 7,946 | 7,957 | 88.55% | 58.74% | 70.63% | 1,220 | 6,254 | 7,271 | 7,273 | 85.64% | 53.76% | 66.05% |
| DP≥8 | AD≥2 | × | PacBio | Iso-Seq(BCM) | HG004 | 1,664 | 2,876 | 11,270 | 11,281 | 87.15% | 79.67% | 83.24% | 2,078 | 3,762 | 10,530 | 10,564 | 83.56% | 73.68% | 78.31% |
| | | | | Iso-Seq(BCM) | HG005 | 1,557 | 2,595 | 17,237 | 17,256 | 91.72% | 86.92% | 89.25% | 2,125 | 3,695 | 16,305 | 16,341 | 88.49% | 81.53% | 84.87% |
| | | | | Iso-Seq(Google) | HG004 | 1,504 | 2,319 | 6,499 | 6,505 | 81.22% | 73.70% | 77.28% | 2,933 | 2,887 | 6,043 | 6,050 | 67.35% | 67.67% | 67.51% |
| | | | | Iso-Seq(Google) | HG005 | 1,326 | 1,684 | 6,327 | 6,343 | 82.71% | 78.98% | 80.80% | 3,010 | 2,140 | 5,955 | 5,964 | 66.46% | 73.56% | 69.83% |
| | | | | MAS-Seq | HG004 | 4,841 | 10,170 | 22,233 | 22,250 | 82.13% | 68.61% | 74.77% | 5,583 | 12,370 | 20,330 | 20,342 | 78.46% | 62.17% | 69.37% |
| | | | ONT | ONT dRNA004 | HG004 | 781 | 5,341 | 5,850 | 5,860 | 88.24% | 52.27% | 65.65% | 1,028 | 5,889 | 5,404 | 5,406 | 84.02% | 47.85% | 60.98% |
| | | | | ONT dRNA004 | HG005 | 720 | 4,309 | 6,275 | 6,286 | 89.72% | 59.29% | 71.40% | 961 | 4,825 | 5,861 | 5,863 | 85.92% | 54.85% | 66.95% |
| DP≥10 | AD≥2 | × | PacBio | Iso-Seq(BCM) | HG004 | 1,213 | 2,258 | 9,092 | 9,102 | 88.24% | 80.11% | 83.98% | 1,587 | 2,934 | 8,563 | 8,597 | 84.42% | 74.48% | 79.14% |
| | | | | Iso-Seq(BCM) | HG005 | 1,154 | 2,056 | 14,404 | 14,421 | 92.59% | 87.51% | 89.98% | 1,648 | 2,912 | 13,714 | 13,752 | 89.30% | 82.49% | 85.76% |
| | | | | Iso-Seq(Google) | HG004 | 1,059 | 1,899 | 5,464 | 5,472 | 83.79% | 74.21% | 78.71% | 2,168 | 2,352 | 5,120 | 5,129 | 70.29% | 68.52% | 69.39% |
| | | | | Iso-Seq(Google) | HG005 | 923 | 1,356 | 5,313 | 5,328 | 85.23% | 79.67% | 82.36% | 2,261 | 1,735 | 5,012 | 5,021 | 68.95% | 74.28% | 71.52% |
| | | | | MAS-Seq | HG004 | 3,908 | 8,903 | 19,211 | 19,224 | 83.11% | 68.33% | 75.00% | 4,636 | 10,740 | 17,698 | 17,708 | 79.25% | 62.23% | 69.72% |
| | | | ONT | ONT dRNA004 | HG004 | 700 | 4,956 | 5,316 | 5,326 | 88.38% | 51.75% | 65.28% | 922 | 5,419 | 4,963 | 4,965 | 84.34% | 47.80% | 61.02% |
| | | | | ONT dRNA004 | HG005 | 641 | 3,927 | 5,669 | 5,679 | 89.86% | 59.08% | 71.29% | 864 | 4,367 | 5,338 | 5,340 | 86.07% | 55.00% | 67.12% |
| DP≥10 | AD≥4 | × | PacBio | Iso-Seq(BCM) | HG004 | 892 | 1,617 | 8,746 | 8,757 | 90.76% | 84.40% | 87.46% | 1,305 | 2,166 | 8,356 | 8,399 | 86.55% | 79.41% | 82.83% |
| | | | | Iso-Seq(BCM) | HG005 | 896 | 1,474 | 13,997 | 14,016 | 93.99% | 90.47% | 92.20% | 1,389 | 2,201 | 13,464 | 13,504 | 90.67% | 85.95% | 88.25% |
| | | | | Iso-Seq(Google) | HG004 | 746 | 1,376 | 5,252 | 5,262 | 87.58% | 79.24% | 83.20% | 1,512 | 1,773 | 4,965 | 4,976 | 76.70% | 73.69% | 75.16% |
| | | | | Iso-Seq(Google) | HG005 | 677 | 1,034 | 5,137 | 5,154 | 88.39% | 83.24% | 85.74% | 1,628 | 1,369 | 4,885 | 4,896 | 75.05% | 78.11% | 76.55% |
| | | | | MAS-Seq | HG004 | 3,164 | 7,289 | 18,645 | 18,659 | 85.50% | 71.89% | 78.11% | 3,933 | 8,870 | 17,372 | 17,383 | 81.55% | 66.20% | 73.08% |
| | | | ONT | ONT dRNA004 | HG004 | 649 | 4,384 | 5,255 | 5,265 | 89.03% | 54.52% | 67.62% | 880 | 4,820 | 4,931 | 4,933 | 84.86% | 50.57% | 63.37% |
| | | | | ONT dRNA004 | HG005 | 600 | 3,425 | 5,609 | 5,619 | 90.35% | 62.09% | 73.60% | 815 | 3,839 | 5,305 | 5,307 | 86.69% | 58.02% | 69.51% |
| DP≥4 | AD≥2 | √ | PacBio | Iso-Seq(BCM) | HG004 | 1,978 | 4,171 | 20,398 | 20,412 | 91.17% | 83.02% | 86.90% | 1,940 | 5,140 | 19,494 | 19,530 | 90.96% | 79.13% | 84.64% |
| | | | | Iso-Seq(BCM) | HG005 | 1,684 | 3,610 | 27,976 | 27,990 | 94.33% | 88.57% | 91.36% | 1,888 | 4,760 | 26,928 | 26,957 | 93.45% | 84.98% | 89.02% |
| | | | | Iso-Seq(Google) | HG004 | 2,604 | 3,251 | 11,271 | 11,276 | 81.24% | 77.61% | 79.38% | 4,140 | 3,685 | 10,916 | 10,921 | 72.51% | 74.76% | 73.62% |
| | | | | Iso-Seq(Google) | HG005 | 2,515 | 2,370 | 10,798 | 10,811 | 81.13% | 82.00% | 81.56% | 4,547 | 2,706 | 10,514 | 10,521 | 69.82% | 79.53% | 74.36% |
| | | | | MAS-Seq | HG004 | 4,926 | 11,752 | 34,117 | 34,127 | 87.39% | 74.38% | 80.36% | 4,612 | 14,052 | 31,805 | 31,810 | 87.34% | 69.36% | 77.32% |
| | | | ONT | ONT dRNA004 | HG004 | 745 | 6,263 | 7,571 | 7,580 | 91.05% | 54.73% | 68.36% | 829 | 6,854 | 6,981 | 6,982 | 89.39% | 50.46% | 64.50% |
| | | | | ONT dRNA004 | HG005 | 712 | 5,260 | 8,267 | 8,274 | 92.08% | 61.11% | 73.47% | 805 | 5,836 | 7,689 | 7,688 | 90.52% | 56.85% | 69.84% |
| DP≥8 | AD≥2 | √ | PacBio | Iso-Seq(BCM) | HG004 | 972 | 2,182 | 11,964 | 11,973 | 92.49% | 84.58% | 88.36% | 1,094 | 2,776 | 11,516 | 11,548 | 91.35% | 80.58% | 85.62% |
| | | | | Iso-Seq(BCM) | HG005 | 857 | 1,889 | 17,943 | 17,956 | 95.44% | 90.48% | 92.89% | 1,094 | 2,658 | 17,342 | 17,372 | 94.08% | 86.71% | 90.24% |
| | | | | Iso-Seq(Google) | HG004 | 1,072 | 1,887 | 6,931 | 6,937 | 86.62% | 78.60% | 82.41% | 2,236 | 2,190 | 6,740 | 6,747 | 75.11% | 75.48% | 75.29% |
| | | | | Iso-Seq(Google) | HG005 | 995 | 1,350 | 6,661 | 6,674 | 87.03% | 83.15% | 85.04% | 2,465 | 1,594 | 6,501 | 6,509 | 72.53% | 80.31% | 76.22% |
| | | | | MAS-Seq | HG004 | 3,117 | 8,438 | 23,965 | 23,974 | 88.49% | 73.96% | 80.58% | 3,286 | 10,065 | 22,635 | 22,639 | 87.33% | 69.22% | 77.23% |
| | | | ONT | ONT dRNA004 | HG004 | 591 | 5,150 | 6,041 | 6,050 | 91.10% | 53.98% | 67.79% | 734 | 5,594 | 5,699 | 5,700 | 88.59% | 50.46% | 64.30% |
| | | | | ONT dRNA004 | HG005 | 538 | 4,123 | 6,461 | 6,468 | 92.32% | 61.05% | 73.49% | 689 | 4,550 | 6,136 | 6,135 | 89.90% | 57.42% | 70.08% |
| DP≥10 | AD≥2 | √ | PacBio | Iso-Seq(BCM) | HG004 | 714 | 1,757 | 9,593 | 9,601 | 93.08% | 84.52% | 88.59% | 852 | 2,197 | 9,300 | 9,332 | 91.63% | 80.89% | 85.93% |
| | | | | Iso-Seq(BCM) | HG005 | 625 | 1,521 | 14,939 | 14,950 | 95.99% | 90.76% | 93.30% | 866 | 2,124 | 14,502 | 14,534 | 94.38% | 87.22% | 90.66% |
| | | | | Iso-Seq(Google) | HG004 | 733 | 1,573 | 5,790 | 5,798 | 88.78% | 78.64% | 83.40% | 1,634 | 1,818 | 5,654 | 5,663 | 77.61% | 75.67% | 76.63% |
| | | | | Iso-Seq(Google) | HG005 | 677 | 1,107 | 5,562 | 5,574 | 89.17% | 83.40% | 86.19% | 1,823 | 1,296 | 5,451 | 5,459 | 74.97% | 80.79% | 77.77% |
| | | | | MAS-Seq | HG004 | 2,545 | 7,532 | 20,582 | 20,587 | 89.00% | 73.21% | 80.34% | 2,781 | 8,877 | 19,561 | 19,563 | 87.55% | 68.78% | 77.04% |
| | | | ONT | ONT dRNA004 | HG004 | 537 | 4,792 | 5,480 | 5,489 | 91.09% | 53.35% | 67.29% | 673 | 5,169 | 5,213 | 5,214 | 88.57% | 50.21% | 64.09% |
| | | | | ONT dRNA004 | HG005 | 488 | 3,771 | 5,825 | 5,832 | 92.28% | 60.70% | 73.23% | 634 | 4,134 | 5,571 | 5,570 | 89.78% | 57.40% | 70.03% |
| DP≥10 | AD≥4 | √ | PacBio | Iso-Seq(BCM) | HG004 | 471 | 1,195 | 9,168 | 9,178 | 95.12% | 88.47% | 91.67% | 640 | 1,499 | 9,023 | 9,064 | 93.40% | 85.75% | 89.42% |
| | | | | Iso-Seq(BCM) | HG005 | 447 | 1,021 | 14,450 | 14,465 | 97.00% | 93.40% | 95.17% | 681 | 1,489 | 14,176 | 14,212 | 95.43% | 90.49% | 92.90% |
| | | | | Iso-Seq(Google) | HG004 | 464 | 1,094 | 5,534 | 5,544 | 92.28% | 83.49% | 87.67% | 1,038 | 1,299 | 5,439 | 5,450 | 84.00% | 80.72% | 82.33% |
| | | | | Iso-Seq(Google) | HG005 | 460 | 814 | 5,357 | 5,371 | 92.11% | 86.81% | 89.38% | 1,228 | 968 | 5,286 | 5,296 | 81.18% | 84.52% | 82.82% |
| | | | | MAS-Seq | HG004 | 1,952 | 6,071 | 19,863 | 19,871 | 91.06% | 76.59% | 83.20% | 2,277 | 7,208 | 19,034 | 19,039 | 89.32% | 72.53% | 80.05% |
| | | | ONT | ONT dRNA004 | HG004 | 495 | 4,229 | 5,410 | 5,419 | 91.63% | 56.13% | 69.61% | 637 | 4,576 | 5,175 | 5,176 | 89.04% | 53.07% | 66.50% |
| | | | | ONT dRNA004 | HG005 | 456 | 3,278 | 5,756 | 5,763 | 92.67% | 63.71% | 75.51% | 598 | 3,620 | 5,524 | 5,524 | 90.23% | 60.41% | 72.37% |

**Supplementary Table 9. Runtime and memory.**

| Testing dataset | Clair3 RNA | | Clair3 | | DeepVariant | | LongcallR | |
|---|---|---|---|---|---|---|---|---|
| | Runtime (min) | Peak memory (GB) | Runtime (min) | Peak memory (GB) | Runtime (min) | Peak memory (GB) | Runtime (min) | Peak memory (GB) |
| PacBio Iso-Seq | 18 | 6.3 | 23 | 4.5 | 41 | 20.0 | 8 | 13.3 |
| PacBio MAS-Seq | 51 | 7.5 | 54 | 58.7 | 49 | 32.8 | 206 | 82.5 |
| ONT cDNA | 34 | 7.2 | 38 | 49.1 | 41 | 21.8 | 64 | 58.2 |
| ONT dRNA004 | 29 | 7.3 | 52 | 84.6 | 60 | 45.6 | 49 | 33.6 |

### Supplementary methods

#### Description of RNA pileup input features

The pileup input includes 18 features as stated below:

A+/C+/G+/T+: The counts of all A/C/G/T nucleotides in the forward strand.

$I_S$+: The counts of insertions with the same starting positions as the candidate site in the forward strand.

$I_{1S}$+: The count of insertions with the highest read support in the forward strand.

$D_S$+: The counts of deletions with the same starting positions as the candidate site in the forward strand.

$D_{1S}$+: The counts of deletions with the highest read support in the forward strand.

$D_R$+: The counts of all non-starting (following) positions of deletions in the forward strand.

A-/C-/G-/T-: The counts of all A/C/G/T nucleotides in the reverse strand.

$I_S$-: The counts of insertions with the same starting positions as the candidate site in the reverse strand.

$I_{1S}$-: The counts of insertions with the highest read support in the reverse strand.

$D_S$-: The counts of deletions with the same starting positions as the candidate site in the reverse strand.

$D_{1S}$-: The counts of deletions with the highest read support in the reverse strand.

$D_R$-: The counts of all non-starting(following) positions of a deletion in the reverse strand.

#### Description of pileup network outputs

The output of both the pileup and full-alignment network has four tasks, including 1) the 21-genotype probabilistic model (21 probabilities); and 2) zygosity (3 probabilities); The details of the four tasks were given in Clair3's manuscript, and are given here again for clarity. The 21-genotype probabilistic model comprises all of the possible genotypes of a diploid sample at a genome position, including 'AA', 'AC', 'AG', 'AT', 'CC', 'CG', 'CT', 'GG', 'GT', 'TT', 'AI', 'CI', 'GI', 'TI', 'AD', 'CD', 'GD', 'TD', 'II', 'DD', and 'ID', where 'A', 'C', 'G', 'T', 'I' (insertion) and 'D' (deletion) denote the six possible alleles. The zygosity task outputs the probability of the input being 1) a homozygous reference (0/0); 2) heterozygous with 1 or 2 alternative alleles (0/1 or 1/2); or 3) a homozygous variant (1/1). The zygosity task is partially redundant to the 21-genotype task, but it makes decisions independently, and it crosschecks the decisions made by the 21-genotype task.

#### Command line used

#### Read alignment

#### Minimap2 (v2.17-r941)

```
# Align ONT reads using minimap2 splice mode to GRCh38 by default
minimap2 -t ${THREADS} -aL --splice -x map-ont ref.fa input.fastq.gz | samtools view -bh -o output.unsorted.bam -
samtools sort -@${THREADS} -o output.sorted.bam output.unsorted.bam && samtools index -@ ${THREADS} output.sorted.bam
```

#### Pbmm2 (1.13.1)

```
# Align PacBio Iso-Seq and MAS-Seq reads using pbmm2 to GRCh38
```

```
pbmm2 -t ${THREADS} --preset ref.fa input.fastq.gz | samtools view -bh -o output.unsorted.bam
-
```

**BAM subsampling**

**Samtools(v1.10)**

```
samtools view -@ ${THREADS} -s 0.${RATIO} -b -o subsampled.bam ${BAM}
samtools index -@ ${THREADS} subsampled.bam
```

**Coverage calculation**

**Mosdepth(v0.2.9)**

```
mosdepth -t ${THREADS} -n -x --quantize 0:15:150: output ${BAM}
```

**Running Clair3-RNA (v0.2.0)**

```
docker run -it \
  -v ${INPUT_DIR}:${INPUT_DIR} \
  -v ${OUTPUT_DIR}:${OUTPUT_DIR} \
  hkubal/clair3-rna:latest \
  /opt/bin/run_clair3_rna \
    --bam_fn ${INPUT_DIR}/sample.bam \
    --ref_fn ${INPUT_DIR}/ref.fa \
    --threads ${THREADS} \
    --platform ${PLATFORM} \
    --output_dir ${OUTPUT_DIR} \
    --enable_phasing_model #optional
```

**Running Clair3 (v 1.0.10)**

```
bash run_clair3.sh \
  -b {INPUT_DIR}/sample.bam
  -f ${INPUT_DIR}/ref.fa \
  -m ${MODEL_PATH} \
  -t ${THREAD} \
  -p ${PLATFORM} \
  -o ${OUTPUT_DIR}
```

**Running longcallR (v 0.1.0)**

```
longcallR \
  --bam-path {INPUT_DIR}/sample.bam \
  --ref-path ${INPUT_DIR}/ref.fa \
  --output ${OUTPUT_DIR}/output \
  --platform ${PLATFORM} \
```

```
  --preset ${PRESET} \
  -t ${THREAD}
```

**Running DeepVariant (v 1.6.1)**

```
docker run \
  -v ${INPUT_DIR}:${INPUT_DIR} \
  -v ${OUTPUT_DIR}:${OUTPUT_DIR} \
  google/deepvariant:"1.6.1" \
  /opt/deepvariant/bin/run_deepvariant \
    --model_type=${PLATFORM} \
    --ref ref.fa \
    --reads {INPUT_DIR}/sample.bam \
    --output_vcf ${OUTPUT_DIR}/output.vcf.gz \
    --num_shards ${THREAD}
```

**Benchmarking**

**hap.py (v0.3.12)**

```
hap.py ${GIAB_BASELINE_VCF} output.vcf.gz \
  -o ${OUTPUT_DIR}/happy \
  -r ${REF} \
  -f ${GIAB_CONFIDENT_BED} \
  --threads ${THREADS} \
  --pass-only \
  --engine=vcfeval
```

**qfy.py (v0.3.12)**

```
# Benchmarking all genome stratifications regions
qfy.py ${OUTPUT_DIR}/happy.vcf.gz \
  -t ga4gh \
  --stratification v3.0-GRCh38-stratifications.tsv \
  -o ${OUTPUT_PREFIX} \
  -r ${REF} \
  --threads ${THREADS}
```

**WhatsHap (v1.4)**

```
# Calculate the switch errors using WhatsHap
whatshap compare \
  --ignore-sample-name \
  --switch-error-bed ${OUTPUT_DIR}/switches.bed \
  --only-snvs \
  ${INPUT_DIR}/phased_variants.vcf.gz
```

### Data availability

#### Reference genomes

| Name | Format | URL(s) |
|---|---|---|
| GRCh38 | FASTA | https://ftp-trace.ncbi.nlm.nih.gov/ReferenceSamples/giab/release/references/GRCh38/GCA_000001405.15_GRCh38_no_alt_analysis_set_maskedGRC_exclusions_v2.fasta.gz |
| GRCh38 Stratification regions (v3.0) | BED | https://ftp-trace.ncbi.nlm.nih.gov/giab/ftp/release/genome-stratifications/v3.0/GRCh38 |

#### GIAB truth variants

| Name | Reference | Version | Format | URL(s) |
|---|---|---|---|---|
| GIAB HG002 | GRCh38 | v4.2.1 | VCF/BED | ftp://ftp-trace.ncbi.nlm.nih.gov/giab/ftp/release/AshkenazimTrio/HG002_NA24385_son/NISTv4.2.1/GRCh38/ |
| GIAB HG004 | GRCh38 | v4.2.1 | VCF/BED | ftp://ftp-trace.ncbi.nlm.nih.gov/giab/ftp/release/AshkenazimTrio/HG004_NA24143_mother/NISTv4.2.1/GRCh38/ |
| GIAB HG005 | GRCh38 | v4.2.1 | VCF/BED | https://ftp-trace.ncbi.nlm.nih.gov/giab/ftp/release/ChineseTrio/HG005_NA24631_son/NISTv4.2.1/GRCh38/ |

#### Pacific Bioscience (PacBio) sequencing data

| Name | Reference | Instruments/Sequ | Format | URL(s) |
|---|---|---|---|---|
| GIAB HG002 | GRCh38 | Iso-Seq/cDNA | BAM | https://ftp-trace.ncbi.nlm.nih.gov/ReferenceSamples/giab/data_RNAseq/AshkenazimTrio/HG002_NA24385_son/Baylor_PacBio/reads/m64139_220127_180020.hifi_reads.bam |
| GIAB HG002 | GRCh38 | Iso-Seq/cDNA | BAM | https://ftp-trace.ncbi.nlm.nih.gov/ReferenceSamples/giab/data_RNAseq/AshkenazimTrio/HG002_NA24385_son/Baylor_PacBio/reads/m64139_220130_061226.hifi_reads.bam |
| GIAB HG002 | GRCh38 | Iso-Seq/cDNA | BAM | https://ftp-trace.ncbi.nlm.nih.gov/ReferenceSamples/giab/data_RNAseq/AshkenazimTrio/HG002_NA24385_son/Baylor_PacBio/reads/m64139_220131_122551.hifi_reads.bam |
| GIAB HG004 | GRCh38 | Iso-Seq/cDNA | BAM | https://ftp-trace.ncbi.nlm.nih.gov/ReferenceSamples/giab/data_RNAseq/AshkenazimTrio/HG004_NA24143_mother/Baylor_PacBio/reads/m64139_220124_190646.hifi_reads.bam |
| GIAB HG005 | GRCh38 | Iso-Seq/cDNA | BAM | https://ftp-trace.ncbi.nlm.nih.gov/ReferenceSamples/giab/data_RNAseq/ChineseTrio/HG005_NA24631_son/Baylor_PacBio/reads/m64139_220129_000012.hifi_reads.bam |
| GIAB HG002 | GRCh38 | Iso-Seq/cDNA | BAM | https://ftp-trace.ncbi.nlm.nih.gov/ReferenceSamples/giab/data_RNAseq/AshkenazimTrio/HG002_NA24385_son/Google_PacBio/reads/GM26105.m64267e_220109_182119.subreads.bam |
| GIAB HG002 | GRCh38 | Iso-Seq/cDNA | BAM | https://ftp-trace.ncbi.nlm.nih.gov/ReferenceSamples/giab/data_RNAseq/AshkenazimTrio/HG002_NA24385_son/Google_PacBio/reads/GM27730.m64267e_220111_003241.subreads.bam |
| GIAB HG002 | GRCh38 | Iso-Seq/cDNA | BAM | https://ftp-trace.ncbi.nlm.nih.gov/ReferenceSamples/giab/data_RNAseq/AshkenazimTrio/HG002_NA24385_son/Google_PacBio/reads/HG002.m64284e_220109_013023.subreads.bam |
| GIAB HG004 | GRCh38 | Iso-Seq/cDNA | BAM | https://ftp-trace.ncbi.nlm.nih.gov/ReferenceSamples/giab/data_RNAseq/AshkenazimTrio/HG004_NA24143_mother/Google_PacBio/reads/HG004.m64284e_220110_074143.subreads.bam |
| GIAB HG005 | GRCh38 | Iso-Seq/cDNA | BAM | https://ftp-trace.ncbi.nlm.nih.gov/ReferenceSamples/giab/data_RNAseq/ChineseTrio/HG005_NA24631_son/Google_PacBio/reads/HG005.m64168e_220110_021232.subreads.bam |
| GIAB HG002 | GRCh38 | MAS-Seq/cDNA | BAM | https://ftp-trace.ncbi.nlm.nih.gov/ReferenceSamples/giab/data_RNAseq/AshkenazimTrio/HG002_NA24385_son/PacBio_Pacbio-MASseq/GM24385/2-FLNC/giab_na24385.hifi_reads.lima.0--0.lima.IsoSeqX_bc02_5p--IsoSeqX_3p.refined.bam |
| GIAB HG002 | GRCh38 | MAS-Seq/cDNA | BAM | https://ftp-trace.ncbi.nlm.nih.gov/ReferenceSamples/giab/data_RNAseq/AshkenazimTrio/HG002_NA24385_son/PacBio_Pacbio-MASseq/GM26105/2-FLNC/giab_na26105.hifi_reads.lima.0--0.lima.IsoSeqX_bc04_5p--IsoSeqX_3p.refined.bam |
| GIAB HG002 | GRCh38 | MAS-Seq/cDNA | BAM | https://ftp-trace.ncbi.nlm.nih.gov/ReferenceSamples/giab/data_RNAseq/AshkenazimTrio/HG002_NA24385_son/PacBio_Pacbio-MASseq/GM27730/2-FLNC/giab_na27730.hifi_reads.lima.0--0.lima.IsoSeqX_bc05_5p--IsoSeqX_3p.refined.bam |
| GIAB HG004 | GRCh38 | MAS-Seq/cDNA | BAM | https://ftp-trace.ncbi.nlm.nih.gov/ReferenceSamples/giab/data_RNAseq/AshkenazimTrio/HG004_NA24143_mother/PacBio_Pacbio-MASseq/2-FLNC/giab_na24143.hifi_reads.lima.0--0.lima.IsoSeqX_bc01_5p--IsoSeqX_3p.refined.bam |
| GIAB HG004 | GRCh38 | DNA | BAM | https://downloads.pacbcloud.com/public/revio/2022Q4/HG004-rep1/analysis/HG004.m84010_220919_232145_s1.GRCh38.bam |

### Oxford Nanopore (ONT) sequencing data

| Name | Reference | Sequencing type | Format | URL(s) |
|---|---|---|---|---|
| GIAB HG002 | GRCh38 | cDNA | BAM | https://s3.amazonaws.com/gtl-public-data/giab/bams/cDNA/05_09_23_R941_GIAB_cDNA_PCS111_NA24385_Guppy_6.4.6_sup.pass.fastq.gz.hg38.bam |
| GIAB HG002 | GRCh38 | cDNA | BAM | https://s3.amazonaws.com/gtl-public-data/giab/bams/cDNA/05_09_23_R941_GIAB_cDNA_PCS111_NA26105_Guppy_6.4.6_sup.pass.fastq.gz.hg38.bam |
| GIAB HG002 | GRCh38 | cDNA | BAM | https://s3.amazonaws.com/gtl-public-data/giab/bams/cDNA/05_09_23_R941_GIAB_cDNA_PCS111_NA27730_Guppy_6.4.6_sup.pass.fastq.gz.hg38.bam |
| GIAB HG004 | GRCh38 | cDNA | BAM | https://s3.amazonaws.com/gtl-public-data/giab/bams/cDNA/05_09_23_R941_GIAB_cDNA_PCS111_NA24143_Guppy_6.4.6_sup.pass.fastq.gz.hg38.bam |
| GIAB HG005 | GRCh38 | cDNA | BAM | https://s3.amazonaws.com/gtl-public-data/giab/bams/cDNA/05_09_23_R941_GIAB_cDNA_PCS111_NA24631_Guppy_6.4.6_sup.pass.fastq.gz.hg38.bam |
| GIAB HG002 | GRCh38 | dRNA002 | BAM | https://s3.amazonaws.com/gtl-public-data/giab/bams/dRNA/03_30_23_R941_DRS_NA24385_dRNA_Guppy_6.4.6_rna_hac_prom.pass.NoU.fastq.gz.hg38.bam |
| GIAB HG002 | GRCh38 | dRNA002 | BAM | https://s3.amazonaws.com/gtl-public-data/giab/bams/dRNA/03_30_23_R941_DRS_NA26105_dRNA_Guppy_6.4.6_rna_hac_prom.pass.NoU.fastq.gz.hg38.bam |
| GIAB HG002 | GRCh38 | dRNA002 | BAM | https://s3.amazonaws.com/gtl-public-data/giab/bams/dRNA/03_30_23_R941_DRS_NA27730_dRNA_Guppy_6.4.6_rna_hac_prom.pass.NoU.fastq.gz.hg38.bam |
| GIAB HG004 | GRCh38 | dRNA002 | BAM | https://s3.amazonaws.com/gtl-public-data/giab/bams/dRNA/03_30_23_R941_DRS_NA24143_dRNA_Guppy_6.4.6_rna_hac_prom.pass.NoU.fastq.gz.hg38.bam |
| GIAB HG005 | GRCh38 | dRNA002 | BAM | https://s3.amazonaws.com/gtl-public-data/giab/bams/dRNA/03_30_23_R941_DRS_NA24631_dRNA_Guppy_6.4.6_rna_hac_prom.pass.NoU.fastq.gz.hg38.bam |
| GIAB HG002 | GRCh38 | dRNA004 | FASTQ | https://www.ncbi.nlm.nih.gov/sra/SRX26304755 |
| GIAB HG004 | GRCh38 | dRNA004 | FASTQ | https://www.ncbi.nlm.nih.gov/sra/SRX26304756 |
| GIAB HG005 | GRCh38 | dRNA004 | FASTQ | https://www.ncbi.nlm.nih.gov/sra/SRX26304757 |
| GIAB HG004 | GRCh38 | DNA | BAM | https://labs.epi2me.io/giab-2023.05/ |
| GIAB HG005 | GRCh38 | DNA | FASTQ | https://s3-us-west-2.amazonaws.com/human-pangenomics/index.html?prefix=NHGRI_UCSC_panel/HG005/nanopore/Guppy_6.1.2/ |
